# MTBseq-nf: Enabling Scalable Tuberculosis Genomics “Big Data” Analysis through a User-Friendly Nextflow Wrapper for MTBseq pipeline

**DOI:** 10.1101/2025.04.17.649337

**Authors:** Abhinav Sharma, Davi Josué Marcon, Johannes Loubser, Karla Valéria Batista Lima, Gian van der Spuy, Emilyn Costa Conceição

**Affiliations:** SAMRC Centre for Tuberculosis Research; Division of Molecular Biology and Human Genetics, Faculty of Medicine and Health Sciences, Stellenbosch University, Cape Town, South Africa; Universidade do Estado do Pará, Instituto de Ciências Biológicas e da Saúde, Pós-Graduação em Biologia Parasitária na Amazônia, Belém, Pará, Brazil; Instituto Evandro Chagas, Seção de Bacteriologia e Micologia, Ananindeua, Pará, Brazil

**Keywords:** Data analysis pipeline, Genomic surveillance, MTBseq, Mycobacterium tuberculosis, Mycobacterium tuberculosis complex, Nextflow, Tuberculosis genomics, Whole- genome sequencing, Workflow

## Abstract

The MTBseq pipeline, published in 2018, was designed to address bioinformatics challenges in tuberculosis research using whole-genome sequencing data. It was the first publicly available pipeline on GitHub to perform full analysis of whole-genome sequencing (WGS) data for Mycobacterium tuberculosis encompassing quality control through mapping, variant calling for lineage classification, drug resistance prediction, and phylogenetic inference. However, the pipeline’s architecture is not optimal for analyses on high-performance computing or cloud computing environments, which often involve large datasets. To optimize the pipeline, we created MTBseq-nf, a Nextflow wrapper which offers shorter execution times through parallelization along with multiple other key improvements. The MTBseq-nf wrapper, as opposed to the linear, batched analysis of samples in the TBfull step of MTBseq pipeline, can execute multiple instances of the same step in parallel and therefore makes full use of the provided computational resources. For evaluation of scalability and reproducibility, we used 90 M. tuberculosis genomes (European Nucleotide Archive – ENA- accession PRJEB7727) for the benchmarking analysis on a dedicated computational server. In our experiments the execution time of MTBseq-nf parallel analysis mode is at least twice as fast as the standard MTBseq pipeline for more than 20 samples. Furthermore, the MTBseq-nf wrapper facilitates reproducibility using the nf-core, bioconda, and biocontainers projects for platform independence. The proposed MTBseq-nf wrapper pipeline is, user-friendly, optimized for hardware efficiency, scalable for larger datasets, and exhibits improved reproducibility.

## INTRODUCTION

Next-generation sequencing (NGS) has revolutionized tuberculosis (TB) research, diagnosis, and surveillance, however analyzing this increasingly voluminous amount of complex bioinformatics data, presents numerous computational challenges (Berger & Yu, 2022). The MTBseq pipeline (Kohl et al., 2018) was published in 2018 to address some of the challenges of data-analysis and improve the reproducibility of whole-genome sequencing (WGS) analysis of Mycobacterium *tuberculosis* complex (MTBC). MTBseq (MTBseq-standard) was one of the first comprehensive end-to-end pipelines and is widely used by researchers working with WGS data to generate and analyze the principal outputs namely (i) strain classification, (ii) phylogenetic tree, (iii) mapping and variant statistics, (iv) Single Nucleotide Polymorphism (SNP) distance matrix and (v) cluster groups. On the other hand, the computational infrastructure available to researchers ranges from the traditional in-house servers or high-performance computing (HPC) platforms. The available options have only been diversified in recent years with the development of (i) batch computing functionality by cloud computing vendors such as Oracle, Amazon Web Services (AWS), Google, Microsoft Azure, International Business Machine (IBM) and Alibaba etc (ii) open source job orchestrators like Kubernetes, Apache Meson and Hashicorp Nomad to analyze the exponentially increasing volume of data in an efficient manner (Stephens, 2015) . This adoption trend of NGS technologies and use of modern job orchestrators is expected to continue for foreseeable future, especially in the context of precision public health and precision medicine. The MTBseq pipeline, in it current form performs sub-optimally for these modern computing environments, therefore we aimed to enhance the MTBseq pipeline for traditional as well as modern job orchestrators utilized in large-scale genomic analysis, while optimizing the costs and making the analysis time predictable. We developed MTBseq-nf, a wrapper built with the Nextflow workflow engine (Di Tommaso et al., 2017) - to provide an alternative to current users of MTBseq-standard with various enhancements in its user-friendliness, maintainability, reproducibility and scalability, for large-scale genomic data analysis.

## MATERIALS AND METHODS

### The design of MTBseq (standard) pipeline

The MTBseq pipeline by Kohl et al. (2018) (MTBseq-standard) relies upon underlying tools such as `GATK3` (McKenna et al., 2010), `PICARD` (*Picard Toolkit*, 2019), `BWA` (Li & Durbin, 2009) and `SAMTOOLS` (Li et al., 2009); and glues these tools together using the `perl5` (Wall et al., 1994) programming language as shown in Figure 1.

**FIGURE 1.**
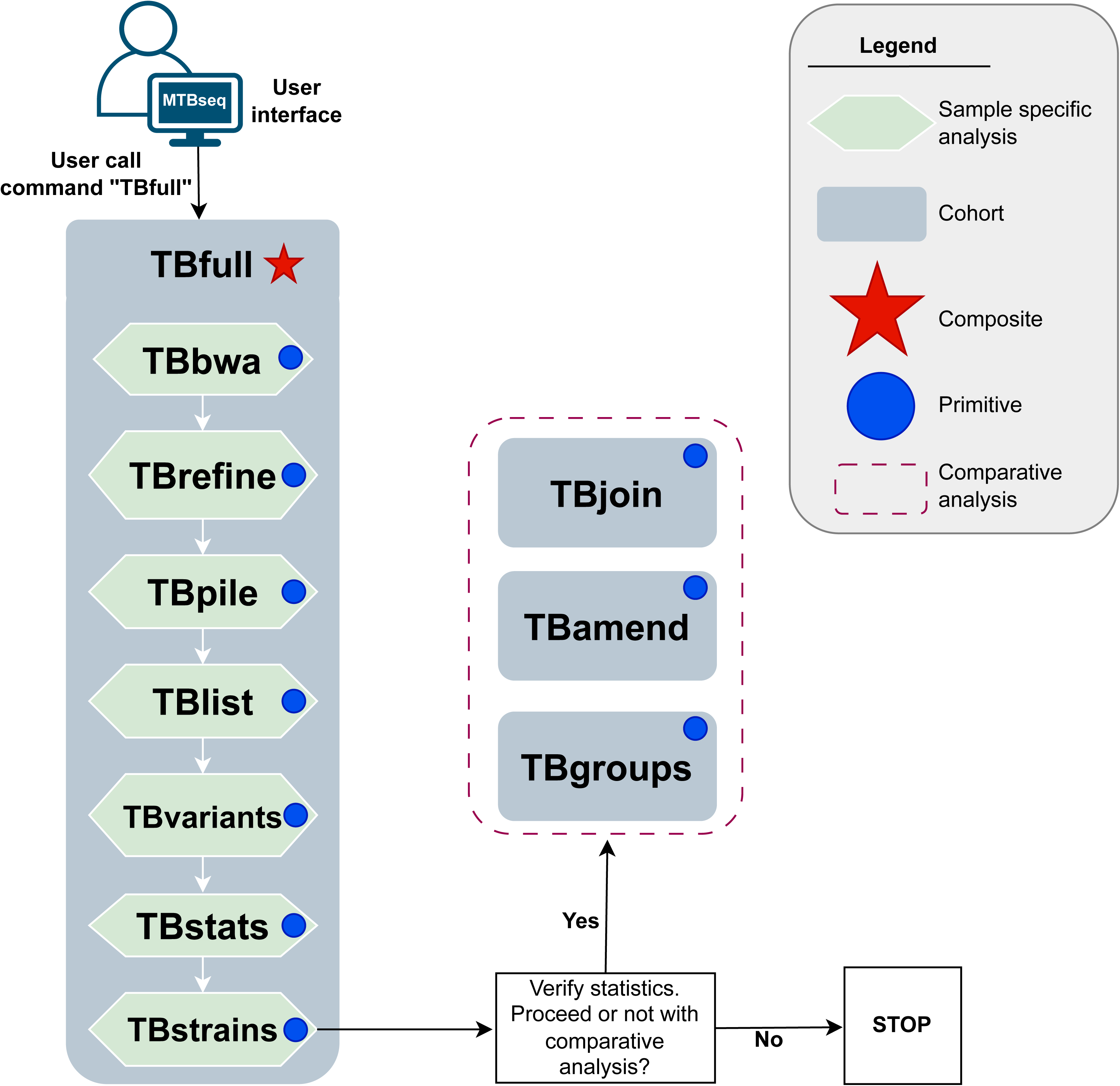
A schematic diagram of MTBseq-standard pipeline, published by Kohl et al (2018).

Its execution model is built upon core abstractions (called **steps**) such as reference mapping (`TBbwa`) and variant calling (`TBvariants`). These steps can be broadly classified into two groups based on (i) the quantity of samples analyzed per step, with sample-specific steps (`TBbwa`, `TBvariants`) and cohort-specific steps (`TBamend`, `TBgroups`) as well as (ii) composition, with primitive single-purpose steps (`TBbwa`, `TBgroups`), and composite multi-purpose step (`TBfull`), combining many of the primitive steps to facilitate routine usage of certain primitive steps, as summarized in Figure 1.

### Implementation of MTBseq-nf wrapper

We developed MTBseq-nf wrapper, on top of the open-source MTBseq pipeline, and implemented numerous enhancements across four broad themes (i) user-friendliness (ii) scalability (iii) reproducibility and (iv) maintainability, as summarized in Table 1.

**TABLE 1.** Summary of thematic enhancements in MTBseq-nf, Nextflow wrapper for the original MTBseq pipeline.

| Theme | Feature |
| --- | --- |
| User-friendliness | Ease of download |
| User-friendliness | Explicit samplesheet |
| User-friendliness | Graphical user interface |
| User-friendliness | MultiQC Summary report |
| User-friendliness | CSV and TSV format cleanup |
| User-friendliness | Remote monitoring |
| User-friendliness | Manual steps |
| User-friendliness | Flexible output location |
| Maintainability | Extensibility |
| Maintainability | Module testing |
| Maintainability | Test dataset |
| Scalability | Parallel execution |
| Scalability | HPC compatibility |
| Scalability | Resource allocation |
| Scalability | Dynamic retries |
| Scalability | Execution cache |
| Scalability | Reduced data footprint |
| Scalability | Reduced cloud computing costs |
| Reproducibility | Declarative parameters file |
| Reproducibility | Portability |
| Reproducibility | Save intermediate files |

In the implementation of MTBseq-nf wrapper pipeline we built upon two core components.

Firstly, the MTBseq-standard pipeline Kohl et al. (2018) exposes a `--step` parameter allowing users to dictate which step of the pipeline they wish to initiate. Figure 1 highlights, how the MTBseq-standard pipeline facilitates the usage of sequential sample-specific steps for routine executions, through a composite `TBfull` step which automates the execution of sample-specific primitive steps from `TBbwa` step until the `TBstrains` step. Upon completion of these steps, the users are expected to provide a TSV file, containing a list of samples and corresponding library names, that should be included in further comparative analysis steps such as `TBjoin`, `TBamed` and `TBgroup` generating the principal outputs of the pipeline as described in Supplemental data.

Secondly, the use of Nextflow workflow manager as per nf-core best-practices pipeline template. Nextflow is a workflow management system designed to address the challenges of high-throughput NGS data at scale in a reproducible manner (Di Tommaso et al., 2017), allowing researchers to create complex workflows that integrate multiple bioinformatics tools into a single, cohesive workflow while maintaining portability across different computing infrastructure. The nf-core community further enhances the Nextflow ecosystem by bringing together Nextflow users through a Slack group, hackathons, seminars, training, and other community initiatives that foster collaboration and knowledge sharing (P. A. Ewels et al., 2020; Langer et al., 2024). Moreover, the nf-core pipeline template is constructed with rigorous standards to guarantee robustness, portability, and user-friendliness, due to its integration with other projects from the nf-core ecosystem, such as `nf-core/configs` and `nf-core/modules`.

Internally, the `MTBseq-standard` pipeline depends on the sequential analysis of input sequences thorough the `foreach` looping structures, with each sample advancing to the subsequent step only after all FASTQ samples have completed an individual step (Figure 2a). On the other hand, the `MTBseq-nf` wrapper pipeline has two execution modes: (i) default and (ii) parallel mode, activated by the `--parallel` parameter. Both of these modes are functionally equivalent to the combination of `TBfull`, `TBgroups`, `TBamend` and `TBjoin` steps of the MTBseq pipeline, The principal insights we employed to implement the `--parallel` mode was to combine the advantages of both (i) MTBseq’s modular step-wise architecture exposed through the `--step` parameter to facilitate the independent execution of individual steps in the MTBseq pipeline, as well as (ii) the inherent task parallelization provided by the Nextflow workflow engine.

**FIGURE 2.**
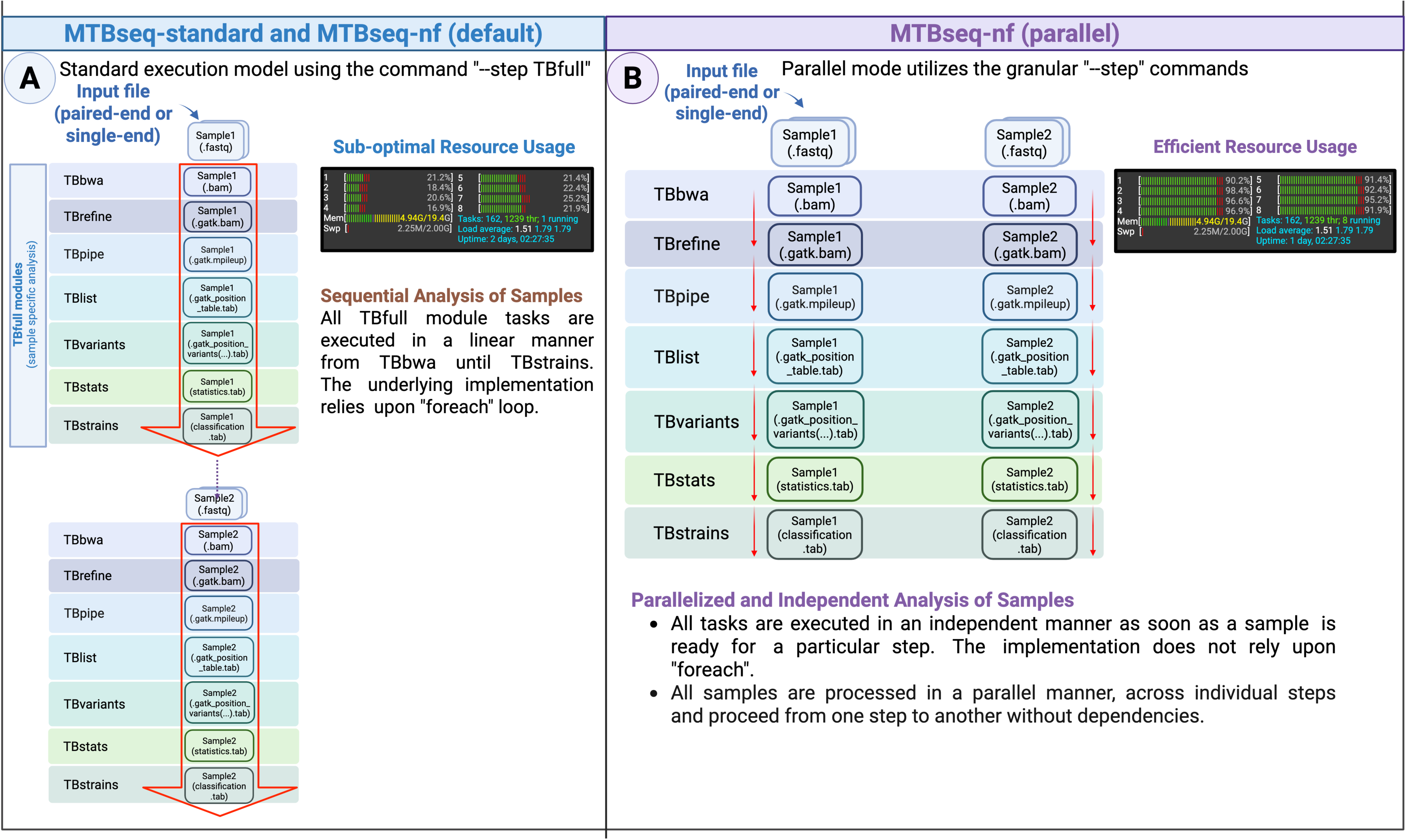
An overview of the MTBseq-standard, MTBseq-nf (default) and the MTBseq-nf `parallel` modes, which allows an individual sample to continue to the next steps, independently from other samples.

Consequently, when executed with the `--parallel` parameter, the MTBseq-nf wrapper pipeline employs the exact same primitive steps utilized by the `TBfull` step and optimizes the movement of intermediate files through the individual steps (Figure 2b); resulting in a significant reduction in the overall execution time of the pipeline, especially when the number of samples in a cohort are large. We depend on the fact that when a sample-specific step is executed with a single sample, the `foreach` loop iterates only once, since only a single sample is accessible to that particular step. Moreover, the MTBseq-nf pipeline (in both modes) automates the generation of a TSV file when all samples undergo the initial phases of the `TBfull` step and initiates the comparative analysis stage, mirroring the behavior of the MTBseq-standard pipeline for subsequent steps, minimizing the manual intervention by users.

### Validation Infrastructure and Dataset

For the computational studies, we selected a virtual server machine on the Oracle Cloud Infrastructure, featuring 32 Central Processing Unit (CPU) (16 Oracle CPUs), 64 GB of RAM, and a boot disk capacity of 2 TB. The essential software requirements for the experiments were installations of (i) Java programming language (Arnold et al., 2005), (ii) Nextflow (Di Tommaso et al., 2017), and (iii) Docker (Merkel, 2014). The MTBseq-standard pipeline v1.1.0 was installed using the bioconda recipe file provided as part of Supplemental data. The decision to utilize a server was crucial for our analysis of execution runtime, considering HPC queue systems impose an unpredictable delay before a submitted job starts execution. The Docker platform was chosen for its extensive utilization in cloud batch computing environments (AWS Batch, Azure Batch and Google Batch) as well as contemporary container orchestration solutions like Kubernetes and Hashicorp Nomad. The selection of Docker containers was influenced by the upstream biocontainer project assets (Veiga Leprevost et al., 2017), which are derived from bioconda recipes (Gruning et al., 2018). This approach enabled us to conduct the risk of non-reproducibility stemming from infrastructural discrepancies and enabled us to conduct an isolated analysis of the primary results of the pipeline.

We relied upon the original dataset utilized by the original study Kohl et al. (2018) to evaluate the reproducibility and scalability of the MTBseq-nf wrapper in terms of (i) growth of execution time versus the cohort size as well as (ii) the validity of results produced. The dataset, as described by Schleusener et al. (2017) - is publicly available under the European Nucleotide Archive (ENA) accession code **PRJEB7727**. Furthermore, Kohl et al. (2018) - analyzed 91 samples for resistance and lineage profiling in a comparative setting as part of Supplementary material of the original publication. These IDs are shared in Supplemental data.

The data submitted to ENA lists 133 paired-end FASTQ files obtained from *Mycobacterium tuberculosis* cultured samples (Supplemental data), and upon meticulous examination, we observed that (i) certain samples had several matching `experiment_accession` entries. The Supplemental data presents a frequency count of `secondary_sample_accession` in relation to `experiment_accession` ERS IDs) (ii) the samples utilized by Kohl et al. (2018) had differences compared to those reported by the ENA project `PRJEB7727`, and (iii) one sample, `ERS457325` lacked any corresponding files in the indicated ENA project. Consequently, to minimize the effect of confounding factors, we opted to deduplicate the samples instead of merging those with identical accession numbers. This was done by utilizing the very first appearance of specific `secondary_sample_accession` IDs from the files acquired from the ENA project and by excluding the missing `ERS457325` (`4730-03`) from our analysis, thereby reducing the final dataset to 90 paired-end FASTQ files. The conclusive samplesheet is provided in Supplemental data.

For the scalability analysis, we divided the dataset into six cohorts with an incremental number of samples namely 5, 10, 20, 40, 80 and finally 90 FASTQ paired-end files as shown in Figure 3. Furthermore, the largest dataset, was used for the validation of principal outputs generated by MTBseq and MTBseq-nf (default and parallel) pipelines.

**FIGURE 3.**
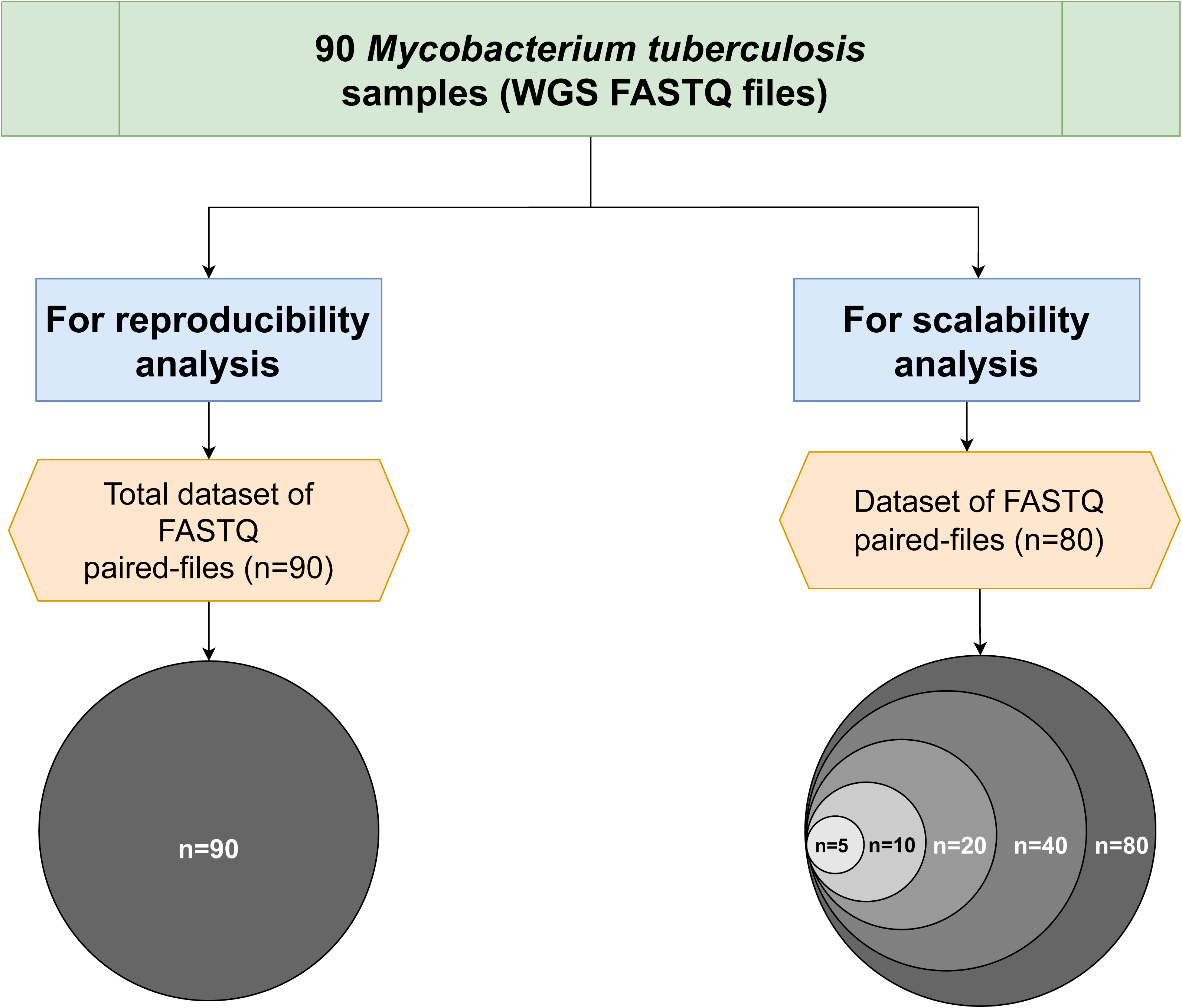
An overview of the 6 subsets, used of the scalability analysis, from the original publication by Kohl et al (2018).

**FIGURE 4.**
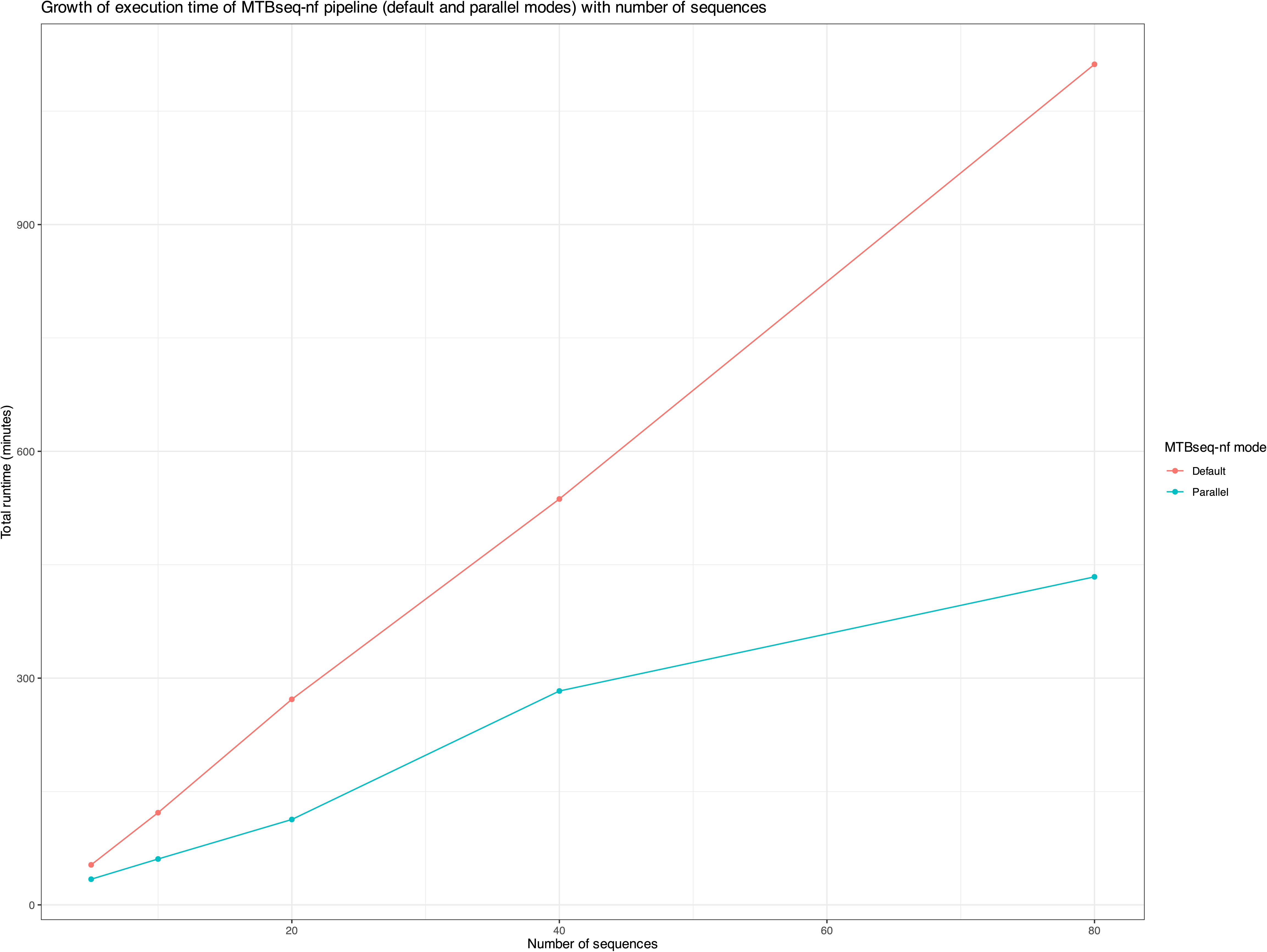
The growth of total runtime of the MTBseq-nf parallel mode and default mode versus the sample size.

### Experimental Setup for Evaluation of Scalability and Reproducibility

To minimize the possibility of regression in MTBseq-nf results (default and parallel modes) in comparison to the MTBseq-standard pipeline. We conducted a triplicated experiments (i) intra-modal analysis to assess reproducibility *within* each mode and (ii) inter-modal analysis to assess reproducibility *across* distinct modes. Consequently, due to this triplicated experimental design, we evaluated the results of nine distinct runs of the pipelines which are highlighted in Supplemental data.

The parameters utilized for these experiments and the associated results are published as part of the Zenodo repository https://doi.org/10.5281/zenodo.14678756, while the names of individual runs and significant parameters are reported in Supplemental data. Moreover, each experiment was conducted independently without employing Nextflow’s resume feature to guarantee fresh execution of each of the individual steps.

To evaluate the scalability of the pipeline, we analyzed the execution time growth for the six specified cohorts using the default mode and parallel mode of MTBseq-nf on these datasets. The executions of MTBSeq-nf in both default and parallel modes were monitored and visualized via the Seqera Platform (Di Tommaso et al., 2025), which serves as a centralized repository for tracking execution metrics and maintaining pipeline configurations.

To evaluate the reproducibility of the different modes of MTBseq-nf, we compared the principal results of inter- and intra-modal triplicated experiments by conducting (i) 3-way diffs using Araxis Merge (team, 2025), (ii) range analysis of numerical data using R language (Team, 2023), along with (iii) phylogenetic tree using IQTREE program (v2.3.4) (Trifinopoulos et al., 2016) as summarized in Supplemental data.

We utilized a 3-way visual diff in order to assess the disparities in the principal results, which are qualitative in nature. For the mapping and variant statistics produced by `TBstats` (including `Mapped Reads` and `SNPs`), we utilized range statistics (min-max analysis) in R to estimate the relative changes across various intra- and inter-modal studies. The scripts utilized for the analysis and visualizations are available in the following repository: https://github.com/abhi18av-phd-projects/mtbseq-nf-publication-analysis.

## RESULTS

### Thematic improvements in MTBseq-nf

The MTBseq-nf wrapper pipeline offers substantial enhancements over the MTBseq-standard pipeline, including an innovative parallel mode that significantly reduces overall execution time for large cohorts. The implemented features span four key themes: (1) user-friendliness, (2) scalability, (3) reproducibility, and (4) maintainability, as highlighted in Table 1 and detailed in Supplemental data.

MTBseq-nf leverages the standardized template from the nf-core community, providing numerous advantages including: (i) configuration files (dotfiles) that address code quality, linting, and testing requirements; (ii) integration with the nf-core/configs project for portability across institutional infrastructures; (iii) access to well-tested modules from the nf-core/modules project; and (iv) a graphical user interface through the nf-schema project for users less familiar with command-line operations (Supplemental data).

### Reproducibility analysis of intra-modal comparison

For the intra-modal comparison of the three pipeline-mode combinations, we conducted triplicated experiments with identical infrastructure, dataset, and parameters as described in Supplemental data.

As summarized in Table 2, the classification and SNP distance matrix results demonstrated consistency across experiments, with differences limited to execution dates, which was expected as successive runs could only begin after previous runs completed. Similarly, the phylogenetic trees remained stable across different experiments. The cluster groups, which are assigned labels based on the SNP matrix, showed some variation in exact labeling across different runs of the TBgroups step, attributable to the inherent nature of the agglomerative algorithm used, though the actual grouping of samples remained accurate.

**TABLE 2.** Summary of intra-modal analysis of principal outputs of triplicated runs. Notably, MTBseq-nf (parallel) mode does not have any differences in the principal results.

| Principal output | MTBseq (standard) | MTBseq-nf (default mode) | MTBseq-nf (parallel mode) |
| --- | --- | --- | --- |
| Classification | No differences | No differences | No differences |
| SNP distance matrix | No differences | No differences | No differences |
| Phylogenetic tree | No differences | No differences | No differences |
| Cluster groups | Consistent agglomeration | Consistent agglomeration | Consistent agglomeration |
| Statistics | Minor differences | Minor differences | No differences |

The intra-modal comparisons, Figure 5, some numeric fields in MTBseq statistics report showed differences in triplicated runs of MTBseq and MTBseq-nf default mode, as shown in Figure 5a and Figure 5b. However, for MTBseq-nf parallel mode, we did not observe variations across any of the numerical fields indicating complete reproducibility of results in the `parallel` mode of MTBseq-nf pipeline.

**FIGURE 5.**
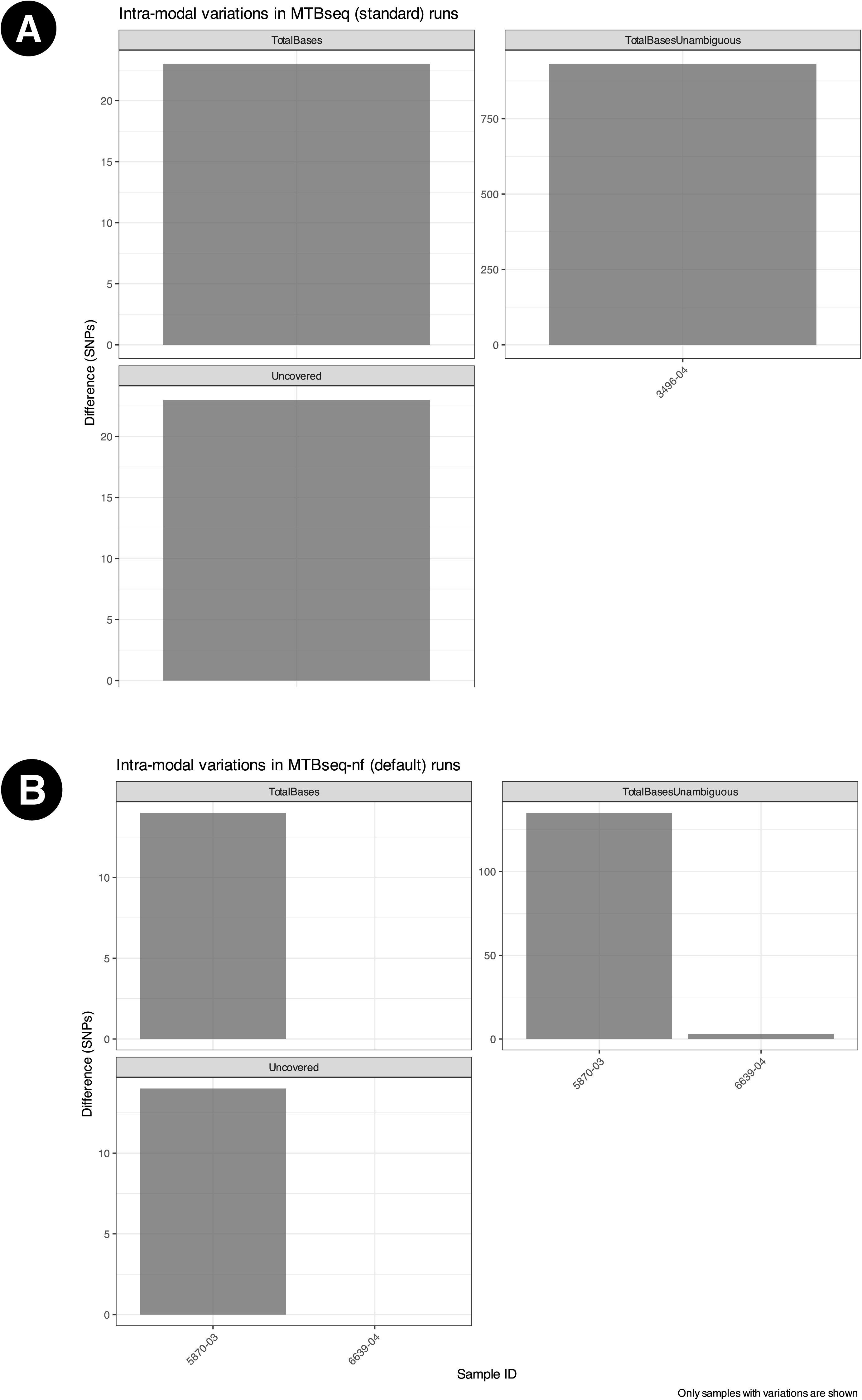
Intra-modal variations across triplicated runs for (a) MTBseq-standard, observed only for three samples (b) MTBseq-nf (default) across triplicated runs, observed only for two sample. No differences were observed across different runs of MTBseq-nf parallel mode.

### Reproducibility analysis of inter-modal comparison

The inter-modal variation has been summarized in Figures-6. Furthermore, the UpSet plots in Figure 7 illustrates the variation in specific columns and corresponding number of samples.

**FIGURE 6.**
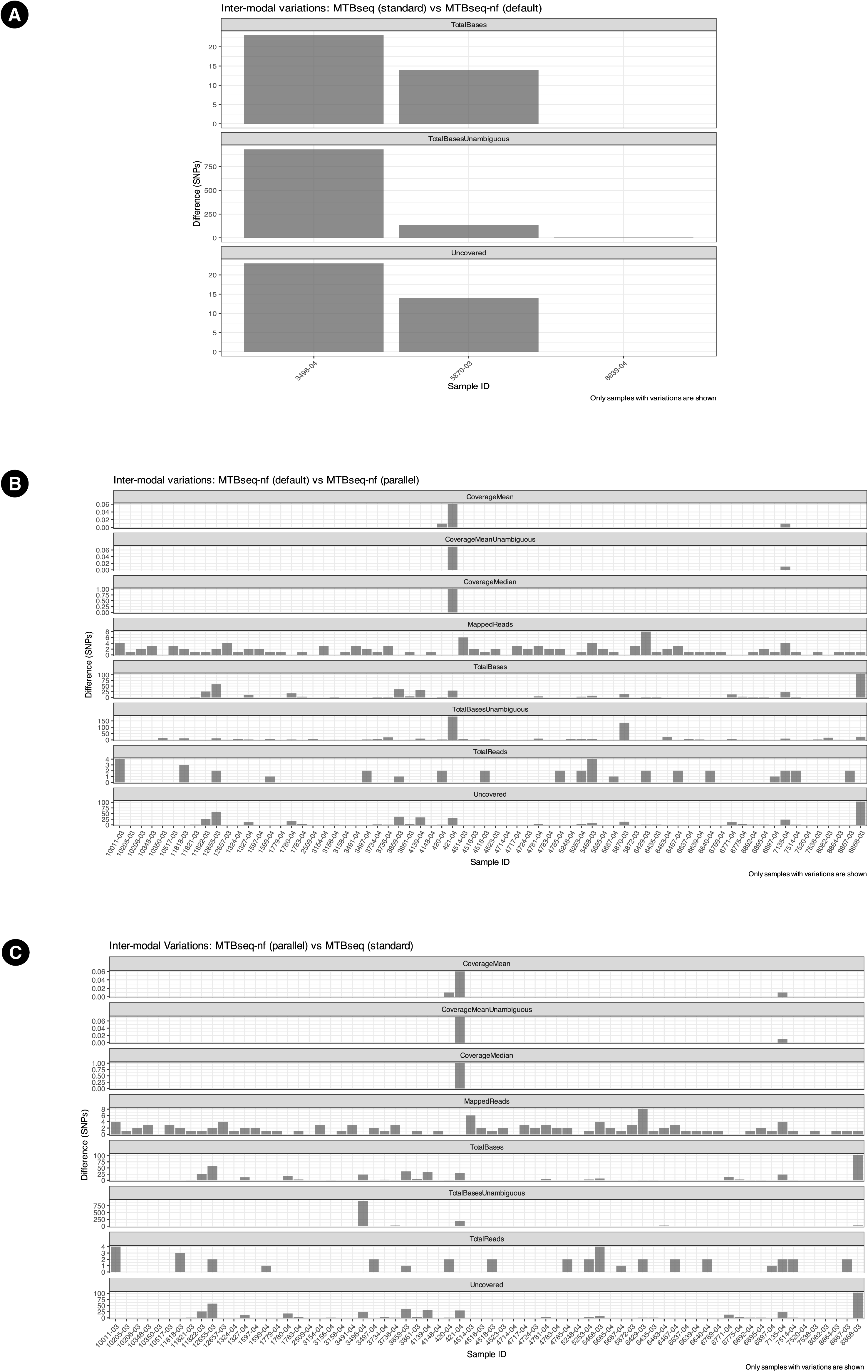
Inter-modal variations between (a) MTBseq-standard and MTBseq-nf (default), wherein only 3 samples has variations. (b) MTBseqs-nf (default) and MTBseq-nf (parallel). (c) MTBseq-nf (parallel) and MTBseq-standard.

**FIGURE 7.**
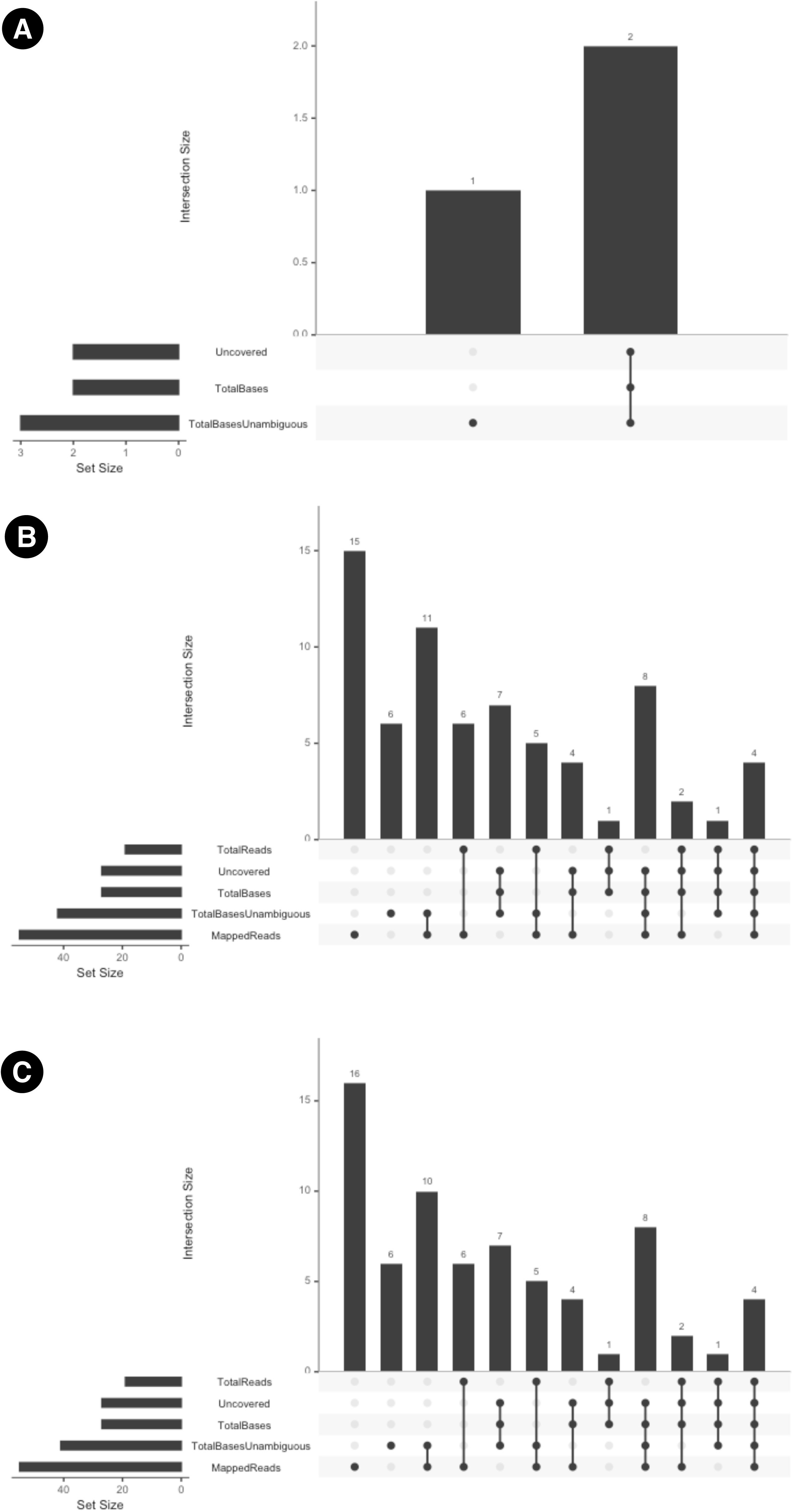
UpSet plots which summarize the inter-modal variations between (a) MTBseq-stan- dard and MTBseq-nf (default). (b) MTBseqs-nf (default) and MTBseq-nf (parallel). (c) MTBseq-nf (parallel) and MTBseq-standard.

**MTBseq-standard vs MTBseq-nf (default).** Figure 6a highlights the numerical variation in SNP counts between MTBseq-standard and MTBseq-nf (default) runs across the fields of Uncovered, TotalBasesUnambguous, and TotalBases and demonstrates that the range of variation between MTBseq-standard and MTBseq-nf (default) is very small.

**MTBseq-nf (parallel) vs MTBseq-nf (default).** When comparing MTBseq-nf (parallel) with MTBseq-nf (default), we observed variations similar to the intra-modal variation of MTBseq- standard triplicated runs, as shown in Figure 5b, and Figure 5c.

**MTBseq-nf (parallel) vs MTBseq-standard.** MTBseq-nf (parallel) vs MTBseq-standard:

For a comprehensive analysis, we compared MTBseq-nf (parallel) with MTBseq-standard, despite the latter being functionally equivalent to MTBseq-nf (default). This comparison is presented in Figure 5e, and Figure 5d.

### Scalability analysis of MTBseq-nf (default) and MTBseq-nf (parallel)

To evaluate execution time growth, we included only one specific run for each mode of MTBseq-nf and excluded MTBseq-standard from the analysis due to the manual intervention step required after TBfull before proceeding to TBstats, TBgroups, and TBamend steps. Since MTBseq-standard and MTBseq-nf (default) are functionally equivalent, comparing total execution time between MTBseq-nf (default) and MTBseq-nf (parallel) was sufficient to analyze the growth curves.

As demonstrated in Figure 4, when processing more than 10 FASTQ paired-end files, the execution time of MTBseq-nf (default) is at least double that of MTBseq-nf (parallel), highlighting the significant performance benefits of the parallel mode optimizations. This difference becomes increasingly pronounced as the sample size grows, making MTBseq-nf (parallel) particularly valuable for large-scale genomic analyses.

## DISCUSSION

The MTBseq-nf pipeline represents a significant optimization effort to modernize the widely adopted MTBseq-standard pipeline for analyzing WGS data from the *M. tuberculosis* culture. In doing so, we address numerous pain points experienced by users of the original MTBseq pipeline, especially in high-throughput and diverse computing environments. One of the major advancements introduced by MTBseq-nf is the optional `--parallel` execution mode. By decoupling sequential dependencies and capitalizing on the Nextflows inherent parallelization features, MTBseq-nf significantly reduces the overall runtime of the pipeline. Our empirical analysis indicated that the execution time scales much more gracefully in the parallel mode, particularly as the number of samples increases — a critical feature for genomic surveillance programs processing hundreds or thousands of isolates.

The task-level isolation introduced by `parallel` mode ensures that each primitive step has sufficient memory (and CPU) during its execution and as soon as the step completes, the relevant docker container exits and the docker container engine ensures that the computational resources of the host system are released and made available for other queued tasks (Di Tommaso et al., 2015). Whereas in the `TBfull` step within MTBseq and MTBseq-nf (default), the consumed memory is not immediately released when the previous process completes as the computations are done sequentially within the same container, therefore the subsequent processes have slightly reduced access to memory, which has an impact upon the memory-intensive computations required by `GATK`.

This improvement with MTBseq-nf `parallel` is especially beneficial for the integration of bioinformatic analysis in routine diagnostics and surveillance work especially in resource-constrained environments such as Low- and Middle-income countries (LMICs), where TB is endemic and timely analysis is vital, and where infrastructure may not be suited to long, sequential compute jobs. Furthermore, the ability to execute the MTBseq-nf pipeline across laptops, servers, HPC clusters, and cloud batch systems allows for unprecedented flexibility and broader adoption.

Reproducibility is a cornerstone of bioinformatics pipelines, particularly when used in clinical and public health contexts. MTBseq-nf leverages Nextflow’s declarative parameter feature, integration with container systems (Kadri et al., 2022) such as Docker, and in-built caching mechanisms to rapidly deliver reproducible results across varied environments. The triplicated intra-modal experiments confirmed the consistency of principal outputs across different runs of MTBseq-nf, particularly in the parallel mode, which showed no measurable variation in statistics or results.

Users benefit from the baseline improvements within the MTBseq-nf (default) mode, without making use of the `parallel` feature. Results obtained from intra-modal (Figure 5a and Figure 5b) analysis as well inter-modal analysis (Figure 6a, Figure 7a) of MTBseq-nf (default) and MTBseq-standard, establish the functional equivalence the results generated by the MTBseq-nf (default) compared with MTBseq-standard.

Moreover, discrepancies in output observed between sequential and parallel executions, such as minor differences in `TBstats` reports, are likely attributable to the memory-intensive behavior of tools like GATK when executed sequentially on all samples in shared containers. In contrast, the task-level isolation in the parallel mode ensures more consistent memory allocation, leading to increased stability of result across executions.

In architecting MTBseq-nf on top of MTBseq-standard, we have prioritized a straight-forward user experience. By adopting the nf-core best practices, users benefit from structured configuration files, standardized output, and simplified parameter management. Features like an explicit sample sheet, an optional Graphic User Interface (GUI) through the `nf-schema` project, and remote execution monitoring and infrastructure management (via the Seqera platform) (Di Tommaso et al., 2025) further enhance usability, especially for users less familiar with the command line. Another valuable addition is the inclusion of modules like `FastQC` and `MultiQC` (P. Ewels et al., 2016), which offer quality control summaries that are now seamlessly integrated into the pipeline without manual intervention. In contrast, integrating such tools into MTBseq-standard would require creation, customization, and testing of Perl-wrapped modules.

The adoption of the nf-core pipeline template ensures the future viability of MTBseq-nf by allowing rapid integration of updates and modules. Unlike the traditional software architecture of the MTBseq-standard pipeline, which makes maintenance and extension daunting, especially for newcomers, to the project; the modular design of MTBseq-nf, owing to the nf-core best-practices template, aligns with modern software engineering practices, including continuous integration and delivery (CI/CD), unit testing (e.g., using nf-test), and community contributions via Github. This design ensures that MTBseq-nf can evolve in response to user needs, bug reports, and new tool integrations — such as IQTREE, which, although not currently included, can be added as an integrated module in future iterations depending on research requirements.

In terms of limitations, despite its many strengths, MTBseq-nf inherits the core computational logic of MTBseq-standard, limiting opportunities for deeper architectural optimization. The reliance on MTBseq’s Perl-based modules, while preserving backward compatibility, poses a ceiling on how extensively the workflow can be modernized. Future work could involve translating these steps into native Nextflow processes or subworkflows, which would improve maintainability and support pluggable alternative tools for the individual steps of MTBseq-standard for alignment and variant calling tools.

Additionally, minor variations in SNP clustering labels across runs, an artifact of the agglomerative clustering algorithm, do not impact downstream phylogenetic analyses but underscore the importance of clear documentation and careful interpretation of clustering outputs. The efficiency gains offered by MTBseq-nf reduce execution time, cost of analysis and contribute to sustainability goals by lowering computational carbon footprints, an increasingly relevant consideration in bioinformatics research (Grealey et al., 2022; Lannelongue et al., 2023). Quantifying and comparing emissions across pipelines could be an interesting area for future investigation.

Furthermore, the versatility of MTBseq-nf enables its deployment in both high-resource and constrained settings, supporting TB control efforts globally. By simplifying complex analyses and promoting reproducibility, the pipeline aligns with the broader goals of precision public health and genomic epidemiology.

## CONCLUSION

In light of the (i) rapidly increasing WGS data for *M. tuberculosis* (ii) along with its applicability in genomic surveillance and precision medicine; the MTBseq-nf offers a more scalable, portable and user-friendly alternative to the MTBseq-pipeline. The MTBseq-nf wrapper optimizes the MTBseq-standard pipeline particularly towards an efficient usage of computational resources for genomic “Big Data” analysis for. In addition, it also improves the user-friendliness, reproducibility, scalability and customisablity as compared to the MTBseq-standard pipeline. These enhancements in MTBseq-nf adds to the versatality of that the original MTBseq-standard as a suitable alternative, not only for routine usage in resource-rich environments, but particularly in resource-constraint LMIC settings where fine-grained control over resource usage is necessary.

## Supporting information

SD-1 Auto-generated graphical user interface for the MTBseq-nf pipeline on Seqera Platform (formerly Nextflow Tower).

SD-2 Summary of validation technique used for principal results.

SD-3.01 Samplesheet ERS (from original publication Kohl et al, 2018 - Supplementary Table S2)

SD-3.02 Report from ENA https://www.ebi.ac.uk/ena/browser/view/PRJEB7727

SD-3.03 Frequency of ERS IDs from original publication

SD-3.04 Final samplesheet used for the analysis

SD-3.05 Metadata report for the Bioproject, generated using nf-core/fetchngs

SD-4 Bioconda recipe for MTBseq v1.1.0: MTBseq_conda_recipe.yml

SD-5 An overview of key enhancements in MTBseq-nf, Nextflow wrapper for the original MTBseq pipeline.

SD-6 Intra-modal analysis, with 3-way HTML diff reports generated by Araxis merge software

SD-7 Growth of total execution time of different modes of MTBseq-nf for 5 datasets with increasing cohort size.

SD-8 Summary of different executions of MTBseq and MTBseq-nf in the triplicated set of experiments.

## ACKNOWLEDGEMENTS

We extend our gratitude towards the authors of MTBseq-standard pipeline, Kohl et al. (2018), for the initiative of publishing MTBSeq pipeline with open source on Github.

## DATA AVAILABILITY STATEMENT

The dataset published on Zenodo https://doi.org/10.5281/zenodo.14678756 contains results and metadata for the nine runs of MTBseq-standard, MTBseq-nf (default) and MTBseq-nf (parallel) including (i) MultiQC reports (ii) runtime metrics (iii) pipeline parameters (iv) results of all experiments.

## CODE AVAILABILITY STATEMENT

MTBseq-nf pipeline https://zenodo.org/records/15234640

MTBseq-nf analysis scripts https://github.com/abhi18av-phd-projects/pub-mtbseq-nf

## GENERATIVE AI STATEMENT

We utilized the Anthropic Claude and QuillBot tool for grammar checks, paraphrasing and summarization.

## MANUSCRIPT ASSETS INDEX

CODE - GITHUB

MTBseq-nf pipeline https://zenodo.org/records/15234640

MTBseq-nf analysis scripts https://github.com/abhi18av-phd-projects/pub-mtbseq-nf

DATA - ZENODO

The dataset published on Zenodo https://doi.org/10.5281/zenodo.14678756 contains results and metadata for the nine runs of MTBseq-standard, MTBseq-nf (default) and MTBseq-nf (parallel) including (i) MultiQC reports (ii) runtime metrics (iii) pipeline parameters (iv) results of all experiments.

## DATA - SUPPLEMENTAL

**SD-1** Auto-generated graphical user interface for the MTBseq-nf pipeline on Seqera Platform (formerly Nextflow Tower).

1

**SD-2** Summary of validation technique used for principal results.

**SD-3** Validation dataset

• **SD-3.01** Samplesheet ERS (from original publication Kohl et al, 2018 - Supplementary Table S2)

• **SD-3.02** Report from ENA https://www.ebi.ac.uk/ena/browser/view/PRJEB7727

• **SD-3.03** Frequency of ERS IDs from original publication

• **SD-3.04** Final samplesheet used for the analysis

• **SD-3.05** Metadata report for the Bioproject, generated using nf-core/fetchngs

**SD-4** Bioconda recipe for MTBseq v1.1.0: MTBseq_conda_recipe.yml

**SD-5** An overview of key enhancements in MTBseq-nf, Nextflow wrapper for the original MTBseq pipeline.

**SD-6** Intra-modal analysis, with 3-way HTML diff reports generated by Araxis merge software

• **SD-6.01** intra-modal-araxiscompare-pub-90samples-mtbseq-nf-parallel- runs-classification.html

• **SD-6.02** intra-modal-araxiscompare-pub-90samples-mtbseq-nf-parallel-runs-cluster-groups.html

• **SD-6.03** intra-modal-araxiscompare-pub-90samples-mtbseq-nf-parallel-runs-snp-matrix.html

• **SD-6.04** intra-modal-araxiscompare-pub-90samples-mtbseq-nf-parallel-runs-statistics.html

• **SD-6.05** intra-modal-araxiscompare-pub-90samples-mtbseq-nf-runs-classification.html

• **SD-6.06** intra-modal-araxiscompare-pub-90samples-mtbseq-nf-runs-cluster-groups.html

• **SD-6.07** intra-modal-araxiscompare-pub-90samples-mtbseq-nf-runs-snp-matrix.html

• **SD-6.08** intra-modal-araxiscompare-pub-90samples-mtbseq-nf-runs-statistics.html

• **SD-6.09** intra-modal-araxiscompare-pub-90samples-mtbseq-standard-runs-classification.html

• **SD-6.10** intra-modal-araxiscompare-pub-90samples-mtbseq-standard-runs-cluster-groups.html

• **SD-6.11** intra-modal-araxiscompare-pub-90samples-mtbseq-standard-runs-snp-matrix.html

• **SD-6.12** intra-modal-araxiscompare-pub-90samples-mtbseq-standard-runs-statistics.html

**SD-7** Growth of total execution time of different modes of MTBseq-nf for 5 datasets with increasing cohort size.

## AUTHOR ROLES AND CONTRIBUTIONS

Author roles were classified using the Contributor Role Taxonomy (CRediT; https://credit.niso.org/) as follows: *Abhinav Sharma\**: conceptualization, funding acquisition, investigation (equal), methodology, project administration, software (equal), validation, writing – original draft, writing – review & editing, and visualization. *Davi Josué Marcon\**: writing – original draft, software (equal), investigation (equal), and writing – review & editing. *Johannes Loubser*: writing – review & editing and validation. *Karla Valéria Batista Lima*: validation, writing – review & editing, and resources. *Gian van der Spuy#*: visualization, conceptualization, and supervision (equal). *Emilyn Costa Conceição#*: supervision (equal), visualization, and conceptualization

