## Supplementary figures and images for "MTBseq-nf: Enabling Scalable Tuberculosis Genomics “Big Data” Analysis through a User-Friendly Nextflow Wrapper for MTBseq pipeline"

### SD-1 Auto-generated graphical user interface for the MTBseq-nf pipeline on Seqera Platform (formerly Nextflow Tower).

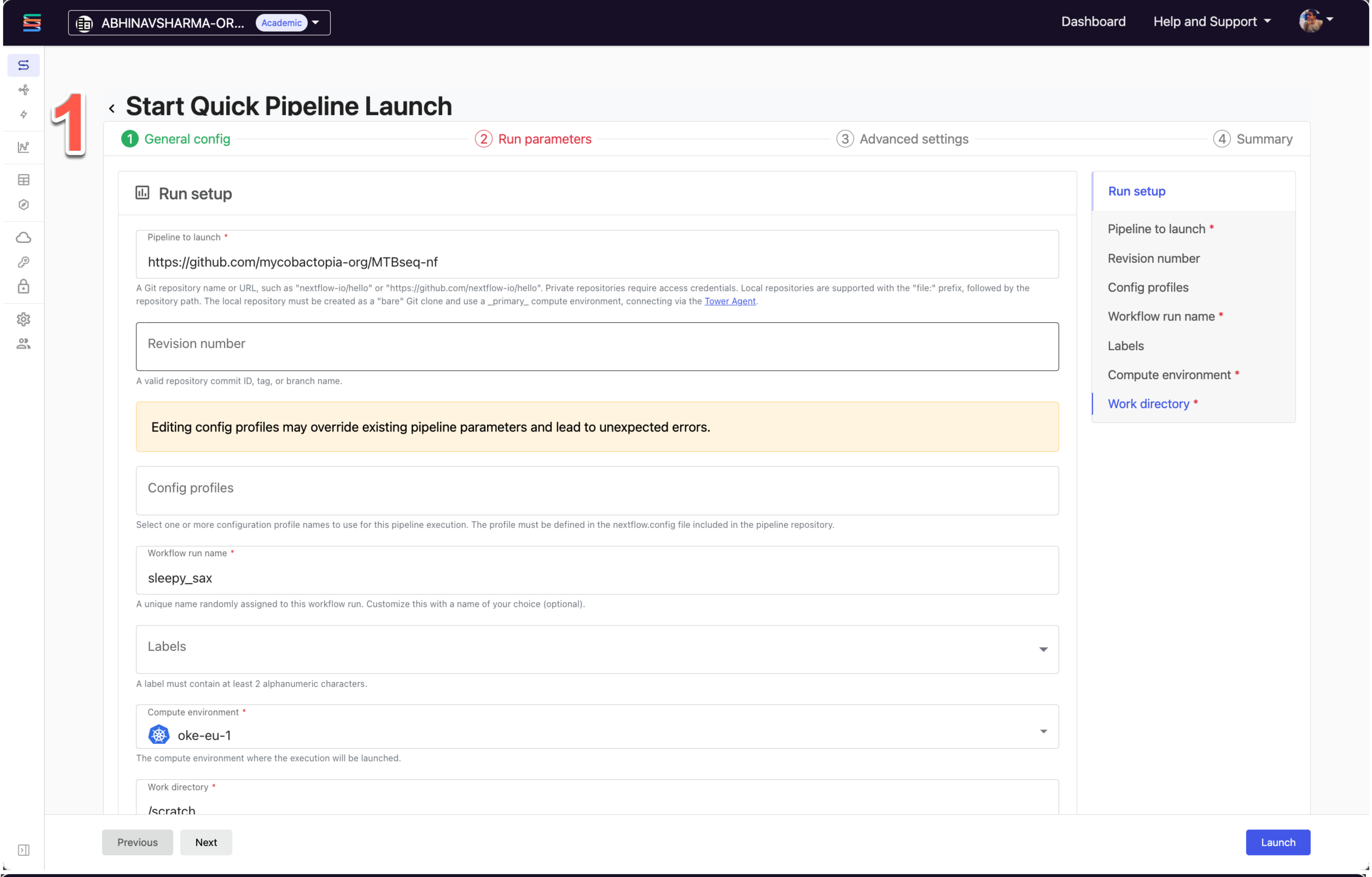

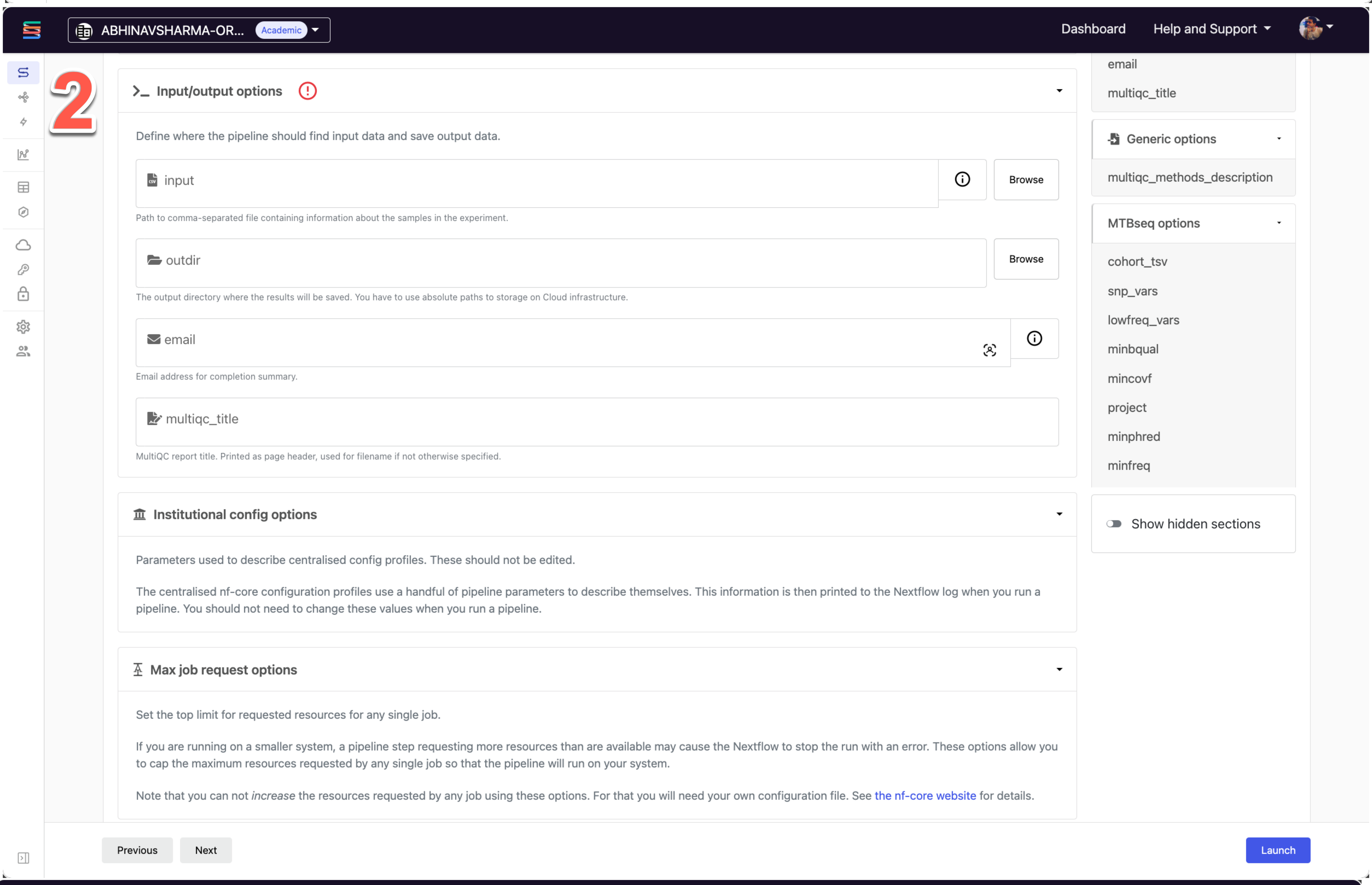

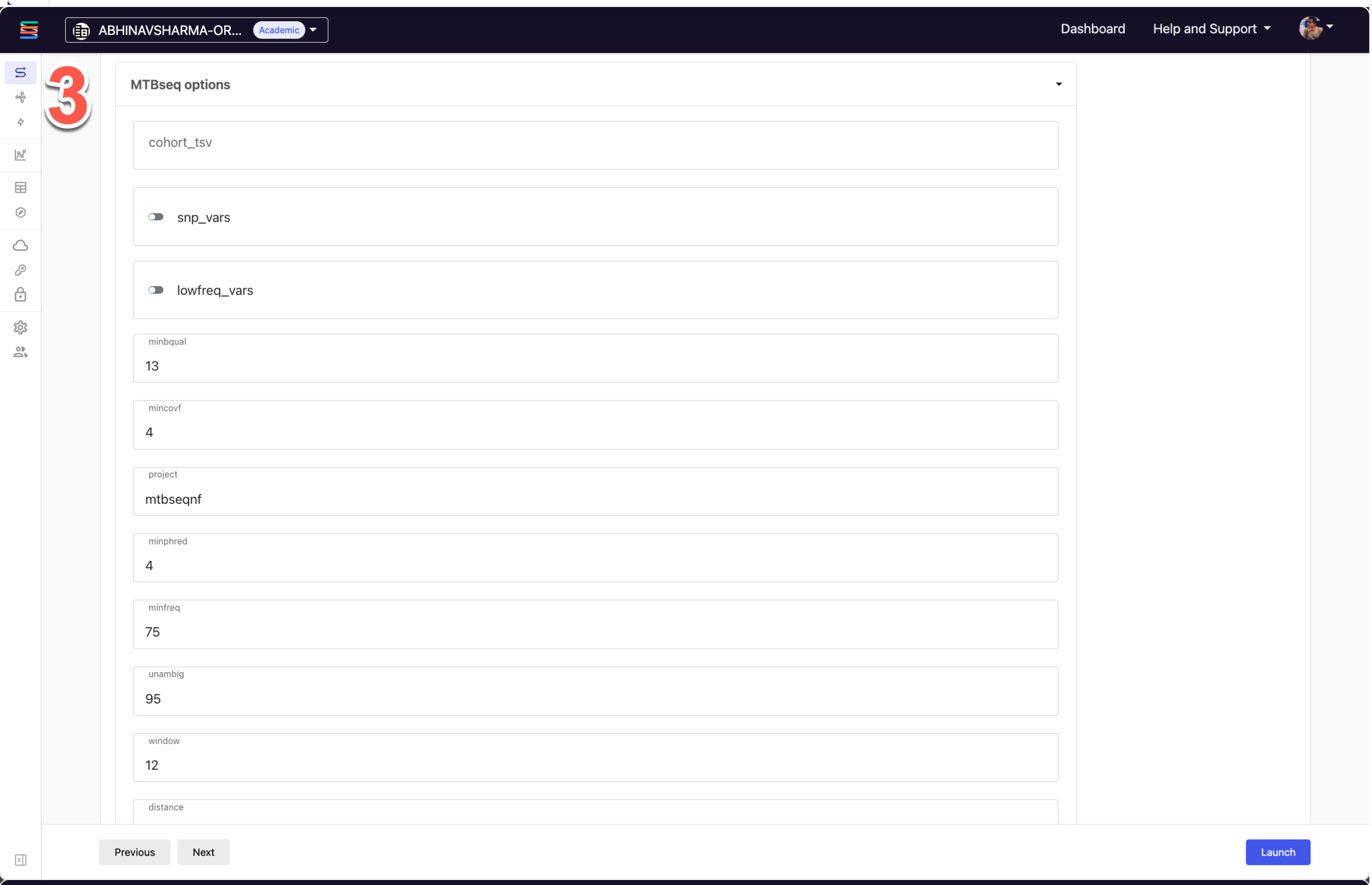

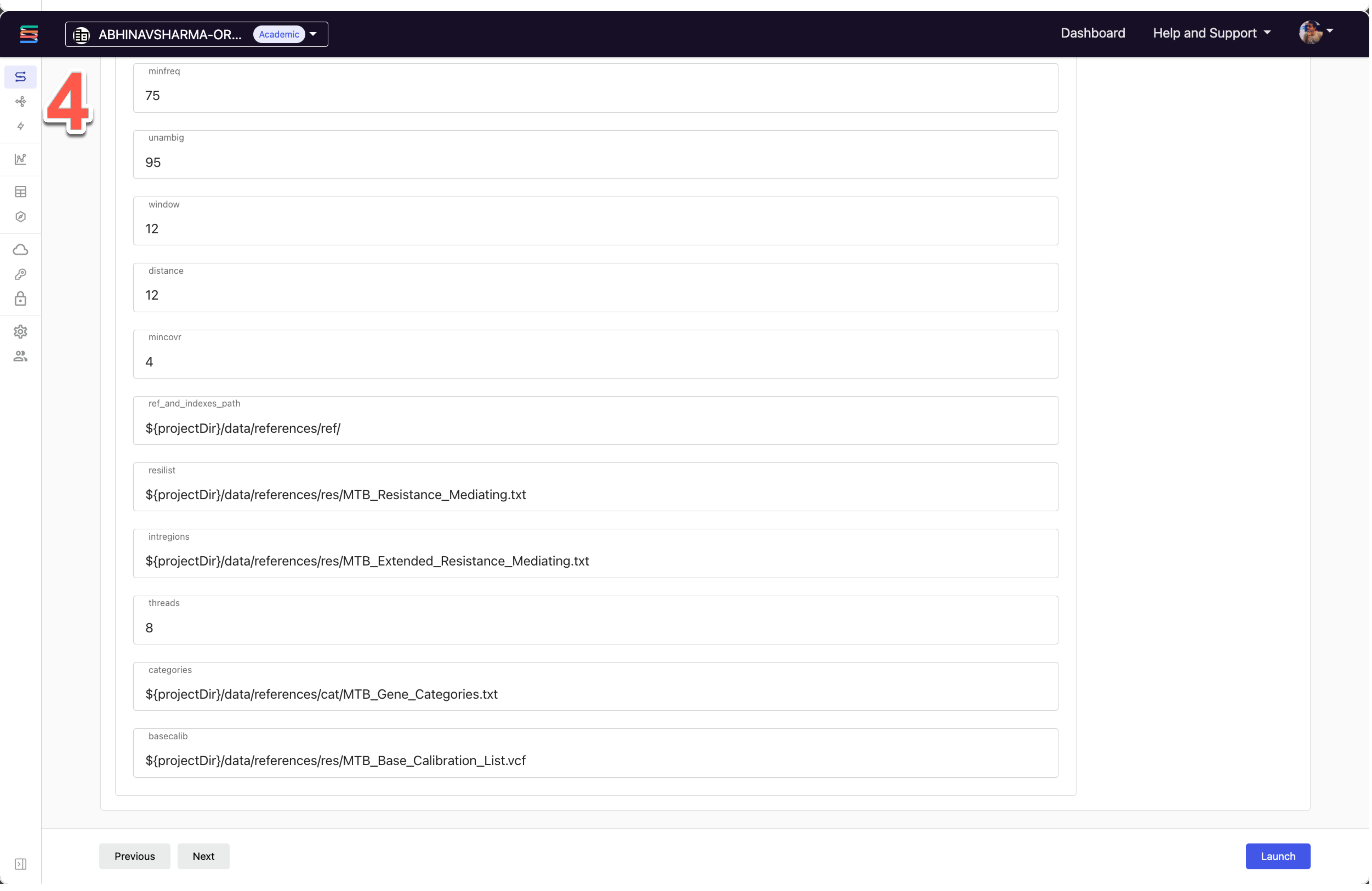
