## Supplementary material for "MTBseq-nf: Enabling Scalable Tuberculosis Genomics “Big Data” Analysis through a User-Friendly Nextflow Wrapper for MTBseq pipeline": SD-2 Summary of validation technique used for principal results.

**Summary of validation techniques used for the different results generated by the MTBseq pipeline.**

| Report name | Validation technique |
|---|---|
| Classification | 3-way diff report |
| SNP distance matrix | 3-way diff report |
| Cluster groups | 3-way diff report |
| Phylogenetic tree | Tree generated by IQTREE |
| Mapping and variant statistics | (i) 3-way diff report and (ii) statistical analysis |
