## Supplementary material for "MTBseq-nf: Enabling Scalable Tuberculosis Genomics “Big Data” Analysis through a User-Friendly Nextflow Wrapper for MTBseq pipeline": SD-5 An overview of key enhancements in MTBseq-nf, Nextflow wrapper for the original MTBseq pipeline.

| Theme | Feature | MTBseq-standard | MTBseq-nf |
|---|---|---|---|
| User-friendliness | Ease of download | Can be downloaded through bioconda or biocontainers. | Given that Nextflow is installed, "nextflow pull" command should suffice for all assets. |
| User-friendliness | Explicit Samplesheet | No explicit samplesheet. The samples are expected to be in the pipeline execution directory with the expected naming convention. | Users must provide a samplesheet pointing to the samples in any location and the samples will be "soft renamed" automatically to fit the MTBseq requirement. |
| User-friendliness | Graphical user interface | None, the user must provide all parameters on the command line. | User can make use of the Seqera Platform or nf-core tools to provide the parameters graphically. In addition, these parameters will be validation prior to the execution of the pipeline. |
| User-friendliness | MultiQC Summary report | The MTBseq pipeline publishes 4 principal results in different directories. | In addition to the individual files, MTBseq-nf produces a compiled MultiQC report with visualizations of these principal results. |
| User-friendliness | Remote monitoring | None, the user must login to the server/cluster and check whether the pipeline has finished or not. | The user can optionally monitor the execution of the pipeline on freely available monitoring capability of Seqera Platform and share the link with fellow researchers. |
| User-friendliness | Manual steps | Users must create a 2-column TSV file for cohort level steps of the MTBseq pipeline. | MTBseq-nf can optionally auto-generate the 2-column TSV file for cohort-level steps and continues onwards with the execution of cohort steps. |
| User-friendliness | Flexible output location | The pipeline produces the resulting files in the pipeline execution directory itself. | The user can opt publish the results in the desired local or cloud location. |
| Maintainability | Extensibility | It is necessary to make changes to the perl5 codebase to add new tools to the pipeline. | The addition of new tested modules is straightforward, we added FASTQC and MULTIQC without changing the baseline MTBseq perl5 codebase. |
| Maintainability | Module testing | Only the integration tests can be conducted with the current design. Module testing would require addition of a Perl specific testing framework. | The modules can either be downloaded or generated using the nf-core command line utility, which generates all the relevant files for module-level testing. |
| Maintainability | Test dataset | The MTBseq pipeline does not come with a test dataset, the users are expected to either test with their own sample or download the samples from ENA used in the original MTBseq publication. | The MTBseq-nf pipeline provides test and test_full profiles for users to make sure that the pipeline is properly configured. The pipeline downloads the test data automatically, derived from the original publication. |
| Scalability | Parallel execution | The individual steps rely upon a "foreach" loop. | The pipeline allows decoupling of individual steps and can analyze the samples in parallel. |
| Scalability | HPC compatibility | The pipeline is usable using traditional hand-crafted job scripts. | Nextflow generates the job scripts for any HPC platform or for cloud executor platforms. In addition, users can customize the precise queue level constrains for each process in MTBseq-nf. |
| Scalability | Resource allocation | The pipeline is capped at maximum of 8 cpus with a minimum recommended memory of FIXME | Users can allocate the resources for a task with precision through process selectors in Nextflow configuration. |
| Scalability | Dynamic retries | None, the user must manually resize the resource constrains upon the failure of pipeline. | MTBseq-nf can automatically resubmit/retry the jobs with higher resources constraints. |
| Scalability | Reduced Data footprint | The TBfull generates the intermediate files as well as the main results in directory of invocation. | The user can make use of the publishDir and workDir configurations in Nextflow to store only the principal results. |
| Reproducibility | Declarative parameters file | The user must provide any custom parameters on the command line using the typical command line flags. | In addition to the flags, the users can provide the parameters in a YAML file or on the command line. |
| Reproducibility | Portability | The users must make decisions and necessary setup for executing the pipeline such as the use of conda, docker, singularity etc. | The nf-core template facilitates easy switching of the package manager or container platform, and the pipeline can download these assets during execution time. In addition, users can rely upon nf-core/configs project for their institutional configuration. |
| Reproducibility | Save intermediate files | No, the TBfull step deletes the intermediate files such as the output of FIXME | Yes, the user can either opt to store all intermediate files in results, which can facilitate troubleshooting or opt to store only the most relevant results. |
