## Supplementary material for "MTBseq-nf: Enabling Scalable Tuberculosis Genomics “Big Data” Analysis through a User-Friendly Nextflow Wrapper for MTBseq pipeline": SD-6 Intra-modal analysis, with 3-way HTML diff reports generated by Araxis merge software: SD-6-02-intra-modal-araxiscompare-pub-90samples-mtbseq-standard-runs-cluster-groups.html

xml version="1.0" encoding="utf-8"?


Araxis Merge File Comparison Report 

### **Araxis Merge File Comparison Report**

Produced by **Araxis Merge** on **2024/12/16, 16:30 GMT+02:00**. See www.araxis.com for information about Merge. This report uses XHTML and CSS2, and is best viewed with a modern standards-compliant browser. For optimum results when printing this report, use landscape orientation and enable printing of background images and colours in your browser.

##### **1. Files compared**

| # | Location | File | Last Modified |
| --- | --- | --- | --- |
| 1 | /Users/abhi/projects/MTBseq-nf/\_resources/publication/manuscript-and-analysis/v2/pub-90samples-mtbseq-standard-run1/Groups | mtbseqnf\_joint\_cf4\_cr4\_fr75\_ph4\_samples90\_amended\_u95\_phylo\_w12\_d12.groups | 2024/10/15, 17:41 GMT+02:00 |
| 2 | /Users/abhi/projects/MTBseq-nf/\_resources/publication/manuscript-and-analysis/v2/pub-90samples-mtbseq-standard-run2/Groups | mtbseqnf\_joint\_cf4\_cr4\_fr75\_ph4\_samples90\_amended\_u95\_phylo\_w12\_d12.groups | 2024/10/16, 05:25 GMT+02:00 |
| 3 | /Users/abhi/projects/MTBseq-nf/\_resources/publication/manuscript-and-analysis/v2/pub-90samples-mtbseq-standard-run3/Groups | mtbseqnf\_joint\_cf4\_cr4\_fr75\_ph4\_samples90\_amended\_u95\_phylo\_w12\_d12.groups | 2024/10/16, 20:21 GMT+02:00 |
| **Note:** Merge considers the second file to be the common ancestor of the others. | | | |

##### **2. Comparison summary**

| Description | Between Files 1 and 2 | | Between Files 2 and 3 | | Relative to Common Ancestor | |
| --- | --- | --- | --- | --- | --- | --- |
|  | Text Blocks | Lines | Text Blocks | Lines | Text Blocks | Lines |
| Unchanged | 1 | 228 | 4 | 212 |  |  |
| Changed | 0 | 0 | 2 | 14 | 2 | 14 |
| Inserted | 0 | 0 | 0 | 0 | 0 | 0 |
| Removed | 0 | 0 | 1 | 2 | 1 | 2 |
| **Note:** An automatic merge would leave 0 conflict(s). | | | | | | |

##### **3. Comparison options**

|  |  |
| --- | --- |
| Whitespace | Consecutive whitespace is treated as a single space |
| Character case | Differences in character case are significant |
| Line endings | Differences in line endings (`CR` and `LF` characters) are ignored |
| CR/LF characters | Not shown in the comparison detail |
| Active line-pairing rules |  |

##### **4. Active regular expressions**

No regular expressions were active.

##### **5. Comparison detail**

|  |  |  |  |  |  |  |  |  |  |  |
| --- | --- | --- | --- | --- | --- | --- | --- | --- | --- | --- |
| 1 |  | ### Output as groups: |  | 1 |  | ### Output as groups: |  | 1 |  | ### Output as groups: |
| 2 |  | > group\_1»  5 |  | 2 |  | > group\_1»  5 |  | 2 |  | > group\_1»  5 |
| 3 |  | 10517-03,1779-04,1780-04,5685-04,6463-04 |  | 3 |  | 10517-03,1779-04,1780-04,5685-04,6463-04 |  | 3 |  | 10517-03,1779-04,1780-04,5685-04,6463-04 |
| 4 |  | > group\_2»  2 |  | 4 |  | > group\_2»  2 |  | 4 |  | > group\_2»  2 |
| 5 |  | 10206-03,3865-03 |  | 5 |  | 10206-03,3865-03 |  | 5 |  | 10206-03,3865-03 |
| 6 |  | > group\_3»  3 |  | 6 |  | > group\_3»  3 |  | 6 |  | > group\_3»  2 |
|  |  |  |  |  |  |  |  | 7 |  | 1597-04,4523-03 |
|  |  |  |  |  |  |  |  | 8 |  | > group\_4»  3 |
| 7 |  | 10207-03,12655-03,3734-04 |  | 7 |  | 10207-03,12655-03,3734-04 |  | 9 |  | 10207-03,12655-03,3734-04 |
| 8 |  | > group\_4»  2 |  | 8 |  | > group\_4»  2 |  |  |  |  |
| 9 |  | 1597-04,4523-03 |  | 9 |  | 1597-04,4523-03 |  |  |  |  |
| 10 |  | > group\_5»  3 |  | 10 |  | > group\_5»  3 |  | 10 |  | > group\_5»  3 |
| 11 |  | 5248-04,7517-04,7538-03 |  | 11 |  | 5248-04,7517-04,7538-03 |  | 11 |  | 5248-04,7517-04,7538-03 |
| 12 |  | > group\_6»  2 |  | 12 |  | > group\_6»  2 |  | 12 |  | > group\_6»  2 |
| 13 |  | 11818-03,4781-04 |  | 13 |  | 11818-03,4781-04 |  | 13 |  | 11818-03,4781-04 |
| 14 |  | > group\_7»  2 |  | 14 |  | > group\_7»  2 |  | 14 |  | > group\_7»  2 |
| 15 |  | 10348-03,6639-04 |  | 15 |  | 10348-03,6639-04 |  | 15 |  | 10348-03,6639-04 |
| 16 |  | > group\_8»  2 |  | 16 |  | > group\_8»  2 |  | 16 |  | > group\_8»  2 |
| 17 |  | 12658-03,7135-04 |  | 17 |  | 12658-03,7135-04 |  | 17 |  | 12658-03,7135-04 |
| 18 |  | > group\_9»  5 |  | 18 |  | > group\_9»  5 |  | 18 |  | > group\_9»  5 |
| 19 |  | 11822-03,1599-04,4148-04,4514-03,6467-04 |  | 19 |  | 11822-03,1599-04,4148-04,4514-03,6467-04 |  | 19 |  | 11822-03,1599-04,4148-04,4514-03,6467-04 |
| 20 |  | > ungrouped»64 |  | 20 |  | > ungrouped»64 |  | 20 |  | > ungrouped»64 |
| 21 |  | 10010-03,10011-03,10012-03,10205-03,10208-03,10349-03,10350-03,11096-03,11097-03,11821-03,12657-03,1322-04,1324-04,1327-04,1783-04,2509-04,3154-04,3156-04,3158-04,3160-04,3491-04,3494-04,3496-04,3497-04,3736-04,3859-03,3861-03,4139-04,4145-04,420-04,421-04,4516-03,4518-03,4712-04,4714-04,4717-04,4724-03,4779-04,4783-04,4785-04,5253-04,5468-03,5472-03,5687-04,5870-03,5872-03,6429-03,6435-03,6637-04,6640-04,6769-04,6771-04,6775-04,6892-04,6895-04,6897-04,7000-03,7514-04,7516-04,7520-04,8082-03,8864-03,8867-03,8868-03 |  | 21 |  | 10010-03,10011-03,10012-03,10205-03,10208-03,10349-03,10350-03,11096-03,11097-03,11821-03,12657-03,1322-04,1324-04,1327-04,1783-04,2509-04,3154-04,3156-04,3158-04,3160-04,3491-04,3494-04,3496-04,3497-04,3736-04,3859-03,3861-03,4139-04,4145-04,420-04,421-04,4516-03,4518-03,4712-04,4714-04,4717-04,4724-03,4779-04,4783-04,4785-04,5253-04,5468-03,5472-03,5687-04,5870-03,5872-03,6429-03,6435-03,6637-04,6640-04,6769-04,6771-04,6775-04,6892-04,6895-04,6897-04,7000-03,7514-04,7516-04,7520-04,8082-03,8864-03,8867-03,8868-03 |  | 21 |  | 10010-03,10011-03,10012-03,10205-03,10208-03,10349-03,10350-03,11096-03,11097-03,11821-03,12657-03,1322-04,1324-04,1327-04,1783-04,2509-04,3154-04,3156-04,3158-04,3160-04,3491-04,3494-04,3496-04,3497-04,3736-04,3859-03,3861-03,4139-04,4145-04,420-04,421-04,4516-03,4518-03,4712-04,4714-04,4717-04,4724-03,4779-04,4783-04,4785-04,5253-04,5468-03,5472-03,5687-04,5870-03,5872-03,6429-03,6435-03,6637-04,6640-04,6769-04,6771-04,6775-04,6892-04,6895-04,6897-04,7000-03,7514-04,7516-04,7520-04,8082-03,8864-03,8867-03,8868-03 |
| 22 |  |  |  | 22 |  |  |  | 22 |  |  |
| 23 |  |  |  | 23 |  |  |  | 23 |  |  |
| 24 |  | ### Output as lists: |  | 24 |  | ### Output as lists: |  | 24 |  | ### Output as lists: |
| 25 |  | 10517-03»   group\_1 |  | 25 |  | 10517-03»   group\_1 |  | 25 |  | 10517-03»   group\_1 |
| 26 |  | 1779-04»group\_1 |  | 26 |  | 1779-04»group\_1 |  | 26 |  | 1779-04»group\_1 |
| 27 |  | 1780-04»group\_1 |  | 27 |  | 1780-04»group\_1 |  | 27 |  | 1780-04»group\_1 |
| 28 |  | 5685-04»group\_1 |  | 28 |  | 5685-04»group\_1 |  | 28 |  | 5685-04»group\_1 |
| 29 |  | 6463-04»group\_1 |  | 29 |  | 6463-04»group\_1 |  | 29 |  | 6463-04»group\_1 |
| 30 |  | 10206-03»   group\_2 |  | 30 |  | 10206-03»   group\_2 |  | 30 |  | 10206-03»   group\_2 |
| 31 |  | 3865-03»group\_2 |  | 31 |  | 3865-03»group\_2 |  | 31 |  | 3865-03»group\_2 |
| 32 |  | 10207-03»   group\_3 |  | 32 |  | 10207-03»   group\_3 |  | 32 |  | 1597-04»group\_3 |
| 33 |  | 12655-03»   group\_3 |  | 33 |  | 12655-03»   group\_3 |  | 33 |  | 4523-03»group\_3 |
| 34 |  | 3734-04»group\_3 |  | 34 |  | 3734-04»group\_3 |  | 34 |  | 10207-03»   group\_4 |
| 35 |  | 1597-04»group\_4 |  | 35 |  | 1597-04»group\_4 |  | 35 |  | 12655-03»   group\_4 |
| 36 |  | 4523-03»group\_4 |  | 36 |  | 4523-03»group\_4 |  | 36 |  | 3734-04»group\_4 |
| 37 |  | 5248-04»group\_5 |  | 37 |  | 5248-04»group\_5 |  | 37 |  | 5248-04»group\_5 |
| 38 |  | 7517-04»group\_5 |  | 38 |  | 7517-04»group\_5 |  | 38 |  | 7517-04»group\_5 |
| 39 |  | 7538-03»group\_5 |  | 39 |  | 7538-03»group\_5 |  | 39 |  | 7538-03»group\_5 |
| 40 |  | 11818-03»   group\_6 |  | 40 |  | 11818-03»   group\_6 |  | 40 |  | 11818-03»   group\_6 |
| 41 |  | 4781-04»group\_6 |  | 41 |  | 4781-04»group\_6 |  | 41 |  | 4781-04»group\_6 |
| 42 |  | 10348-03»   group\_7 |  | 42 |  | 10348-03»   group\_7 |  | 42 |  | 10348-03»   group\_7 |
| 43 |  | 6639-04»group\_7 |  | 43 |  | 6639-04»group\_7 |  | 43 |  | 6639-04»group\_7 |
| 44 |  | 12658-03»   group\_8 |  | 44 |  | 12658-03»   group\_8 |  | 44 |  | 12658-03»   group\_8 |
| 45 |  | 7135-04»group\_8 |  | 45 |  | 7135-04»group\_8 |  | 45 |  | 7135-04»group\_8 |
| 46 |  | 11822-03»   group\_9 |  | 46 |  | 11822-03»   group\_9 |  | 46 |  | 11822-03»   group\_9 |
| 47 |  | 1599-04»group\_9 |  | 47 |  | 1599-04»group\_9 |  | 47 |  | 1599-04»group\_9 |
| 48 |  | 4148-04»group\_9 |  | 48 |  | 4148-04»group\_9 |  | 48 |  | 4148-04»group\_9 |
| 49 |  | 4514-03»group\_9 |  | 49 |  | 4514-03»group\_9 |  | 49 |  | 4514-03»group\_9 |
| 50 |  | 6467-04»group\_9 |  | 50 |  | 6467-04»group\_9 |  | 50 |  | 6467-04»group\_9 |
| 51 |  | 10010-03»   ungrouped |  | 51 |  | 10010-03»   ungrouped |  | 51 |  | 10010-03»   ungrouped |
| 52 |  | 10011-03»   ungrouped |  | 52 |  | 10011-03»   ungrouped |  | 52 |  | 10011-03»   ungrouped |
| 53 |  | 10012-03»   ungrouped |  | 53 |  | 10012-03»   ungrouped |  | 53 |  | 10012-03»   ungrouped |
| 54 |  | 10205-03»   ungrouped |  | 54 |  | 10205-03»   ungrouped |  | 54 |  | 10205-03»   ungrouped |
| 55 |  | 10208-03»   ungrouped |  | 55 |  | 10208-03»   ungrouped |  | 55 |  | 10208-03»   ungrouped |
| 56 |  | 10349-03»   ungrouped |  | 56 |  | 10349-03»   ungrouped |  | 56 |  | 10349-03»   ungrouped |
| 57 |  | 10350-03»   ungrouped |  | 57 |  | 10350-03»   ungrouped |  | 57 |  | 10350-03»   ungrouped |
| 58 |  | 11096-03»   ungrouped |  | 58 |  | 11096-03»   ungrouped |  | 58 |  | 11096-03»   ungrouped |
| 59 |  | 11097-03»   ungrouped |  | 59 |  | 11097-03»   ungrouped |  | 59 |  | 11097-03»   ungrouped |
| 60 |  | 11821-03»   ungrouped |  | 60 |  | 11821-03»   ungrouped |  | 60 |  | 11821-03»   ungrouped |
| 61 |  | 12657-03»   ungrouped |  | 61 |  | 12657-03»   ungrouped |  | 61 |  | 12657-03»   ungrouped |
| 62 |  | 1322-04»ungrouped |  | 62 |  | 1322-04»ungrouped |  | 62 |  | 1322-04»ungrouped |
| 63 |  | 1324-04»ungrouped |  | 63 |  | 1324-04»ungrouped |  | 63 |  | 1324-04»ungrouped |
| 64 |  | 1327-04»ungrouped |  | 64 |  | 1327-04»ungrouped |  | 64 |  | 1327-04»ungrouped |
| 65 |  | 1783-04»ungrouped |  | 65 |  | 1783-04»ungrouped |  | 65 |  | 1783-04»ungrouped |
| 66 |  | 2509-04»ungrouped |  | 66 |  | 2509-04»ungrouped |  | 66 |  | 2509-04»ungrouped |
| 67 |  | 3154-04»ungrouped |  | 67 |  | 3154-04»ungrouped |  | 67 |  | 3154-04»ungrouped |
| 68 |  | 3156-04»ungrouped |  | 68 |  | 3156-04»ungrouped |  | 68 |  | 3156-04»ungrouped |
| 69 |  | 3158-04»ungrouped |  | 69 |  | 3158-04»ungrouped |  | 69 |  | 3158-04»ungrouped |
| 70 |  | 3160-04»ungrouped |  | 70 |  | 3160-04»ungrouped |  | 70 |  | 3160-04»ungrouped |
| 71 |  | 3491-04»ungrouped |  | 71 |  | 3491-04»ungrouped |  | 71 |  | 3491-04»ungrouped |
| 72 |  | 3494-04»ungrouped |  | 72 |  | 3494-04»ungrouped |  | 72 |  | 3494-04»ungrouped |
| 73 |  | 3496-04»ungrouped |  | 73 |  | 3496-04»ungrouped |  | 73 |  | 3496-04»ungrouped |
| 74 |  | 3497-04»ungrouped |  | 74 |  | 3497-04»ungrouped |  | 74 |  | 3497-04»ungrouped |
| 75 |  | 3736-04»ungrouped |  | 75 |  | 3736-04»ungrouped |  | 75 |  | 3736-04»ungrouped |
| 76 |  | 3859-03»ungrouped |  | 76 |  | 3859-03»ungrouped |  | 76 |  | 3859-03»ungrouped |
| 77 |  | 3861-03»ungrouped |  | 77 |  | 3861-03»ungrouped |  | 77 |  | 3861-03»ungrouped |
| 78 |  | 4139-04»ungrouped |  | 78 |  | 4139-04»ungrouped |  | 78 |  | 4139-04»ungrouped |
| 79 |  | 4145-04»ungrouped |  | 79 |  | 4145-04»ungrouped |  | 79 |  | 4145-04»ungrouped |
| 80 |  | 420-04» ungrouped |  | 80 |  | 420-04» ungrouped |  | 80 |  | 420-04» ungrouped |
| 81 |  | 421-04» ungrouped |  | 81 |  | 421-04» ungrouped |  | 81 |  | 421-04» ungrouped |
| 82 |  | 4516-03»ungrouped |  | 82 |  | 4516-03»ungrouped |  | 82 |  | 4516-03»ungrouped |
| 83 |  | 4518-03»ungrouped |  | 83 |  | 4518-03»ungrouped |  | 83 |  | 4518-03»ungrouped |
| 84 |  | 4712-04»ungrouped |  | 84 |  | 4712-04»ungrouped |  | 84 |  | 4712-04»ungrouped |
| 85 |  | 4714-04»ungrouped |  | 85 |  | 4714-04»ungrouped |  | 85 |  | 4714-04»ungrouped |
| 86 |  | 4717-04»ungrouped |  | 86 |  | 4717-04»ungrouped |  | 86 |  | 4717-04»ungrouped |
| 87 |  | 4724-03»ungrouped |  | 87 |  | 4724-03»ungrouped |  | 87 |  | 4724-03»ungrouped |
| 88 |  | 4779-04»ungrouped |  | 88 |  | 4779-04»ungrouped |  | 88 |  | 4779-04»ungrouped |
| 89 |  | 4783-04»ungrouped |  | 89 |  | 4783-04»ungrouped |  | 89 |  | 4783-04»ungrouped |
| 90 |  | 4785-04»ungrouped |  | 90 |  | 4785-04»ungrouped |  | 90 |  | 4785-04»ungrouped |
| 91 |  | 5253-04»ungrouped |  | 91 |  | 5253-04»ungrouped |  | 91 |  | 5253-04»ungrouped |
| 92 |  | 5468-03»ungrouped |  | 92 |  | 5468-03»ungrouped |  | 92 |  | 5468-03»ungrouped |
| 93 |  | 5472-03»ungrouped |  | 93 |  | 5472-03»ungrouped |  | 93 |  | 5472-03»ungrouped |
| 94 |  | 5687-04»ungrouped |  | 94 |  | 5687-04»ungrouped |  | 94 |  | 5687-04»ungrouped |
| 95 |  | 5870-03»ungrouped |  | 95 |  | 5870-03»ungrouped |  | 95 |  | 5870-03»ungrouped |
| 96 |  | 5872-03»ungrouped |  | 96 |  | 5872-03»ungrouped |  | 96 |  | 5872-03»ungrouped |
| 97 |  | 6429-03»ungrouped |  | 97 |  | 6429-03»ungrouped |  | 97 |  | 6429-03»ungrouped |
| 98 |  | 6435-03»ungrouped |  | 98 |  | 6435-03»ungrouped |  | 98 |  | 6435-03»ungrouped |
| 99 |  | 6637-04»ungrouped |  | 99 |  | 6637-04»ungrouped |  | 99 |  | 6637-04»ungrouped |
| 100 |  | 6640-04»ungrouped |  | 100 |  | 6640-04»ungrouped |  | 100 |  | 6640-04»ungrouped |
| 101 |  | 6769-04»ungrouped |  | 101 |  | 6769-04»ungrouped |  | 101 |  | 6769-04»ungrouped |
| 102 |  | 6771-04»ungrouped |  | 102 |  | 6771-04»ungrouped |  | 102 |  | 6771-04»ungrouped |
| 103 |  | 6775-04»ungrouped |  | 103 |  | 6775-04»ungrouped |  | 103 |  | 6775-04»ungrouped |
| 104 |  | 6892-04»ungrouped |  | 104 |  | 6892-04»ungrouped |  | 104 |  | 6892-04»ungrouped |
| 105 |  | 6895-04»ungrouped |  | 105 |  | 6895-04»ungrouped |  | 105 |  | 6895-04»ungrouped |
| 106 |  | 6897-04»ungrouped |  | 106 |  | 6897-04»ungrouped |  | 106 |  | 6897-04»ungrouped |
| 107 |  | 7000-03»ungrouped |  | 107 |  | 7000-03»ungrouped |  | 107 |  | 7000-03»ungrouped |
| 108 |  | 7514-04»ungrouped |  | 108 |  | 7514-04»ungrouped |  | 108 |  | 7514-04»ungrouped |
| 109 |  | 7516-04»ungrouped |  | 109 |  | 7516-04»ungrouped |  | 109 |  | 7516-04»ungrouped |
| 110 |  | 7520-04»ungrouped |  | 110 |  | 7520-04»ungrouped |  | 110 |  | 7520-04»ungrouped |
| 111 |  | 8082-03»ungrouped |  | 111 |  | 8082-03»ungrouped |  | 111 |  | 8082-03»ungrouped |
| 112 |  | 8864-03»ungrouped |  | 112 |  | 8864-03»ungrouped |  | 112 |  | 8864-03»ungrouped |
| 113 |  | 8867-03»ungrouped |  | 113 |  | 8867-03»ungrouped |  | 113 |  | 8867-03»ungrouped |
| 114 |  | 8868-03»ungrouped |  | 114 |  | 8868-03»ungrouped |  | 114 |  | 8868-03»ungrouped |

Araxis Merge (but not the data content of this report) is Copyright © 1993–2024 Araxis Ltd (www.araxis.com). All rights reserved.
