## Supplementary material for "MTBseq-nf: Enabling Scalable Tuberculosis Genomics “Big Data” Analysis through a User-Friendly Nextflow Wrapper for MTBseq pipeline": SD-6 Intra-modal analysis, with 3-way HTML diff reports generated by Araxis merge software: SD-6-03-intra-modal-araxiscompare-pub-90samples-mtbseq-standard-runs-snp-matrix.html

xml version="1.0" encoding="utf-8"?


Araxis Merge File Comparison Report 

### **Araxis Merge File Comparison Report**

Produced by **Araxis Merge** on **2024/12/16, 16:26 GMT+02:00**. See www.araxis.com for information about Merge. This report uses XHTML and CSS2, and is best viewed with a modern standards-compliant browser. For optimum results when printing this report, use landscape orientation and enable printing of background images and colours in your browser.

##### **1. Files compared**

| # | Location | File | Last Modified |
| --- | --- | --- | --- |
| 1 | /Users/abhi/projects/MTBseq-nf/\_resources/publication/manuscript-and-analysis/v2/pub-90samples-mtbseq-standard-run1/Groups | mtbseqnf\_joint\_cf4\_cr4\_fr75\_ph4\_samples90\_amended\_u95\_phylo\_w12.matrix | 2024/10/15, 17:41 GMT+02:00 |
| 2 | /Users/abhi/projects/MTBseq-nf/\_resources/publication/manuscript-and-analysis/v2/pub-90samples-mtbseq-standard-run2/Groups | mtbseqnf\_joint\_cf4\_cr4\_fr75\_ph4\_samples90\_amended\_u95\_phylo\_w12.matrix | 2024/10/16, 05:25 GMT+02:00 |
| 3 | /Users/abhi/projects/MTBseq-nf/\_resources/publication/manuscript-and-analysis/v2/pub-90samples-mtbseq-standard-run3/Groups | mtbseqnf\_joint\_cf4\_cr4\_fr75\_ph4\_samples90\_amended\_u95\_phylo\_w12.matrix | 2024/10/16, 20:21 GMT+02:00 |
| **Note:** Merge considers the second file to be the common ancestor of the others. | | | |

##### **2. Comparison summary**

| Description | Between Files 1 and 2 | | Between Files 2 and 3 | | Relative to Common Ancestor | |
| --- | --- | --- | --- | --- | --- | --- |
|  | Text Blocks | Lines | Text Blocks | Lines | Text Blocks | Lines |
| Unchanged | 1 | 182 | 1 | 182 |  |  |
| Changed | 0 | 0 | 0 | 0 | 0 | 0 |
| Inserted | 0 | 0 | 0 | 0 | 0 | 0 |
| Removed | 0 | 0 | 0 | 0 | 0 | 0 |
| **Note:** An automatic merge would leave 0 conflict(s). | | | | | | |

##### **3. Comparison options**

|  |  |
| --- | --- |
| Whitespace | Consecutive whitespace is treated as a single space |
| Character case | Differences in character case are significant |
| Line endings | Differences in line endings (`CR` and `LF` characters) are ignored |
| CR/LF characters | Not shown in the comparison detail |
| Active line-pairing rules |  |

##### **4. Active regular expressions**

No regular expressions were active.

##### **5. Comparison detail**

|  |  |  |  |  |  |  |  |  |  |  |
| --- | --- | --- | --- | --- | --- | --- | --- | --- | --- | --- |
| 1 |  | 10010-03\_lib951» |  | 1 |  | 10010-03\_lib951» |  | 1 |  | 10010-03\_lib951» |
| 2 |  | 10011-03\_lib914»1484» |  | 2 |  | 10011-03\_lib914»1484» |  | 2 |  | 10011-03\_lib914»1484» |
| 3 |  | 10012-03\_lib970»1449»   665» |  | 3 |  | 10012-03\_lib970»1449»   665» |  | 3 |  | 10012-03\_lib970»1449»   665» |
| 4 |  | 10205-03\_lib915»1418»   214»601» |  | 4 |  | 10205-03\_lib915»1418»   214»601» |  | 4 |  | 10205-03\_lib915»1418»   214»601» |
| 5 |  | 10206-03\_lib916»1477»   33» 658»207» |  | 5 |  | 10206-03\_lib916»1477»   33» 658»207» |  | 5 |  | 10206-03\_lib916»1477»   33» 658»207» |
| 6 |  | 10207-03\_lib973»1459»   589»640»527»582» |  | 6 |  | 10207-03\_lib973»1459»   589»640»527»582» |  | 6 |  | 10207-03\_lib973»1459»   589»640»527»582» |
| 7 |  | 10208-03\_lib1613»   1577»   1637»   1598»   1571»   1630»   1610» |  | 7 |  | 10208-03\_lib1613»   1577»   1637»   1598»   1571»   1630»   1610» |  | 7 |  | 10208-03\_lib1613»   1577»   1637»   1598»   1571»   1630»   1610» |
| 8 |  | 10348-03\_lib917»1450»   366»633»304»359»555»1601» |  | 8 |  | 10348-03\_lib917»1450»   366»633»304»359»555»1601» |  | 8 |  | 10348-03\_lib917»1450»   366»633»304»359»555»1601» |
| 9 |  | 10349-03\_lib924»1430»   646»425»584»639»619»1579»   612» |  | 9 |  | 10349-03\_lib924»1430»   646»425»584»639»619»1579»   612» |  | 9 |  | 10349-03\_lib924»1430»   646»425»584»639»619»1579»   612» |
| 10 |  | 10350-03\_lib1470»   1432»   648»477»584»641»621»1581»   614»456» |  | 10 |  | 10350-03\_lib1470»   1432»   648»477»584»641»621»1581»   614»456» |  | 10 |  | 10350-03\_lib1470»   1432»   648»477»584»641»621»1581»   614»456» |
| 11 |  | 10517-03\_lib939»1445»   575»628»513»568»512»1598»   541»607»609» |  | 11 |  | 10517-03\_lib939»1445»   575»628»513»568»512»1598»   541»607»609» |  | 11 |  | 10517-03\_lib939»1445»   575»628»513»568»512»1598»   541»607»609» |
| 12 |  | 11096-03\_lib940»1441»   573»624»509»566»510»1594»   539»605»605»44» |  | 12 |  | 11096-03\_lib940»1441»   573»624»509»566»510»1594»   539»605»605»44» |  | 12 |  | 11096-03\_lib940»1441»   573»624»509»566»510»1594»   539»605»605»44» |
| 13 |  | 11097-03\_lib933»1468»   686»139»622»679»659»1617»   652»444»496»647»643» |  | 13 |  | 11097-03\_lib933»1468»   686»139»622»679»659»1617»   652»444»496»647»643» |  | 13 |  | 11097-03\_lib933»1468»   686»139»622»679»659»1617»   652»444»496»647»643» |
| 14 |  | 11818-03\_lib902»1415»   631»460»567»624»604»1564»   597»439»193»592»588»479» |  | 14 |  | 11818-03\_lib902»1415»   631»460»567»624»604»1564»   597»439»193»592»588»479» |  | 14 |  | 11818-03\_lib902»1415»   631»460»567»624»604»1564»   597»439»193»592»588»479» |
| 15 |  | 11821-03\_lib952»336»1532»   1497»   1466»   1525»   1505»   1625»   1498»   1478»   1480»   1493»   1489»   1516»   1463» |  | 15 |  | 11821-03\_lib952»336»1532»   1497»   1466»   1525»   1505»   1625»   1498»   1478»   1480»   1493»   1489»   1516»   1463» |  | 15 |  | 11821-03\_lib952»336»1532»   1497»   1466»   1525»   1505»   1625»   1498»   1478»   1480»   1493»   1489»   1516»   1463» |
| 16 |  | 11822-03\_lib903»1413»   629»456»565»622»602»1562»   595»435»181»590»586»475»174»1461» |  | 16 |  | 11822-03\_lib903»1413»   629»456»565»622»602»1562»   595»435»181»590»586»475»174»1461» |  | 16 |  | 11822-03\_lib903»1413»   629»456»565»622»602»1562»   595»435»181»590»586»475»174»1461» |
| 17 |  | 12655-03\_lib1516»   1459»   589»640»527»582»4»  1610»   555»619»621»512»510»659»604»1505»   602» |  | 17 |  | 12655-03\_lib1516»   1459»   589»640»527»582»4»  1610»   555»619»621»512»510»659»604»1505»   602» |  | 17 |  | 12655-03\_lib1516»   1459»   589»640»527»582»4»  1610»   555»619»621»512»510»659»604»1505»   602» |
| 18 |  | 12657-03\_lib934»1458»   676»117»612»669»649»1607»   640»434»486»637»633»148»469»1506»   465»649» |  | 18 |  | 12657-03\_lib934»1458»   676»117»612»669»649»1607»   640»434»486»637»633»148»469»1506»   465»649» |  | 18 |  | 12657-03\_lib934»1458»   676»117»612»669»649»1607»   640»434»486»637»633»148»469»1506»   465»649» |
| 19 |  | 12658-03\_lib1469»   1475»   941»906»879»934»918»1626»   907»887»891»904»902»927»874»1523»   872»918»917» |  | 19 |  | 12658-03\_lib1469»   1475»   941»906»879»934»918»1626»   907»887»891»904»902»927»874»1523»   872»918»917» |  | 19 |  | 12658-03\_lib1469»   1475»   941»906»879»934»918»1626»   907»887»891»904»902»927»874»1523»   872»918»917» |
| 20 |  | 1322-04\_lib925» 1471»   603»650»539»596»538»1620»   567»633»633»310»306»671»616»1519»   614»538»661»932» |  | 20 |  | 1322-04\_lib925» 1471»   603»650»539»596»538»1620»   567»633»633»310»306»671»616»1519»   614»538»661»932» |  | 20 |  | 1322-04\_lib925» 1471»   603»650»539»596»538»1620»   567»633»633»310»306»671»616»1519»   614»538»661»932» |
| 21 |  | 1324-04\_lib931» 1424»   554»603»490»546»527»1573»   518»582»582»515»511»622»565»1472»   563»527»612»883»539» |  | 21 |  | 1324-04\_lib931» 1424»   554»603»490»546»527»1573»   518»582»582»515»511»622»565»1472»   563»527»612»883»539» |  | 21 |  | 1324-04\_lib931» 1424»   554»603»490»546»527»1573»   518»582»582»515»511»622»565»1472»   563»527»612»883»539» |
| 22 |  | 1327-04\_lib957» 1544»   1600»   1563»   1536»   1593»   1575»   681»1568»   1544»   1546»   1563»   1559»   1582»   1529»   1592»   1527»   1575»   1572»   1590»   1587»   1538» |  | 22 |  | 1327-04\_lib957» 1544»   1600»   1563»   1536»   1593»   1575»   681»1568»   1544»   1546»   1563»   1559»   1582»   1529»   1592»   1527»   1575»   1572»   1590»   1587»   1538» |  | 22 |  | 1327-04\_lib957» 1544»   1600»   1563»   1536»   1593»   1575»   681»1568»   1544»   1546»   1563»   1559»   1582»   1529»   1592»   1527»   1575»   1572»   1590»   1587»   1538» |
| 23 |  | 1597-04\_lib918» 1439»   233»618»141»226»546»1590»   323»601»603»532»528»639»586»1487»   582»546»629»896»558»509»1555» |  | 23 |  | 1597-04\_lib918» 1439»   233»618»141»226»546»1590»   323»601»603»532»528»639»586»1487»   582»546»629»896»558»509»1555» |  | 23 |  | 1597-04\_lib918» 1439»   233»618»141»226»546»1590»   323»601»603»532»528»639»586»1487»   582»546»629»896»558»509»1555» |
| 24 |  | 1599-04\_lib974» 1414»   630»457»566»623»603»1563»   596»436»182»591»587»476»175»1462»   11» 603»464»873»615»564»1528»   583» |  | 24 |  | 1599-04\_lib974» 1414»   630»457»566»623»603»1563»   596»436»182»591»587»476»175»1462»   11» 603»464»873»615»564»1528»   583» |  | 24 |  | 1599-04\_lib974» 1414»   630»457»566»623»603»1563»   596»436»182»591»587»476»175»1462»   11» 603»464»873»615»564»1528»   583» |
| 25 |  | 1779-04\_lib941» 1442»   572»625»510»565»509»1595»   538»604»606»7»  41» 644»589»1490»   587»509»634»901»307»512»1560»   529»588» |  | 25 |  | 1779-04\_lib941» 1442»   572»625»510»565»509»1595»   538»604»606»7»  41» 644»589»1490»   587»509»634»901»307»512»1560»   529»588» |  | 25 |  | 1779-04\_lib941» 1442»   572»625»510»565»509»1595»   538»604»606»7»  41» 644»589»1490»   587»509»634»901»307»512»1560»   529»588» |
| 26 |  | 1780-04\_lib942» 1442»   572»625»510»565»509»1595»   538»604»606»7»  41» 644»589»1490»   587»509»634»901»307»512»1560»   529»588»0» |  | 26 |  | 1780-04\_lib942» 1442»   572»625»510»565»509»1595»   538»604»606»7»  41» 644»589»1490»   587»509»634»901»307»512»1560»   529»588»0» |  | 26 |  | 1780-04\_lib942» 1442»   572»625»510»565»509»1595»   538»604»606»7»  41» 644»589»1490»   587»509»634»901»307»512»1560»   529»588»0» |
| 27 |  | 1783-04\_lib936» 1459»   677»104»613»670»650»1608»   643»435»487»638»634»149»470»1507»   466»650»127»916»662»613»1573»   630»467»635»635» |  | 27 |  | 1783-04\_lib936» 1459»   677»104»613»670»650»1608»   643»435»487»638»634»149»470»1507»   466»650»127»916»662»613»1573»   630»467»635»635» |  | 27 |  | 1783-04\_lib936» 1459»   677»104»613»670»650»1608»   643»435»487»638»634»149»470»1507»   466»650»127»916»662»613»1573»   630»467»635»635» |
| 28 |  | 2509-04\_lib1471»1479»   35» 660»209»28» 584»1632»   361»641»643»570»568»681»626»1527»   624»584»671»936»598»549»1595»   228»625»567»567»672» |  | 28 |  | 2509-04\_lib1471»1479»   35» 660»209»28» 584»1632»   361»641»643»570»568»681»626»1527»   624»584»671»936»598»549»1595»   228»625»567»567»672» |  | 28 |  | 2509-04\_lib1471»1479»   35» 660»209»28» 584»1632»   361»641»643»570»568»681»626»1527»   624»584»671»936»598»549»1595»   228»625»567»567»672» |
| 29 |  | 3154-04\_lib905» 1416»   634»461»570»627»607»1567»   600»440»186»595»591»480»179»1464»   45» 607»470»877»619»568»1532»   587»46» 592»592»471»629» |  | 29 |  | 3154-04\_lib905» 1416»   634»461»570»627»607»1567»   600»440»186»595»591»480»179»1464»   45» 607»470»877»619»568»1532»   587»46» 592»592»471»629» |  | 29 |  | 3154-04\_lib905» 1416»   634»461»570»627»607»1567»   600»440»186»595»591»480»179»1464»   45» 607»470»877»619»568»1532»   587»46» 592»592»471»629» |
| 30 |  | 3156-04\_lib926» 1439»   655»484»591»648»626»1586»   621»465»463»616»612»503»446»1487»   444»628»493»896»640»588»1549»   608»445»613»613»494»650»449» |  | 30 |  | 3156-04\_lib926» 1439»   655»484»591»648»626»1586»   621»465»463»616»612»503»446»1487»   444»628»493»896»640»588»1549»   608»445»613»613»494»650»449» |  | 30 |  | 3156-04\_lib926» 1439»   655»484»591»648»626»1586»   621»465»463»616»612»503»446»1487»   444»628»493»896»640»588»1549»   608»445»613»613»494»650»449» |
| 31 |  | 3158-04\_lib943» 1445»   575»628»513»568»512»1598»   541»607»609»16» 44» 647»592»1493»   590»512»637»904»310»515»1563»   532»591»13» 13» 638»570»595»616» |  | 31 |  | 3158-04\_lib943» 1445»   575»628»513»568»512»1598»   541»607»609»16» 44» 647»592»1493»   590»512»637»904»310»515»1563»   532»591»13» 13» 638»570»595»616» |  | 31 |  | 3158-04\_lib943» 1445»   575»628»513»568»512»1598»   541»607»609»16» 44» 647»592»1493»   590»512»637»904»310»515»1563»   532»591»13» 13» 638»570»595»616» |
| 32 |  | 3160-04\_lib958» 1537»   1591»   1554»   1527»   1584»   1566»   674»1559»   1535»   1537»   1554»   1550»   1573»   1520»   1585»   1518»   1566»   1563»   1581»   1578»   1529»   105»1546»   1519»   1551»   1551»   1564»   1586»   1523»   1542»   1554» |  | 32 |  | 3160-04\_lib958» 1537»   1591»   1554»   1527»   1584»   1566»   674»1559»   1535»   1537»   1554»   1550»   1573»   1520»   1585»   1518»   1566»   1563»   1581»   1578»   1529»   105»1546»   1519»   1551»   1551»   1564»   1586»   1523»   1542»   1554» |  | 32 |  | 3160-04\_lib958» 1537»   1591»   1554»   1527»   1584»   1566»   674»1559»   1535»   1537»   1554»   1550»   1573»   1520»   1585»   1518»   1566»   1563»   1581»   1578»   1529»   105»1546»   1519»   1551»   1551»   1564»   1586»   1523»   1542»   1554» |
| 33 |  | 3491-04\_lib897» 1555»   1489»   1454»   1427»   1482»   1466»   1708»   1457»   1435»   1437»   1451»   1447»   1473»   1420»   1603»   1418»   1466»   1463»   1484»   1477»   1429»   1671»   1446»   1419»   1448»   1448»   1464»   1484»   1423»   1444»   1451»   1662» |  | 33 |  | 3491-04\_lib897» 1555»   1489»   1454»   1427»   1482»   1466»   1708»   1457»   1435»   1437»   1451»   1447»   1473»   1420»   1603»   1418»   1466»   1463»   1484»   1477»   1429»   1671»   1446»   1419»   1448»   1448»   1464»   1484»   1423»   1444»   1451»   1662» |  | 33 |  | 3491-04\_lib897» 1555»   1489»   1454»   1427»   1482»   1466»   1708»   1457»   1435»   1437»   1451»   1447»   1473»   1420»   1603»   1418»   1466»   1463»   1484»   1477»   1429»   1671»   1446»   1419»   1448»   1448»   1464»   1484»   1423»   1444»   1451»   1662» |
| 34 |  | 3494-04\_lib953» 332»1528»   1493»   1462»   1521»   1501»   1621»   1494»   1474»   1476»   1489»   1485»   1512»   1459»   78» 1457»   1501»   1502»   1519»   1515»   1468»   1588»   1483»   1458»   1486»   1486»   1503»   1523»   1460»   1483»   1489»   1581»   1599» |  | 34 |  | 3494-04\_lib953» 332»1528»   1493»   1462»   1521»   1501»   1621»   1494»   1474»   1476»   1489»   1485»   1512»   1459»   78» 1457»   1501»   1502»   1519»   1515»   1468»   1588»   1483»   1458»   1486»   1486»   1503»   1523»   1460»   1483»   1489»   1581»   1599» |  | 34 |  | 3494-04\_lib953» 332»1528»   1493»   1462»   1521»   1501»   1621»   1494»   1474»   1476»   1489»   1485»   1512»   1459»   78» 1457»   1501»   1502»   1519»   1515»   1468»   1588»   1483»   1458»   1486»   1486»   1503»   1523»   1460»   1483»   1489»   1581»   1599» |
| 35 |  | 3496-04\_lib906» 1414»   630»455»566»623»603»1563»   596»436»182»591»587»474»175»1462»   41» 603»464»873»615»564»1528»   583»42» 588»588»465»625»38» 445»591»1519»   1419»   1458» |  | 35 |  | 3496-04\_lib906» 1414»   630»455»566»623»603»1563»   596»436»182»591»587»474»175»1462»   41» 603»464»873»615»564»1528»   583»42» 588»588»465»625»38» 445»591»1519»   1419»   1458» |  | 35 |  | 3496-04\_lib906» 1414»   630»455»566»623»603»1563»   596»436»182»591»587»474»175»1462»   41» 603»464»873»615»564»1528»   583»42» 588»588»465»625»38» 445»591»1519»   1419»   1458» |
| 36 |  | 3497-04\_lib927» 1433»   565»614»501»558»500»1584»   531»595»595»272»268»633»578»1481»   576»500»623»894»188»501»1549»   520»577»269»269»624»560»581»602»272»1540»   1439»   1477»   577» |  | 36 |  | 3497-04\_lib927» 1433»   565»614»501»558»500»1584»   531»595»595»272»268»633»578»1481»   576»500»623»894»188»501»1549»   520»577»269»269»624»560»581»602»272»1540»   1439»   1477»   577» |  | 36 |  | 3497-04\_lib927» 1433»   565»614»501»558»500»1584»   531»595»595»272»268»633»578»1481»   576»500»623»894»188»501»1549»   520»577»269»269»624»560»581»602»272»1540»   1439»   1477»   577» |
| 37 |  | 3734-04\_lib895» 1458»   588»639»526»581»3»  1609»   554»618»620»511»509»658»603»1504»   601»1»  648»917»537»526»1574»   545»602»508»508»649»583»606»627»511»1565»   1465»   1500»   602»499» |  | 37 |  | 3734-04\_lib895» 1458»   588»639»526»581»3»  1609»   554»618»620»511»509»658»603»1504»   601»1»  648»917»537»526»1574»   545»602»508»508»649»583»606»627»511»1565»   1465»   1500»   602»499» |  | 37 |  | 3734-04\_lib895» 1458»   588»639»526»581»3»  1609»   554»618»620»511»509»658»603»1504»   601»1»  648»917»537»526»1574»   545»602»508»508»649»583»606»627»511»1565»   1465»   1500»   602»499» |
| 38 |  | 3736-04\_lib937» 1455»   673»118»609»666»648»1608»   639»433»485»636»632»149»468»1503»   464»648»115»914»660»609»1573»   626»465»633»633»128»668»469»494»636»1564»   1462»   1499»   463»622»647» |  | 38 |  | 3736-04\_lib937» 1455»   673»118»609»666»648»1608»   639»433»485»636»632»149»468»1503»   464»648»115»914»660»609»1573»   626»465»633»633»128»668»469»494»636»1564»   1462»   1499»   463»622»647» |  | 38 |  | 3736-04\_lib937» 1455»   673»118»609»666»648»1608»   639»433»485»636»632»149»468»1503»   464»648»115»914»660»609»1573»   626»465»633»633»128»668»469»494»636»1564»   1462»   1499»   463»622»647» |
| 39 |  | 3859-03\_lib921» 1471»   687»202»623»680»660»1620»   653»445»497»648»644»221»480»1519»   475»660»211»930»672»623»1585»   640»476»645»645»212»682»480»504»648»1576»   1476»   1515»   476»634»659»212» |  | 39 |  | 3859-03\_lib921» 1471»   687»202»623»680»660»1620»   653»445»497»648»644»221»480»1519»   475»660»211»930»672»623»1585»   640»476»645»645»212»682»480»504»648»1576»   1476»   1515»   476»634»659»212» |  | 39 |  | 3859-03\_lib921» 1471»   687»202»623»680»660»1620»   653»445»497»648»644»221»480»1519»   475»660»211»930»672»623»1585»   640»476»645»645»212»682»480»504»648»1576»   1476»   1515»   476»634»659»212» |
| 40 |  | 3861-03\_lib922» 1442»   578»625»512»571»497»1593»   542»604»606»501»497»644»589»1489»   585»497»634»907»523»512»1560»   531»586»498»498»635»573»590»615»501»1551»   1453»   1485»   586»487»496»631»645» |  | 40 |  | 3861-03\_lib922» 1442»   578»625»512»571»497»1593»   542»604»606»501»497»644»589»1489»   585»497»634»907»523»512»1560»   531»586»498»498»635»573»590»615»501»1551»   1453»   1485»   586»487»496»631»645» |  | 40 |  | 3861-03\_lib922» 1442»   578»625»512»571»497»1593»   542»604»606»501»497»644»589»1489»   585»497»634»907»523»512»1560»   531»586»498»498»635»573»590»615»501»1551»   1453»   1485»   586»487»496»631»645» |
| 41 |  | 3865-03\_lib971» 1480»   36» 661»210»11» 585»1633»   362»642»644»571»569»682»627»1528»   625»585»672»937»599»549»1596»   229»626»568»568»673»31» 630»651»571»1587»   1485»   1524»   626»561»584»669»683»574» |  | 41 |  | 3865-03\_lib971» 1480»   36» 661»210»11» 585»1633»   362»642»644»571»569»682»627»1528»   625»585»672»937»599»549»1596»   229»626»568»568»673»31» 630»651»571»1587»   1485»   1524»   626»561»584»669»683»574» |  | 41 |  | 3865-03\_lib971» 1480»   36» 661»210»11» 585»1633»   362»642»644»571»569»682»627»1528»   625»585»672»937»599»549»1596»   229»626»568»568»673»31» 630»651»571»1587»   1485»   1524»   626»561»584»669»683»574» |
| 42 |  | 4139-04\_lib959» 1553»   1612»   1573»   1546»   1605»   1585»   400»1578»   1554»   1556»   1573»   1569»   1592»   1539»   1601»   1537»   1585»   1582»   1601»   1595»   1548»   656»1565»   1538»   1570»   1570»   1583»   1607»   1542»   1561»   1573»   649»1683»   1597»   1538»   1557»   1584»   1583»   1595»   1570»   1608» |  | 42 |  | 4139-04\_lib959» 1553»   1612»   1573»   1546»   1605»   1585»   400»1578»   1554»   1556»   1573»   1569»   1592»   1539»   1601»   1537»   1585»   1582»   1601»   1595»   1548»   656»1565»   1538»   1570»   1570»   1583»   1607»   1542»   1561»   1573»   649»1683»   1597»   1538»   1557»   1584»   1583»   1595»   1570»   1608» |  | 42 |  | 4139-04\_lib959» 1553»   1612»   1573»   1546»   1605»   1585»   400»1578»   1554»   1556»   1573»   1569»   1592»   1539»   1601»   1537»   1585»   1582»   1601»   1595»   1548»   656»1565»   1538»   1570»   1570»   1583»   1607»   1542»   1561»   1573»   649»1683»   1597»   1538»   1557»   1584»   1583»   1595»   1570»   1608» |
| 43 |  | 4145-04\_lib919» 1434»   230»617»162»223»543»1587»   320»600»600»529»525»638»583»1482»   581»542»628»895»555»506»1551»   185»582»526»526»629»225»586»607»529»1542»   1443»   1478»   582»517»541»625»639»528»226»1560» |  | 43 |  | 4145-04\_lib919» 1434»   230»617»162»223»543»1587»   320»600»600»529»525»638»583»1482»   581»542»628»895»555»506»1551»   185»582»526»526»629»225»586»607»529»1542»   1443»   1478»   582»517»541»625»639»528»226»1560» |  | 43 |  | 4145-04\_lib919» 1434»   230»617»162»223»543»1587»   320»600»600»529»525»638»583»1482»   581»542»628»895»555»506»1551»   185»582»526»526»629»225»586»607»529»1542»   1443»   1478»   582»517»541»625»639»528»226»1560» |
| 44 |  | 4148-04\_lib907» 1414»   630»457»566»623»603»1563»   596»436»182»591»587»476»175»1462»   11» 603»466»873»615»564»1528»   583»12» 588»588»467»625»46» 445»591»1519»   1419»   1458»   42» 577»602»465»476»586»626»1536»   582» |  | 44 |  | 4148-04\_lib907» 1414»   630»457»566»623»603»1563»   596»436»182»591»587»476»175»1462»   11» 603»466»873»615»564»1528»   583»12» 588»588»467»625»46» 445»591»1519»   1419»   1458»   42» 577»602»465»476»586»626»1536»   582» |  | 44 |  | 4148-04\_lib907» 1414»   630»457»566»623»603»1563»   596»436»182»591»587»476»175»1462»   11» 603»466»873»615»564»1528»   583»12» 588»588»467»625»46» 445»591»1519»   1419»   1458»   42» 577»602»465»476»586»626»1536»   582» |
| 45 |  | 420-04\_lib935»  1448»   666»111»602»659»643»1601»   634»428»480»631»627»142»463»1496»   459»643»108»907»655»606»1566»   619»460»628»628»121»661»464»487»631»1557»   1455»   1492»   458»617»642»93» 205»628»662»1576»   618»460» |  | 45 |  | 420-04\_lib935»  1448»   666»111»602»659»643»1601»   634»428»480»631»627»142»463»1496»   459»643»108»907»655»606»1566»   619»460»628»628»121»661»464»487»631»1557»   1455»   1492»   458»617»642»93» 205»628»662»1576»   618»460» |  | 45 |  | 420-04\_lib935»  1448»   666»111»602»659»643»1601»   634»428»480»631»627»142»463»1496»   459»643»108»907»655»606»1566»   619»460»628»628»121»661»464»487»631»1557»   1455»   1492»   458»617»642»93» 205»628»662»1576»   618»460» |
| 46 |  | 421-04\_lib956»  1533»   1589»   1552»   1525»   1582»   1564»   670»1557»   1533»   1535»   1552»   1548»   1571»   1518»   1581»   1516»   1564»   1561»   1579»   1576»   1527»   99» 1544»   1517»   1549»   1549»   1562»   1584»   1521»   1540»   1552»   94» 1660»   1577»   1517»   1538»   1563»   1562»   1574»   1549»   1585»   645»1540»   1517»   1555» |  | 46 |  | 421-04\_lib956»  1533»   1589»   1552»   1525»   1582»   1564»   670»1557»   1533»   1535»   1552»   1548»   1571»   1518»   1581»   1516»   1564»   1561»   1579»   1576»   1527»   99» 1544»   1517»   1549»   1549»   1562»   1584»   1521»   1540»   1552»   94» 1660»   1577»   1517»   1538»   1563»   1562»   1574»   1549»   1585»   645»1540»   1517»   1555» |  | 46 |  | 421-04\_lib956»  1533»   1589»   1552»   1525»   1582»   1564»   670»1557»   1533»   1535»   1552»   1548»   1571»   1518»   1581»   1516»   1564»   1561»   1579»   1576»   1527»   99» 1544»   1517»   1549»   1549»   1562»   1584»   1521»   1540»   1552»   94» 1660»   1577»   1517»   1538»   1563»   1562»   1574»   1549»   1585»   645»1540»   1517»   1555» |
| 47 |  | 4514-03\_lib901» 1410»   626»453»562»619»599»1559»   592»432»178»587»583»472»171»1458»   5»  599»462»869»611»560»1524»   579»8»  584»584»463»621»42» 441»587»1515»   1415»   1454»   38» 573»598»461»472»582»622»1534»   578»8»  456»1513» |  | 47 |  | 4514-03\_lib901» 1410»   626»453»562»619»599»1559»   592»432»178»587»583»472»171»1458»   5»  599»462»869»611»560»1524»   579»8»  584»584»463»621»42» 441»587»1515»   1415»   1454»   38» 573»598»461»472»582»622»1534»   578»8»  456»1513» |  | 47 |  | 4514-03\_lib901» 1410»   626»453»562»619»599»1559»   592»432»178»587»583»472»171»1458»   5»  599»462»869»611»560»1524»   579»8»  584»584»463»621»42» 441»587»1515»   1415»   1454»   38» 573»598»461»472»582»622»1534»   578»8»  456»1513» |
| 48 |  | 4516-03\_lib947» 1427»   643»472»579»636»616»1574»   607»453»451»604»600»491»434»1475»   432»616»481»886»626»578»1539»   598»433»601»601»482»638»437»340»604»1532»   1432»   1471»   433»590»615»482»492»601»639»1551»   595»433»475»1530»   429» |  | 48 |  | 4516-03\_lib947» 1427»   643»472»579»636»616»1574»   607»453»451»604»600»491»434»1475»   432»616»481»886»626»578»1539»   598»433»601»601»482»638»437»340»604»1532»   1432»   1471»   433»590»615»482»492»601»639»1551»   595»433»475»1530»   429» |  | 48 |  | 4516-03\_lib947» 1427»   643»472»579»636»616»1574»   607»453»451»604»600»491»434»1475»   432»616»481»886»626»578»1539»   598»433»601»601»482»638»437»340»604»1532»   1432»   1471»   433»590»615»482»492»601»639»1551»   595»433»475»1530»   429» |
| 49 |  | 4518-03\_lib949» 154»1480»   1445»   1414»   1473»   1457»   1575»   1448»   1428»   1430»   1443»   1439»   1466»   1413»   332»1411»   1457»   1456»   1473»   1469»   1422»   1542»   1435»   1412»   1440»   1440»   1457»   1475»   1414»   1435»   1443»   1535»   1553»   328»1412»   1431»   1456»   1453»   1469»   1440»   1476»   1551»   1430»   1412»   1446»   1531»   1408»   1423» |  | 49 |  | 4518-03\_lib949» 154»1480»   1445»   1414»   1473»   1457»   1575»   1448»   1428»   1430»   1443»   1439»   1466»   1413»   332»1411»   1457»   1456»   1473»   1469»   1422»   1542»   1435»   1412»   1440»   1440»   1457»   1475»   1414»   1435»   1443»   1535»   1553»   328»1412»   1431»   1456»   1453»   1469»   1440»   1476»   1551»   1430»   1412»   1446»   1531»   1408»   1423» |  | 49 |  | 4518-03\_lib949» 154»1480»   1445»   1414»   1473»   1457»   1575»   1448»   1428»   1430»   1443»   1439»   1466»   1413»   332»1411»   1457»   1456»   1473»   1469»   1422»   1542»   1435»   1412»   1440»   1440»   1457»   1475»   1414»   1435»   1443»   1535»   1553»   328»1412»   1431»   1456»   1453»   1469»   1440»   1476»   1551»   1430»   1412»   1446»   1531»   1408»   1423» |
| 50 |  | 4523-03\_lib972» 1440»   234»619»142»227»547»1591»   324»602»604»533»529»640»587»1488»   583»547»630»897»559»510»1556»   1»  584»530»530»631»229»588»609»533»1547»   1447»   1484»   584»521»546»627»641»532»230»1566»   186»584»620»1545»   580»599»1436» |  | 50 |  | 4523-03\_lib972» 1440»   234»619»142»227»547»1591»   324»602»604»533»529»640»587»1488»   583»547»630»897»559»510»1556»   1»  584»530»530»631»229»588»609»533»1547»   1447»   1484»   584»521»546»627»641»532»230»1566»   186»584»620»1545»   580»599»1436» |  | 50 |  | 4523-03\_lib972» 1440»   234»619»142»227»547»1591»   324»602»604»533»529»640»587»1488»   583»547»630»897»559»510»1556»   1»  584»530»530»631»229»588»609»533»1547»   1447»   1484»   584»521»546»627»641»532»230»1566»   186»584»620»1545»   580»599»1436» |
| 51 |  | 4712-04\_lib960» 1533»   1592»   1553»   1526»   1585»   1565»   380»1558»   1534»   1536»   1553»   1549»   1572»   1519»   1581»   1517»   1565»   1562»   1581»   1575»   1528»   636»1545»   1518»   1550»   1550»   1563»   1587»   1522»   1541»   1553»   629»1663»   1577»   1518»   1537»   1564»   1563»   1575»   1550»   1588»   40» 1540»   1518»   1556»   625»1514»   1531»   1531»   1546» |  | 51 |  | 4712-04\_lib960» 1533»   1592»   1553»   1526»   1585»   1565»   380»1558»   1534»   1536»   1553»   1549»   1572»   1519»   1581»   1517»   1565»   1562»   1581»   1575»   1528»   636»1545»   1518»   1550»   1550»   1563»   1587»   1522»   1541»   1553»   629»1663»   1577»   1518»   1537»   1564»   1563»   1575»   1550»   1588»   40» 1540»   1518»   1556»   625»1514»   1531»   1531»   1546» |  | 51 |  | 4712-04\_lib960» 1533»   1592»   1553»   1526»   1585»   1565»   380»1558»   1534»   1536»   1553»   1549»   1572»   1519»   1581»   1517»   1565»   1562»   1581»   1575»   1528»   636»1545»   1518»   1550»   1550»   1563»   1587»   1522»   1541»   1553»   629»1663»   1577»   1518»   1537»   1564»   1563»   1575»   1550»   1588»   40» 1540»   1518»   1556»   625»1514»   1531»   1531»   1546» |
| 52 |  | 4714-04\_lib1472»1451»   581»628»517»574»554»1600»   543»611»611»542»538»647»594»1499»   592»554»635»910»566»219»1565»   536»593»539»539»638»576»597»618»542»1556»   1456»   1495»   591»528»553»636»650»541»577»1575»   533»593»631»1554»   589»606»1449»   537»1555» |  | 52 |  | 4714-04\_lib1472»1451»   581»628»517»574»554»1600»   543»611»611»542»538»647»594»1499»   592»554»635»910»566»219»1565»   536»593»539»539»638»576»597»618»542»1556»   1456»   1495»   591»528»553»636»650»541»577»1575»   533»593»631»1554»   589»606»1449»   537»1555» |  | 52 |  | 4714-04\_lib1472»1451»   581»628»517»574»554»1600»   543»611»611»542»538»647»594»1499»   592»554»635»910»566»219»1565»   536»593»539»539»638»576»597»618»542»1556»   1456»   1495»   591»528»553»636»650»541»577»1575»   533»593»631»1554»   589»606»1449»   537»1555» |
| 53 |  | 4717-04\_lib898» 1574»   1508»   1473»   1446»   1501»   1485»   1727»   1476»   1454»   1456»   1470»   1466»   1492»   1439»   1622»   1437»   1485»   1482»   1503»   1496»   1448»   1690»   1465»   1438»   1467»   1467»   1483»   1503»   1442»   1461»   1470»   1681»   195»1618»   1438»   1458»   1484»   1481»   1495»   1472»   1504»   1702»   1462»   1438»   1474»   1679»   1434»   1451»   1572»   1466»   1682»   1475» |  | 53 |  | 4717-04\_lib898» 1574»   1508»   1473»   1446»   1501»   1485»   1727»   1476»   1454»   1456»   1470»   1466»   1492»   1439»   1622»   1437»   1485»   1482»   1503»   1496»   1448»   1690»   1465»   1438»   1467»   1467»   1483»   1503»   1442»   1461»   1470»   1681»   195»1618»   1438»   1458»   1484»   1481»   1495»   1472»   1504»   1702»   1462»   1438»   1474»   1679»   1434»   1451»   1572»   1466»   1682»   1475» |  | 53 |  | 4717-04\_lib898» 1574»   1508»   1473»   1446»   1501»   1485»   1727»   1476»   1454»   1456»   1470»   1466»   1492»   1439»   1622»   1437»   1485»   1482»   1503»   1496»   1448»   1690»   1465»   1438»   1467»   1467»   1483»   1503»   1442»   1461»   1470»   1681»   195»1618»   1438»   1458»   1484»   1481»   1495»   1472»   1504»   1702»   1462»   1438»   1474»   1679»   1434»   1451»   1572»   1466»   1682»   1475» |
| 54 |  | 4724-03\_lib1517»1472»   938»903»876»931»915»1623»   904»884»888»901»899»924»871»1520»   869»915»914»37» 929»880»1587»   893»870»898»898»913»933»874»893»901»1578»   1481»   1516»   870»891»914»911»927»904»934»1598»   892»870»904»1576»   866»883»1470»   894»1578»   907»1500» |  | 54 |  | 4724-03\_lib1517»1472»   938»903»876»931»915»1623»   904»884»888»901»899»924»871»1520»   869»915»914»37» 929»880»1587»   893»870»898»898»913»933»874»893»901»1578»   1481»   1516»   870»891»914»911»927»904»934»1598»   892»870»904»1576»   866»883»1470»   894»1578»   907»1500» |  | 54 |  | 4724-03\_lib1517»1472»   938»903»876»931»915»1623»   904»884»888»901»899»924»871»1520»   869»915»914»37» 929»880»1587»   893»870»898»898»913»933»874»893»901»1578»   1481»   1516»   870»891»914»911»927»904»934»1598»   892»870»904»1576»   866»883»1470»   894»1578»   907»1500» |
| 55 |  | 4779-04\_lib944» 1448»   580»631»516»573»517»1601»   546»612»612»111»107»650»595»1496»   593»517»640»909»313»518»1566»   535»594»108»108»641»575»598»619»111»1557»   1454»   1492»   594»275»516»639»651»504»576»1576»   532»594»634»1555»   590»607»1446»   536»1556»   545»1473»   906» |  | 55 |  | 4779-04\_lib944» 1448»   580»631»516»573»517»1601»   546»612»612»111»107»650»595»1496»   593»517»640»909»313»518»1566»   535»594»108»108»641»575»598»619»111»1557»   1454»   1492»   594»275»516»639»651»504»576»1576»   532»594»634»1555»   590»607»1446»   536»1556»   545»1473»   906» |  | 55 |  | 4779-04\_lib944» 1448»   580»631»516»573»517»1601»   546»612»612»111»107»650»595»1496»   593»517»640»909»313»518»1566»   535»594»108»108»641»575»598»619»111»1557»   1454»   1492»   594»275»516»639»651»504»576»1576»   532»594»634»1555»   590»607»1446»   536»1556»   545»1473»   906» |
| 56 |  | 4781-04\_lib908» 1417»   633»462»569»626»606»1566»   599»441»195»594»590»481»2»  1465»   176»606»471»876»618»567»1531»   588»177»591»591»472»628»181»448»594»1522»   1422»   1461»   177»580»605»470»482»591»629»1541»   585»177»465»1520»   173»436»1415»   589»1521»   596»1441»   873»597» |  | 56 |  | 4781-04\_lib908» 1417»   633»462»569»626»606»1566»   599»441»195»594»590»481»2»  1465»   176»606»471»876»618»567»1531»   588»177»591»591»472»628»181»448»594»1522»   1422»   1461»   177»580»605»470»482»591»629»1541»   585»177»465»1520»   173»436»1415»   589»1521»   596»1441»   873»597» |  | 56 |  | 4781-04\_lib908» 1417»   633»462»569»626»606»1566»   599»441»195»594»590»481»2»  1465»   176»606»471»876»618»567»1531»   588»177»591»591»472»628»181»448»594»1522»   1422»   1461»   177»580»605»470»482»591»629»1541»   585»177»465»1520»   173»436»1415»   589»1521»   596»1441»   873»597» |
| 57 |  | 4783-04\_lib896» 1461»   591»642»529»584»154»1612»   557»620»623»514»512»661»606»1507»   604»154»651»920»540»529»1577»   548»605»511»511»652»586»609»630»514»1568»   1467»   1503»   605»502»153»650»662»499»587»1587»   545»605»645»1566»   601»618»1459»   549»1567»   556»1486»   917»519»608» |  | 57 |  | 4783-04\_lib896» 1461»   591»642»529»584»154»1612»   557»620»623»514»512»661»606»1507»   604»154»651»920»540»529»1577»   548»605»511»511»652»586»609»630»514»1568»   1467»   1503»   605»502»153»650»662»499»587»1587»   545»605»645»1566»   601»618»1459»   549»1567»   556»1486»   917»519»608» |  | 57 |  | 4783-04\_lib896» 1461»   591»642»529»584»154»1612»   557»620»623»514»512»661»606»1507»   604»154»651»920»540»529»1577»   548»605»511»511»652»586»609»630»514»1568»   1467»   1503»   605»502»153»650»662»499»587»1587»   545»605»645»1566»   601»618»1459»   549»1567»   556»1486»   917»519»608» |
| 58 |  | 4785-04\_lib909» 1418»   634»459»570»627»607»1567»   600»438»186»595»591»478»179»1466»   145»607»468»877»619»568»1532»   587»146»592»592»469»629»150»449»595»1523»   1423»   1462»   146»581»606»467»479»590»630»1542»   586»146»462»1521»   142»437»1416»   588»1522»   597»1442»   874»598»181»609» |  | 58 |  | 4785-04\_lib909» 1418»   634»459»570»627»607»1567»   600»438»186»595»591»478»179»1466»   145»607»468»877»619»568»1532»   587»146»592»592»469»629»150»449»595»1523»   1423»   1462»   146»581»606»467»479»590»630»1542»   586»146»462»1521»   142»437»1416»   588»1522»   597»1442»   874»598»181»609» |  | 58 |  | 4785-04\_lib909» 1418»   634»459»570»627»607»1567»   600»438»186»595»591»478»179»1466»   145»607»468»877»619»568»1532»   587»146»592»592»469»629»150»449»595»1523»   1423»   1462»   146»581»606»467»479»590»630»1542»   586»146»462»1521»   142»437»1416»   588»1522»   597»1442»   874»598»181»609» |
| 59 |  | 5248-04\_lib928» 1443»   661»494»597»654»636»1594»   627»475»473»624»620»513»456»1491»   454»638»503»902»650»600»1561»   616»455»621»621»504»656»459»174»624»1554»   1452»   1487»   455»612»637»500»514»625»657»1573»   613»455»493»1552»   451»350»1439»   617»1553»   628»1471»   899»627»458»640»459» |  | 59 |  | 5248-04\_lib928» 1443»   661»494»597»654»636»1594»   627»475»473»624»620»513»456»1491»   454»638»503»902»650»600»1561»   616»455»621»621»504»656»459»174»624»1554»   1452»   1487»   455»612»637»500»514»625»657»1573»   613»455»493»1552»   451»350»1439»   617»1553»   628»1471»   899»627»458»640»459» |  | 59 |  | 5248-04\_lib928» 1443»   661»494»597»654»636»1594»   627»475»473»624»620»513»456»1491»   454»638»503»902»650»600»1561»   616»455»621»621»504»656»459»174»624»1554»   1452»   1487»   455»612»637»500»514»625»657»1573»   613»455»493»1552»   451»350»1439»   617»1553»   628»1471»   899»627»458»640»459» |
| 60 |  | 5253-04\_lib961» 1538»   1602»   1563»   1536»   1595»   1575»   391»1568»   1544»   1546»   1563»   1559»   1582»   1529»   1588»   1527»   1575»   1572»   1593»   1587»   1538»   648»1557»   1528»   1560»   1560»   1573»   1597»   1532»   1553»   1563»   641»1673»   1584»   1528»   1549»   1574»   1573»   1585»   1560»   1598»   301»1552»   1528»   1566»   637»1524»   1541»   1538»   1558»   281»1565»   1692»   1590»   1566»   1531»   1577»   1532»   1563» |  | 60 |  | 5253-04\_lib961» 1538»   1602»   1563»   1536»   1595»   1575»   391»1568»   1544»   1546»   1563»   1559»   1582»   1529»   1588»   1527»   1575»   1572»   1593»   1587»   1538»   648»1557»   1528»   1560»   1560»   1573»   1597»   1532»   1553»   1563»   641»1673»   1584»   1528»   1549»   1574»   1573»   1585»   1560»   1598»   301»1552»   1528»   1566»   637»1524»   1541»   1538»   1558»   281»1565»   1692»   1590»   1566»   1531»   1577»   1532»   1563» |  | 60 |  | 5253-04\_lib961» 1538»   1602»   1563»   1536»   1595»   1575»   391»1568»   1544»   1546»   1563»   1559»   1582»   1529»   1588»   1527»   1575»   1572»   1593»   1587»   1538»   648»1557»   1528»   1560»   1560»   1573»   1597»   1532»   1553»   1563»   641»1673»   1584»   1528»   1549»   1574»   1573»   1585»   1560»   1598»   301»1552»   1528»   1566»   637»1524»   1541»   1538»   1558»   281»1565»   1692»   1590»   1566»   1531»   1577»   1532»   1563» |
| 61 |  | 5468-03\_lib912» 1479»   235»660»207»228»586»1632»   363»643»643»572»568»681»626»1527»   624»586»671»938»598»549»1597»   228»625»569»569»672»230»629»650»572»1588»   1486»   1523»   625»560»585»668»682»573»231»1607»   223»625»661»1586»   621»638»1475»   229»1587»   576»1505»   935»575»628»588»629»656»1597» |  | 61 |  | 5468-03\_lib912» 1479»   235»660»207»228»586»1632»   363»643»643»572»568»681»626»1527»   624»586»671»938»598»549»1597»   228»625»569»569»672»230»629»650»572»1588»   1486»   1523»   625»560»585»668»682»573»231»1607»   223»625»661»1586»   621»638»1475»   229»1587»   576»1505»   935»575»628»588»629»656»1597» |  | 61 |  | 5468-03\_lib912» 1479»   235»660»207»228»586»1632»   363»643»643»572»568»681»626»1527»   624»586»671»938»598»549»1597»   228»625»569»569»672»230»629»650»572»1588»   1486»   1523»   625»560»585»668»682»573»231»1607»   223»625»661»1586»   621»638»1475»   229»1587»   576»1505»   935»575»628»588»629»656»1597» |
| 62 |  | 5472-03\_lib938» 1441»   573»624»509»566»510»1594»   539»603»603»104»100»643»586»1489»   584»510»633»902»304»509»1559»   528»585»101»101»634»568»589»612»104»1550»   1447»   1485»   585»266»509»630»644»495»569»1569»   525»585»627»1548»   581»600»1439»   529»1549»   538»1466»   899»105»588»512»589»620»1559»   568» |  | 62 |  | 5472-03\_lib938» 1441»   573»624»509»566»510»1594»   539»603»603»104»100»643»586»1489»   584»510»633»902»304»509»1559»   528»585»101»101»634»568»589»612»104»1550»   1447»   1485»   585»266»509»630»644»495»569»1569»   525»585»627»1548»   581»600»1439»   529»1549»   538»1466»   899»105»588»512»589»620»1559»   568» |  | 62 |  | 5472-03\_lib938» 1441»   573»624»509»566»510»1594»   539»603»603»104»100»643»586»1489»   584»510»633»902»304»509»1559»   528»585»101»101»634»568»589»612»104»1550»   1447»   1485»   585»266»509»630»644»495»569»1569»   525»585»627»1548»   581»600»1439»   529»1549»   538»1466»   899»105»588»512»589»620»1559»   568» |
| 63 |  | 5685-04\_lib945» 1446»   576»629»514»569»513»1599»   542»608»610»1»  45» 648»593»1494»   591»513»638»905»311»516»1564»   533»592»8»  8»  639»571»596»617»17» 1555»   1452»   1490»   592»273»512»637»649»502»572»1574»   530»592»632»1553»   588»605»1444»   534»1554»   543»1471»   902»112»595»515»596»625»1564»   573»105» |  | 63 |  | 5685-04\_lib945» 1446»   576»629»514»569»513»1599»   542»608»610»1»  45» 648»593»1494»   591»513»638»905»311»516»1564»   533»592»8»  8»  639»571»596»617»17» 1555»   1452»   1490»   592»273»512»637»649»502»572»1574»   530»592»632»1553»   588»605»1444»   534»1554»   543»1471»   902»112»595»515»596»625»1564»   573»105» |  | 63 |  | 5685-04\_lib945» 1446»   576»629»514»569»513»1599»   542»608»610»1»  45» 648»593»1494»   591»513»638»905»311»516»1564»   533»592»8»  8»  639»571»596»617»17» 1555»   1452»   1490»   592»273»512»637»649»502»572»1574»   530»592»632»1553»   588»605»1444»   534»1554»   543»1471»   902»112»595»515»596»625»1564»   573»105» |
| 64 |  | 5687-04\_lib946» 1449»   581»632»517»574»518»1602»   547»613»613»112»108»651»596»1497»   594»518»641»910»312»519»1567»   536»595»109»109»642»576»599»620»112»1558»   1455»   1493»   595»274»517»640»652»505»577»1577»   533»595»635»1556»   591»608»1447»   537»1557»   546»1474»   907»113»598»520»599»628»1567»   576»56» 113» |  | 64 |  | 5687-04\_lib946» 1449»   581»632»517»574»518»1602»   547»613»613»112»108»651»596»1497»   594»518»641»910»312»519»1567»   536»595»109»109»642»576»599»620»112»1558»   1455»   1493»   595»274»517»640»652»505»577»1577»   533»595»635»1556»   591»608»1447»   537»1557»   546»1474»   907»113»598»520»599»628»1567»   576»56» 113» |  | 64 |  | 5687-04\_lib946» 1449»   581»632»517»574»518»1602»   547»613»613»112»108»651»596»1497»   594»518»641»910»312»519»1567»   536»595»109»109»642»576»599»620»112»1558»   1455»   1493»   595»274»517»640»652»505»577»1577»   533»595»635»1556»   591»608»1447»   537»1557»   546»1474»   907»113»598»520»599»628»1567»   576»56» 113» |
| 65 |  | 5870-03\_lib966» 1438»   1498»   1463»   1432»   1491»   1475»   583»1464»   1444»   1446»   1461»   1457»   1482»   1429»   1486»   1427»   1475»   1472»   1483»   1487»   1438»   230»1451»   1428»   1458»   1458»   1473»   1493»   1432»   1451»   1461»   223»1570»   1482»   1428»   1449»   1474»   1469»   1485»   1458»   1494»   558»1447»   1428»   1462»   219»1424»   1441»   1436»   1452»   538»1465»   1589»   1480»   1464»   1431»   1477»   1432»   1454»   550»1493»   1457»   1462»   1465» |  | 65 |  | 5870-03\_lib966» 1438»   1498»   1463»   1432»   1491»   1475»   583»1464»   1444»   1446»   1461»   1457»   1482»   1429»   1486»   1427»   1475»   1472»   1483»   1487»   1438»   230»1451»   1428»   1458»   1458»   1473»   1493»   1432»   1451»   1461»   223»1570»   1482»   1428»   1449»   1474»   1469»   1485»   1458»   1494»   558»1447»   1428»   1462»   219»1424»   1441»   1436»   1452»   538»1465»   1589»   1480»   1464»   1431»   1477»   1432»   1454»   550»1493»   1457»   1462»   1465» |  | 65 |  | 5870-03\_lib966» 1438»   1498»   1463»   1432»   1491»   1475»   583»1464»   1444»   1446»   1461»   1457»   1482»   1429»   1486»   1427»   1475»   1472»   1483»   1487»   1438»   230»1451»   1428»   1458»   1458»   1473»   1493»   1432»   1451»   1461»   223»1570»   1482»   1428»   1449»   1474»   1469»   1485»   1458»   1494»   558»1447»   1428»   1462»   219»1424»   1441»   1436»   1452»   538»1465»   1589»   1480»   1464»   1431»   1477»   1432»   1454»   550»1493»   1457»   1462»   1465» |
| 66 |  | 5872-03\_lib900» 1441»   655»434»593»648»630»1592»   621»415»469»618»616»455»452»1489»   448»630»445»898»642»595»1557»   610»449»615»615»446»648»453»474»618»1548»   1446»   1485»   449»606»629»444»456»617»651»1567»   609»449»437»1546»   445»464»1439»   611»1547»   622»1465»   895»623»454»632»451»482»1557»   652»616»619»624»1455» |  | 66 |  | 5872-03\_lib900» 1441»   655»434»593»648»630»1592»   621»415»469»618»616»455»452»1489»   448»630»445»898»642»595»1557»   610»449»615»615»446»648»453»474»618»1548»   1446»   1485»   449»606»629»444»456»617»651»1567»   609»449»437»1546»   445»464»1439»   611»1547»   622»1465»   895»623»454»632»451»482»1557»   652»616»619»624»1455» |  | 66 |  | 5872-03\_lib900» 1441»   655»434»593»648»630»1592»   621»415»469»618»616»455»452»1489»   448»630»445»898»642»595»1557»   610»449»615»615»446»648»453»474»618»1548»   1446»   1485»   449»606»629»444»456»617»651»1567»   609»449»437»1546»   445»464»1439»   611»1547»   622»1465»   895»623»454»632»451»482»1557»   652»616»619»624»1455» |
| 67 |  | 6429-03\_lib923» 1446»   662»282»598»655»635»1594»   628»420»472»623»619»301»455»1494»   451»635»291»905»647»598»1560»   615»452»620»620»292»657»456»479»623»1551»   1451»   1490»   451»609»634»292»303»620»658»1570»   614»452»285»1549»   448»467»1444»   616»1550»   624»1470»   902»626»457»637»452»489»1560»   657»619»624»627»1460»   431» |  | 67 |  | 6429-03\_lib923» 1446»   662»282»598»655»635»1594»   628»420»472»623»619»301»455»1494»   451»635»291»905»647»598»1560»   615»452»620»620»292»657»456»479»623»1551»   1451»   1490»   451»609»634»292»303»620»658»1570»   614»452»285»1549»   448»467»1444»   616»1550»   624»1470»   902»626»457»637»452»489»1560»   657»619»624»627»1460»   431» |  | 67 |  | 6429-03\_lib923» 1446»   662»282»598»655»635»1594»   628»420»472»623»619»301»455»1494»   451»635»291»905»647»598»1560»   615»452»620»620»292»657»456»479»623»1551»   1451»   1490»   451»609»634»292»303»620»658»1570»   614»452»285»1549»   448»467»1444»   616»1550»   624»1470»   902»626»457»637»452»489»1560»   657»619»624»627»1460»   431» |
| 68 |  | 6435-03\_lib955» 1522»   1580»   1539»   1514»   1573»   1553»   659»1546»   1520»   1524»   1541»   1537»   1558»   1507»   1570»   1503»   1553»   1548»   1568»   1565»   1516»   306»1531»   1504»   1538»   1538»   1549»   1575»   1508»   1529»   1541»   299»1651»   1566»   1504»   1527»   1552»   1549»   1561»   1536»   1576»   634»1529»   1504»   1542»   295»1500»   1519»   1520»   1532»   614»1543»   1670»   1565»   1544»   1509»   1555»   1508»   1541»   626»1575»   1537»   1542»   1545»   139»1533»   1536» |  | 68 |  | 6435-03\_lib955» 1522»   1580»   1539»   1514»   1573»   1553»   659»1546»   1520»   1524»   1541»   1537»   1558»   1507»   1570»   1503»   1553»   1548»   1568»   1565»   1516»   306»1531»   1504»   1538»   1538»   1549»   1575»   1508»   1529»   1541»   299»1651»   1566»   1504»   1527»   1552»   1549»   1561»   1536»   1576»   634»1529»   1504»   1542»   295»1500»   1519»   1520»   1532»   614»1543»   1670»   1565»   1544»   1509»   1555»   1508»   1541»   626»1575»   1537»   1542»   1545»   139»1533»   1536» |  | 68 |  | 6435-03\_lib955» 1522»   1580»   1539»   1514»   1573»   1553»   659»1546»   1520»   1524»   1541»   1537»   1558»   1507»   1570»   1503»   1553»   1548»   1568»   1565»   1516»   306»1531»   1504»   1538»   1538»   1549»   1575»   1508»   1529»   1541»   299»1651»   1566»   1504»   1527»   1552»   1549»   1561»   1536»   1576»   634»1529»   1504»   1542»   295»1500»   1519»   1520»   1532»   614»1543»   1670»   1565»   1544»   1509»   1555»   1508»   1541»   626»1575»   1537»   1542»   1545»   139»1533»   1536» |
| 69 |  | 6463-04\_lib910» 1442»   572»625»510»565»509»1595»   538»604»606»9»  41» 644»589»1490»   587»509»634»901»307»512»1560»   529»588»6»  6»  635»567»592»613»13» 1551»   1448»   1486»   588»269»508»633»645»498»568»1570»   526»588»628»1549»   584»601»1440»   530»1550»   539»1467»   898»108»591»511»592»621»1560»   569»101»10» 109»1458»   615»620»1538» |  | 69 |  | 6463-04\_lib910» 1442»   572»625»510»565»509»1595»   538»604»606»9»  41» 644»589»1490»   587»509»634»901»307»512»1560»   529»588»6»  6»  635»567»592»613»13» 1551»   1448»   1486»   588»269»508»633»645»498»568»1570»   526»588»628»1549»   584»601»1440»   530»1550»   539»1467»   898»108»591»511»592»621»1560»   569»101»10» 109»1458»   615»620»1538» |  | 69 |  | 6463-04\_lib910» 1442»   572»625»510»565»509»1595»   538»604»606»9»  41» 644»589»1490»   587»509»634»901»307»512»1560»   529»588»6»  6»  635»567»592»613»13» 1551»   1448»   1486»   588»269»508»633»645»498»568»1570»   526»588»628»1549»   584»601»1440»   530»1550»   539»1467»   898»108»591»511»592»621»1560»   569»101»10» 109»1458»   615»620»1538» |
| 70 |  | 6467-04\_lib1450»1413»   629»456»565»622»602»1562»   595»435»181»590»586»475»174»1461»   10» 602»465»872»614»563»1527»   582»11» 587»587»466»624»45» 444»590»1518»   1418»   1457»   41» 576»601»464»475»585»625»1537»   581»11» 459»1516»   7»  432»1411»   583»1517»   592»1437»   869»593»176»604»145»454»1527»   624»584»591»594»1427»   448»451»1503»   587» |  | 70 |  | 6467-04\_lib1450»1413»   629»456»565»622»602»1562»   595»435»181»590»586»475»174»1461»   10» 602»465»872»614»563»1527»   582»11» 587»587»466»624»45» 444»590»1518»   1418»   1457»   41» 576»601»464»475»585»625»1537»   581»11» 459»1516»   7»  432»1411»   583»1517»   592»1437»   869»593»176»604»145»454»1527»   624»584»591»594»1427»   448»451»1503»   587» |  | 70 |  | 6467-04\_lib1450»1413»   629»456»565»622»602»1562»   595»435»181»590»586»475»174»1461»   10» 602»465»872»614»563»1527»   582»11» 587»587»466»624»45» 444»590»1518»   1418»   1457»   41» 576»601»464»475»585»625»1537»   581»11» 459»1516»   7»  432»1411»   583»1517»   592»1437»   869»593»176»604»145»454»1527»   624»584»591»594»1427»   448»451»1503»   587» |
| 71 |  | 6637-04\_lib899» 1576»   1510»   1475»   1448»   1503»   1487»   1729»   1478»   1456»   1458»   1470»   1466»   1494»   1441»   1624»   1439»   1487»   1484»   1503»   1496»   1450»   1692»   1467»   1440»   1467»   1467»   1485»   1505»   1444»   1465»   1470»   1683»   195»1620»   1440»   1458»   1486»   1483»   1497»   1474»   1506»   1704»   1464»   1440»   1476»   1681»   1436»   1453»   1574»   1468»   1684»   1477»   210»1500»   1473»   1443»   1488»   1444»   1473»   1694»   1507»   1466»   1471»   1474»   1591»   1467»   1472»   1672»   1467»   1439» |  | 71 |  | 6637-04\_lib899» 1576»   1510»   1475»   1448»   1503»   1487»   1729»   1478»   1456»   1458»   1470»   1466»   1494»   1441»   1624»   1439»   1487»   1484»   1503»   1496»   1450»   1692»   1467»   1440»   1467»   1467»   1485»   1505»   1444»   1465»   1470»   1683»   195»1620»   1440»   1458»   1486»   1483»   1497»   1474»   1506»   1704»   1464»   1440»   1476»   1681»   1436»   1453»   1574»   1468»   1684»   1477»   210»1500»   1473»   1443»   1488»   1444»   1473»   1694»   1507»   1466»   1471»   1474»   1591»   1467»   1472»   1672»   1467»   1439» |  | 71 |  | 6637-04\_lib899» 1576»   1510»   1475»   1448»   1503»   1487»   1729»   1478»   1456»   1458»   1470»   1466»   1494»   1441»   1624»   1439»   1487»   1484»   1503»   1496»   1450»   1692»   1467»   1440»   1467»   1467»   1485»   1505»   1444»   1465»   1470»   1683»   195»1620»   1440»   1458»   1486»   1483»   1497»   1474»   1506»   1704»   1464»   1440»   1476»   1681»   1436»   1453»   1574»   1468»   1684»   1477»   210»1500»   1473»   1443»   1488»   1444»   1473»   1694»   1507»   1466»   1471»   1474»   1591»   1467»   1472»   1672»   1467»   1439» |
| 72 |  | 6639-04\_lib965» 1453»   367»634»305»360»558»1604»   3»  615»617»544»542»655»600»1501»   598»558»643»910»570»521»1571»   324»599»541»541»646»362»603»624»544»1562»   1460»   1497»   599»534»557»642»656»545»363»1581»   321»599»637»1560»   595»610»1449»   325»1561»   546»1479»   907»549»602»560»603»630»1571»   364»542»545»550»1467»   624»631»1549»   541»598»1481» |  | 72 |  | 6639-04\_lib965» 1453»   367»634»305»360»558»1604»   3»  615»617»544»542»655»600»1501»   598»558»643»910»570»521»1571»   324»599»541»541»646»362»603»624»544»1562»   1460»   1497»   599»534»557»642»656»545»363»1581»   321»599»637»1560»   595»610»1449»   325»1561»   546»1479»   907»549»602»560»603»630»1571»   364»542»545»550»1467»   624»631»1549»   541»598»1481» |  | 72 |  | 6639-04\_lib965» 1453»   367»634»305»360»558»1604»   3»  615»617»544»542»655»600»1501»   598»558»643»910»570»521»1571»   324»599»541»541»646»362»603»624»544»1562»   1460»   1497»   599»534»557»642»656»545»363»1581»   321»599»637»1560»   595»610»1449»   325»1561»   546»1479»   907»549»602»560»603»630»1571»   364»542»545»550»1467»   624»631»1549»   541»598»1481» |
| 73 |  | 6640-04\_lib929» 1424»   642»469»576»635»615»1573»   608»448»194»603»599»488»187»1472»   169»615»478»885»627»576»1538»   595»170»600»600»479»637»174»457»603»1529»   1429»   1468»   170»589»614»477»489»596»638»1548»   592»170»472»1527»   166»443»1422»   596»1528»   605»1448»   882»606»189»617»172»467»1538»   637»597»604»607»1438»   461»464»1514»   600»169»1450»   611» |  | 73 |  | 6640-04\_lib929» 1424»   642»469»576»635»615»1573»   608»448»194»603»599»488»187»1472»   169»615»478»885»627»576»1538»   595»170»600»600»479»637»174»457»603»1529»   1429»   1468»   170»589»614»477»489»596»638»1548»   592»170»472»1527»   166»443»1422»   596»1528»   605»1448»   882»606»189»617»172»467»1538»   637»597»604»607»1438»   461»464»1514»   600»169»1450»   611» |  | 73 |  | 6640-04\_lib929» 1424»   642»469»576»635»615»1573»   608»448»194»603»599»488»187»1472»   169»615»478»885»627»576»1538»   595»170»600»600»479»637»174»457»603»1529»   1429»   1468»   170»589»614»477»489»596»638»1548»   592»170»472»1527»   166»443»1422»   596»1528»   605»1448»   882»606»189»617»172»467»1538»   637»597»604»607»1438»   461»464»1514»   600»169»1450»   611» |
| 74 |  | 6769-04\_lib962» 1518»   1576»   1535»   1510»   1569»   1549»   654»1542»   1518»   1520»   1537»   1533»   1556»   1503»   1566»   1501»   1549»   1546»   1562»   1561»   1512»   302»1529»   1502»   1534»   1534»   1545»   1571»   1506»   1525»   1537»   295»1647»   1562»   1502»   1523»   1548»   1547»   1559»   1534»   1572»   629»1525»   1502»   1540»   291»1498»   1515»   1514»   1530»   609»1539»   1666»   1559»   1540»   1505»   1551»   1506»   1537»   621»1571»   1533»   1538»   1541»   136»1531»   1534»   202»1534»   1501»   1668»   1545»   1512» |  | 74 |  | 6769-04\_lib962» 1518»   1576»   1535»   1510»   1569»   1549»   654»1542»   1518»   1520»   1537»   1533»   1556»   1503»   1566»   1501»   1549»   1546»   1562»   1561»   1512»   302»1529»   1502»   1534»   1534»   1545»   1571»   1506»   1525»   1537»   295»1647»   1562»   1502»   1523»   1548»   1547»   1559»   1534»   1572»   629»1525»   1502»   1540»   291»1498»   1515»   1514»   1530»   609»1539»   1666»   1559»   1540»   1505»   1551»   1506»   1537»   621»1571»   1533»   1538»   1541»   136»1531»   1534»   202»1534»   1501»   1668»   1545»   1512» |  | 74 |  | 6769-04\_lib962» 1518»   1576»   1535»   1510»   1569»   1549»   654»1542»   1518»   1520»   1537»   1533»   1556»   1503»   1566»   1501»   1549»   1546»   1562»   1561»   1512»   302»1529»   1502»   1534»   1534»   1545»   1571»   1506»   1525»   1537»   295»1647»   1562»   1502»   1523»   1548»   1547»   1559»   1534»   1572»   629»1525»   1502»   1540»   291»1498»   1515»   1514»   1530»   609»1539»   1666»   1559»   1540»   1505»   1551»   1506»   1537»   621»1571»   1533»   1538»   1541»   136»1531»   1534»   202»1534»   1501»   1668»   1545»   1512» |
| 75 |  | 6771-04\_lib1462»1431»   561»610»497»554»534»1577»   521»591»591»522»518»629»574»1479»   572»534»617»890»542»201»1544»   516»573»519»519»620»554»577»598»522»1535»   1436»   1475»   573»506»533»618»630»519»557»1554»   513»573»613»1533»   569»584»1429»   517»1534»   210»1455»   887»525»576»536»577»608»1544»   556»516»523»524»1444»   600»605»1522»   519»572»1457»   524»585»1518» |  | 75 |  | 6771-04\_lib1462»1431»   561»610»497»554»534»1577»   521»591»591»522»518»629»574»1479»   572»534»617»890»542»201»1544»   516»573»519»519»620»554»577»598»522»1535»   1436»   1475»   573»506»533»618»630»519»557»1554»   513»573»613»1533»   569»584»1429»   517»1534»   210»1455»   887»525»576»536»577»608»1544»   556»516»523»524»1444»   600»605»1522»   519»572»1457»   524»585»1518» |  | 75 |  | 6771-04\_lib1462»1431»   561»610»497»554»534»1577»   521»591»591»522»518»629»574»1479»   572»534»617»890»542»201»1544»   516»573»519»519»620»554»577»598»522»1535»   1436»   1475»   573»506»533»618»630»519»557»1554»   513»573»613»1533»   569»584»1429»   517»1534»   210»1455»   887»525»576»536»577»608»1544»   556»516»523»524»1444»   600»605»1522»   519»572»1457»   524»585»1518» |
| 76 |  | 6775-04\_lib963» 1542»   1598»   1561»   1534»   1591»   1573»   679»1566»   1542»   1544»   1561»   1557»   1580»   1527»   1590»   1525»   1573»   1570»   1588»   1585»   1536»   108»1553»   1526»   1558»   1558»   1571»   1593»   1530»   1549»   1561»   103»1669»   1586»   1526»   1547»   1572»   1571»   1583»   1558»   1594»   654»1549»   1526»   1564»   29» 1522»   1539»   1540»   1554»   634»1563»   1688»   1585»   1564»   1529»   1575»   1530»   1561»   646»1595»   1557»   1562»   1565»   228»1555»   1558»   304»1558»   1525»   1690»   1569»   1536»   300»1542» |  | 76 |  | 6775-04\_lib963» 1542»   1598»   1561»   1534»   1591»   1573»   679»1566»   1542»   1544»   1561»   1557»   1580»   1527»   1590»   1525»   1573»   1570»   1588»   1585»   1536»   108»1553»   1526»   1558»   1558»   1571»   1593»   1530»   1549»   1561»   103»1669»   1586»   1526»   1547»   1572»   1571»   1583»   1558»   1594»   654»1549»   1526»   1564»   29» 1522»   1539»   1540»   1554»   634»1563»   1688»   1585»   1564»   1529»   1575»   1530»   1561»   646»1595»   1557»   1562»   1565»   228»1555»   1558»   304»1558»   1525»   1690»   1569»   1536»   300»1542» |  | 76 |  | 6775-04\_lib963» 1542»   1598»   1561»   1534»   1591»   1573»   679»1566»   1542»   1544»   1561»   1557»   1580»   1527»   1590»   1525»   1573»   1570»   1588»   1585»   1536»   108»1553»   1526»   1558»   1558»   1571»   1593»   1530»   1549»   1561»   103»1669»   1586»   1526»   1547»   1572»   1571»   1583»   1558»   1594»   654»1549»   1526»   1564»   29» 1522»   1539»   1540»   1554»   634»1563»   1688»   1585»   1564»   1529»   1575»   1530»   1561»   646»1595»   1557»   1562»   1565»   228»1555»   1558»   304»1558»   1525»   1690»   1569»   1536»   300»1542» |
| 77 |  | 6892-04\_lib964» 1542»   1600»   1563»   1536»   1593»   1575»   683»1568»   1544»   1546»   1563»   1559»   1582»   1529»   1592»   1527»   1575»   1572»   1592»   1587»   1538»   64» 1557»   1528»   1560»   1560»   1573»   1595»   1532»   1551»   1563»   107»1671»   1588»   1528»   1549»   1574»   1573»   1585»   1560»   1596»   658»1551»   1528»   1566»   101»1524»   1539»   1542»   1558»   638»1565»   1690»   1589»   1566»   1531»   1577»   1532»   1561»   646»1597»   1559»   1564»   1567»   232»1557»   1560»   308»1560»   1527»   1692»   1571»   1538»   304»1544»   110» |  | 77 |  | 6892-04\_lib964» 1542»   1600»   1563»   1536»   1593»   1575»   683»1568»   1544»   1546»   1563»   1559»   1582»   1529»   1592»   1527»   1575»   1572»   1592»   1587»   1538»   64» 1557»   1528»   1560»   1560»   1573»   1595»   1532»   1551»   1563»   107»1671»   1588»   1528»   1549»   1574»   1573»   1585»   1560»   1596»   658»1551»   1528»   1566»   101»1524»   1539»   1542»   1558»   638»1565»   1690»   1589»   1566»   1531»   1577»   1532»   1561»   646»1597»   1559»   1564»   1567»   232»1557»   1560»   308»1560»   1527»   1692»   1571»   1538»   304»1544»   110» |  | 77 |  | 6892-04\_lib964» 1542»   1600»   1563»   1536»   1593»   1575»   683»1568»   1544»   1546»   1563»   1559»   1582»   1529»   1592»   1527»   1575»   1572»   1592»   1587»   1538»   64» 1557»   1528»   1560»   1560»   1573»   1595»   1532»   1551»   1563»   107»1671»   1588»   1528»   1549»   1574»   1573»   1585»   1560»   1596»   658»1551»   1528»   1566»   101»1524»   1539»   1542»   1558»   638»1565»   1690»   1589»   1566»   1531»   1577»   1532»   1561»   646»1597»   1559»   1564»   1567»   232»1557»   1560»   308»1560»   1527»   1692»   1571»   1538»   304»1544»   110» |
| 78 |  | 6895-04\_lib1459»1558»   1489»   1455»   1427»   1482»   1468»   1709»   1455»   1436»   1439»   1453»   1449»   1474»   1422»   1606»   1419»   1468»   1464»   1482»   1477»   1431»   1674»   1445»   1420»   1450»   1450»   1465»   1484»   1424»   1446»   1453»   1665»   453»1602»   1420»   1441»   1467»   1463»   1477»   1452»   1485»   1685»   1443»   1420»   1456»   1663»   1416»   1432»   1556»   1446»   1665»   1458»   472»1479»   1456»   1424»   1470»   1424»   1452»   1675»   1486»   1449»   1454»   1457»   1572»   1447»   1452»   1653»   1450»   1419»   474»1458»   1430»   1650»   1436»   1672»   1674» |  | 78 |  | 6895-04\_lib1459»1558»   1489»   1455»   1427»   1482»   1468»   1709»   1455»   1436»   1439»   1453»   1449»   1474»   1422»   1606»   1419»   1468»   1464»   1482»   1477»   1431»   1674»   1445»   1420»   1450»   1450»   1465»   1484»   1424»   1446»   1453»   1665»   453»1602»   1420»   1441»   1467»   1463»   1477»   1452»   1485»   1685»   1443»   1420»   1456»   1663»   1416»   1432»   1556»   1446»   1665»   1458»   472»1479»   1456»   1424»   1470»   1424»   1452»   1675»   1486»   1449»   1454»   1457»   1572»   1447»   1452»   1653»   1450»   1419»   474»1458»   1430»   1650»   1436»   1672»   1674» |  | 78 |  | 6895-04\_lib1459»1558»   1489»   1455»   1427»   1482»   1468»   1709»   1455»   1436»   1439»   1453»   1449»   1474»   1422»   1606»   1419»   1468»   1464»   1482»   1477»   1431»   1674»   1445»   1420»   1450»   1450»   1465»   1484»   1424»   1446»   1453»   1665»   453»1602»   1420»   1441»   1467»   1463»   1477»   1452»   1485»   1685»   1443»   1420»   1456»   1663»   1416»   1432»   1556»   1446»   1665»   1458»   472»1479»   1456»   1424»   1470»   1424»   1452»   1675»   1486»   1449»   1454»   1457»   1572»   1447»   1452»   1653»   1450»   1419»   474»1458»   1430»   1650»   1436»   1672»   1674» |
| 79 |  | 6897-04\_lib954» 346»1542»   1507»   1476»   1535»   1515»   1635»   1508»   1488»   1490»   1503»   1499»   1526»   1473»   92» 1471»   1515»   1516»   1533»   1529»   1482»   1602»   1497»   1472»   1500»   1500»   1517»   1537»   1474»   1497»   1503»   1595»   1613»   72» 1472»   1491»   1514»   1513»   1529»   1499»   1538»   1611»   1492»   1472»   1506»   1591»   1468»   1485»   342»1498»   1591»   1509»   1632»   1530»   1506»   1475»   1517»   1476»   1501»   1598»   1537»   1499»   1504»   1507»   1496»   1499»   1502»   1580»   1500»   1471»   1634»   1511»   1482»   1576»   1489»   1600»   1602»   1616» |  | 79 |  | 6897-04\_lib954» 346»1542»   1507»   1476»   1535»   1515»   1635»   1508»   1488»   1490»   1503»   1499»   1526»   1473»   92» 1471»   1515»   1516»   1533»   1529»   1482»   1602»   1497»   1472»   1500»   1500»   1517»   1537»   1474»   1497»   1503»   1595»   1613»   72» 1472»   1491»   1514»   1513»   1529»   1499»   1538»   1611»   1492»   1472»   1506»   1591»   1468»   1485»   342»1498»   1591»   1509»   1632»   1530»   1506»   1475»   1517»   1476»   1501»   1598»   1537»   1499»   1504»   1507»   1496»   1499»   1502»   1580»   1500»   1471»   1634»   1511»   1482»   1576»   1489»   1600»   1602»   1616» |  | 79 |  | 6897-04\_lib954» 346»1542»   1507»   1476»   1535»   1515»   1635»   1508»   1488»   1490»   1503»   1499»   1526»   1473»   92» 1471»   1515»   1516»   1533»   1529»   1482»   1602»   1497»   1472»   1500»   1500»   1517»   1537»   1474»   1497»   1503»   1595»   1613»   72» 1472»   1491»   1514»   1513»   1529»   1499»   1538»   1611»   1492»   1472»   1506»   1591»   1468»   1485»   342»1498»   1591»   1509»   1632»   1530»   1506»   1475»   1517»   1476»   1501»   1598»   1537»   1499»   1504»   1507»   1496»   1499»   1502»   1580»   1500»   1471»   1634»   1511»   1482»   1576»   1489»   1600»   1602»   1616» |
| 80 |  | 7000-03\_lib913» 1475»   393»658»327»386»584»1626»   197»639»641»570»566»679»624»1523»   622»584»669»934»594»547»1593»   348»623»567»567»670»388»627»648»570»1584»   1484»   1519»   623»558»583»666»680»567»389»1603»   343»623»661»1582»   617»634»1471»   349»1583»   574»1503»   931»573»626»586»627»654»1593»   388»566»571»574»1489»   648»655»1571»   567»622»1505»   198»633»1567»   552»1591»   1593»   1482»   1533» |  | 80 |  | 7000-03\_lib913» 1475»   393»658»327»386»584»1626»   197»639»641»570»566»679»624»1523»   622»584»669»934»594»547»1593»   348»623»567»567»670»388»627»648»570»1584»   1484»   1519»   623»558»583»666»680»567»389»1603»   343»623»661»1582»   617»634»1471»   349»1583»   574»1503»   931»573»626»586»627»654»1593»   388»566»571»574»1489»   648»655»1571»   567»622»1505»   198»633»1567»   552»1591»   1593»   1482»   1533» |  | 80 |  | 7000-03\_lib913» 1475»   393»658»327»386»584»1626»   197»639»641»570»566»679»624»1523»   622»584»669»934»594»547»1593»   348»623»567»567»670»388»627»648»570»1584»   1484»   1519»   623»558»583»666»680»567»389»1603»   343»623»661»1582»   617»634»1471»   349»1583»   574»1503»   931»573»626»586»627»654»1593»   388»566»571»574»1489»   648»655»1571»   567»622»1505»   198»633»1567»   552»1591»   1593»   1482»   1533» |
| 81 |  | 7135-04\_lib987» 1474»   940»905»878»933»917»1625»   906»886»890»903»901»926»873»1522»   871»917»916»5»  931»882»1589»   895»872»900»900»915»935»876»895»903»1580»   1483»   1518»   872»893»916»913»929»906»936»1600»   894»872»906»1578»   868»885»1472»   896»1580»   909»1502»   38» 908»875»919»876»901»1592»   937»901»904»909»1482»   897»904»1567»   900»871»1502»   909»884»1561»   889»1587»   1591»   1481»   1532»   933» |  | 81 |  | 7135-04\_lib987» 1474»   940»905»878»933»917»1625»   906»886»890»903»901»926»873»1522»   871»917»916»5»  931»882»1589»   895»872»900»900»915»935»876»895»903»1580»   1483»   1518»   872»893»916»913»929»906»936»1600»   894»872»906»1578»   868»885»1472»   896»1580»   909»1502»   38» 908»875»919»876»901»1592»   937»901»904»909»1482»   897»904»1567»   900»871»1502»   909»884»1561»   889»1587»   1591»   1481»   1532»   933» |  | 81 |  | 7135-04\_lib987» 1474»   940»905»878»933»917»1625»   906»886»890»903»901»926»873»1522»   871»917»916»5»  931»882»1589»   895»872»900»900»915»935»876»895»903»1580»   1483»   1518»   872»893»916»913»929»906»936»1600»   894»872»906»1578»   868»885»1472»   896»1580»   909»1502»   38» 908»875»919»876»901»1592»   937»901»904»909»1482»   897»904»1567»   900»871»1502»   909»884»1561»   889»1587»   1591»   1481»   1532»   933» |
| 82 |  | 7514-04\_lib1460»1469»   385»650»321»378»576»1620»   247»633»633»562»558»671»616»1517»   614»576»659»928»586»539»1587»   340»615»559»559»662»380»619»638»562»1578»   1476»   1513»   615»550»575»658»672»561»381»1597»   337»615»653»1576»   611»624»1463»   341»1577»   564»1495»   925»565»618»578»619»644»1587»   380»558»563»566»1483»   642»647»1565»   559»614»1497»   248»627»1561»   542»1585»   1587»   1474»   1527»   274»927» |  | 82 |  | 7514-04\_lib1460»1469»   385»650»321»378»576»1620»   247»633»633»562»558»671»616»1517»   614»576»659»928»586»539»1587»   340»615»559»559»662»380»619»638»562»1578»   1476»   1513»   615»550»575»658»672»561»381»1597»   337»615»653»1576»   611»624»1463»   341»1577»   564»1495»   925»565»618»578»619»644»1587»   380»558»563»566»1483»   642»647»1565»   559»614»1497»   248»627»1561»   542»1585»   1587»   1474»   1527»   274»927» |  | 82 |  | 7514-04\_lib1460»1469»   385»650»321»378»576»1620»   247»633»633»562»558»671»616»1517»   614»576»659»928»586»539»1587»   340»615»559»559»662»380»619»638»562»1578»   1476»   1513»   615»550»575»658»672»561»381»1597»   337»615»653»1576»   611»624»1463»   341»1577»   564»1495»   925»565»618»578»619»644»1587»   380»558»563»566»1483»   642»647»1565»   559»614»1497»   248»627»1561»   542»1585»   1587»   1474»   1527»   274»927» |
| 83 |  | 7516-04\_lib920» 1475»   391»656»327»384»582»1626»   253»639»639»568»564»677»622»1523»   620»582»665»934»592»545»1593»   346»621»565»565»668»386»625»644»568»1584»   1482»   1519»   621»556»581»664»678»567»387»1603»   343»621»659»1582»   617»630»1469»   347»1583»   570»1501»   931»571»624»584»625»650»1593»   386»564»569»572»1489»   648»653»1571»   565»620»1503»   254»633»1567»   548»1591»   1593»   1480»   1533»   280»933»46» |  | 83 |  | 7516-04\_lib920» 1475»   391»656»327»384»582»1626»   253»639»639»568»564»677»622»1523»   620»582»665»934»592»545»1593»   346»621»565»565»668»386»625»644»568»1584»   1482»   1519»   621»556»581»664»678»567»387»1603»   343»621»659»1582»   617»630»1469»   347»1583»   570»1501»   931»571»624»584»625»650»1593»   386»564»569»572»1489»   648»653»1571»   565»620»1503»   254»633»1567»   548»1591»   1593»   1480»   1533»   280»933»46» |  | 83 |  | 7516-04\_lib920» 1475»   391»656»327»384»582»1626»   253»639»639»568»564»677»622»1523»   620»582»665»934»592»545»1593»   346»621»565»565»668»386»625»644»568»1584»   1482»   1519»   621»556»581»664»678»567»387»1603»   343»621»659»1582»   617»630»1469»   347»1583»   570»1501»   931»571»624»584»625»650»1593»   386»564»569»572»1489»   648»653»1571»   565»620»1503»   254»633»1567»   548»1591»   1593»   1480»   1533»   280»933»46» |
| 84 |  | 7517-04\_lib930» 1442»   660»493»596»653»635»1593»   626»474»472»623»619»512»455»1490»   453»637»502»901»649»599»1560»   615»454»620»620»503»655»458»173»623»1553»   1451»   1486»   454»611»636»499»513»624»656»1572»   612»454»492»1551»   450»349»1438»   616»1552»   627»1470»   898»626»457»639»458»3»  1562»   655»619»624»627»1453»   481»488»1540»   620»453»1472»   629»466»1536»   607»1560»   1560»   1451»   1500»   653»900»643»649» |  | 84 |  | 7517-04\_lib930» 1442»   660»493»596»653»635»1593»   626»474»472»623»619»512»455»1490»   453»637»502»901»649»599»1560»   615»454»620»620»503»655»458»173»623»1553»   1451»   1486»   454»611»636»499»513»624»656»1572»   612»454»492»1551»   450»349»1438»   616»1552»   627»1470»   898»626»457»639»458»3»  1562»   655»619»624»627»1453»   481»488»1540»   620»453»1472»   629»466»1536»   607»1560»   1560»   1451»   1500»   653»900»643»649» |  | 84 |  | 7517-04\_lib930» 1442»   660»493»596»653»635»1593»   626»474»472»623»619»512»455»1490»   453»637»502»901»649»599»1560»   615»454»620»620»503»655»458»173»623»1553»   1451»   1486»   454»611»636»499»513»624»656»1572»   612»454»492»1551»   450»349»1438»   616»1552»   627»1470»   898»626»457»639»458»3»  1562»   655»619»624»627»1453»   481»488»1540»   620»453»1472»   629»466»1536»   607»1560»   1560»   1451»   1500»   653»900»643»649» |
| 85 |  | 7520-04\_lib1461»1482»   38» 663»212»31» 587»1635»   364»644»646»573»571»684»629»1530»   627»587»674»939»601»552»1598»   231»628»570»570»675»31» 632»653»573»1589»   1487»   1526»   628»563»586»671»685»576»34» 1610»   228»628»664»1587»   624»641»1478»   232»1590»   579»1506»   936»578»631»589»632»659»1600»   233»571»574»579»1496»   653»660»1578»   570»627»1508»   365»640»1574»   559»1596»   1598»   1487»   1540»   391»938»383»389»658» |  | 85 |  | 7520-04\_lib1461»1482»   38» 663»212»31» 587»1635»   364»644»646»573»571»684»629»1530»   627»587»674»939»601»552»1598»   231»628»570»570»675»31» 632»653»573»1589»   1487»   1526»   628»563»586»671»685»576»34» 1610»   228»628»664»1587»   624»641»1478»   232»1590»   579»1506»   936»578»631»589»632»659»1600»   233»571»574»579»1496»   653»660»1578»   570»627»1508»   365»640»1574»   559»1596»   1598»   1487»   1540»   391»938»383»389»658» |  | 85 |  | 7520-04\_lib1461»1482»   38» 663»212»31» 587»1635»   364»644»646»573»571»684»629»1530»   627»587»674»939»601»552»1598»   231»628»570»570»675»31» 632»653»573»1589»   1487»   1526»   628»563»586»671»685»576»34» 1610»   228»628»664»1587»   624»641»1478»   232»1590»   579»1506»   936»578»631»589»632»659»1600»   233»571»574»579»1496»   653»660»1578»   570»627»1508»   365»640»1574»   559»1596»   1598»   1487»   1540»   391»938»383»389»658» |
| 86 |  | 7538-03\_lib948» 1442»   660»493»596»653»635»1593»   626»474»472»623»619»512»455»1490»   453»637»502»901»649»599»1560»   615»454»620»620»503»655»458»173»623»1553»   1451»   1486»   454»611»636»499»513»624»656»1572»   612»454»492»1551»   450»349»1438»   616»1552»   627»1470»   898»626»457»639»458»3»  1562»   655»619»624»627»1453»   481»488»1540»   620»453»1472»   629»466»1536»   607»1560»   1560»   1451»   1500»   653»900»643»649»2»  658» |  | 86 |  | 7538-03\_lib948» 1442»   660»493»596»653»635»1593»   626»474»472»623»619»512»455»1490»   453»637»502»901»649»599»1560»   615»454»620»620»503»655»458»173»623»1553»   1451»   1486»   454»611»636»499»513»624»656»1572»   612»454»492»1551»   450»349»1438»   616»1552»   627»1470»   898»626»457»639»458»3»  1562»   655»619»624»627»1453»   481»488»1540»   620»453»1472»   629»466»1536»   607»1560»   1560»   1451»   1500»   653»900»643»649»2»  658» |  | 86 |  | 7538-03\_lib948» 1442»   660»493»596»653»635»1593»   626»474»472»623»619»512»455»1490»   453»637»502»901»649»599»1560»   615»454»620»620»503»655»458»173»623»1553»   1451»   1486»   454»611»636»499»513»624»656»1572»   612»454»492»1551»   450»349»1438»   616»1552»   627»1470»   898»626»457»639»458»3»  1562»   655»619»624»627»1453»   481»488»1540»   620»453»1472»   629»466»1536»   607»1560»   1560»   1451»   1500»   653»900»643»649»2»  658» |
| 87 |  | 8082-03\_lib932» 1459»   677»120»613»670»652»1610»   643»437»489»640»636»151»472»1507»   468»652»117»916»664»615»1575»   630»469»637»637»130»672»473»496»640»1566»   1464»   1503»   467»626»651»116»214»637»673»1585»   629»469»107»1564»   465»484»1457»   631»1565»   640»1483»   913»643»474»654»471»504»1575»   672»636»641»644»1473»   446»294»1551»   637»468»1485»   646»481»1549»   622»1573»   1575»   1465»   1517»   670»915»662»668»503»675»503» |  | 87 |  | 8082-03\_lib932» 1459»   677»120»613»670»652»1610»   643»437»489»640»636»151»472»1507»   468»652»117»916»664»615»1575»   630»469»637»637»130»672»473»496»640»1566»   1464»   1503»   467»626»651»116»214»637»673»1585»   629»469»107»1564»   465»484»1457»   631»1565»   640»1483»   913»643»474»654»471»504»1575»   672»636»641»644»1473»   446»294»1551»   637»468»1485»   646»481»1549»   622»1573»   1575»   1465»   1517»   670»915»662»668»503»675»503» |  | 87 |  | 8082-03\_lib932» 1459»   677»120»613»670»652»1610»   643»437»489»640»636»151»472»1507»   468»652»117»916»664»615»1575»   630»469»637»637»130»672»473»496»640»1566»   1464»   1503»   467»626»651»116»214»637»673»1585»   629»469»107»1564»   465»484»1457»   631»1565»   640»1483»   913»643»474»654»471»504»1575»   672»636»641»644»1473»   446»294»1551»   637»468»1485»   646»481»1549»   622»1573»   1575»   1465»   1517»   670»915»662»668»503»675»503» |
| 88 |  | 8864-03\_lib967» 1556»   1614»   1577»   1550»   1607»   1589»   697»1582»   1558»   1560»   1577»   1573»   1596»   1543»   1604»   1541»   1589»   1586»   1606»   1601»   1552»   64» 1571»   1542»   1574»   1574»   1587»   1609»   1546»   1565»   1577»   121»1685»   1600»   1542»   1563»   1588»   1587»   1599»   1574»   1610»   672»1565»   1542»   1580»   115»1538»   1553»   1554»   1572»   652»1579»   1704»   1603»   1580»   1545»   1591»   1546»   1575»   662»1611»   1573»   1578»   1581»   246»1571»   1574»   322»1574»   1541»   1706»   1585»   1552»   318»1558»   124»78» 1688»   1614»   1607»   1605»   1601»   1607»   1574»   1612»   1574»   1589» |  | 88 |  | 8864-03\_lib967» 1556»   1614»   1577»   1550»   1607»   1589»   697»1582»   1558»   1560»   1577»   1573»   1596»   1543»   1604»   1541»   1589»   1586»   1606»   1601»   1552»   64» 1571»   1542»   1574»   1574»   1587»   1609»   1546»   1565»   1577»   121»1685»   1600»   1542»   1563»   1588»   1587»   1599»   1574»   1610»   672»1565»   1542»   1580»   115»1538»   1553»   1554»   1572»   652»1579»   1704»   1603»   1580»   1545»   1591»   1546»   1575»   662»1611»   1573»   1578»   1581»   246»1571»   1574»   322»1574»   1541»   1706»   1585»   1552»   318»1558»   124»78» 1688»   1614»   1607»   1605»   1601»   1607»   1574»   1612»   1574»   1589» |  | 88 |  | 8864-03\_lib967» 1556»   1614»   1577»   1550»   1607»   1589»   697»1582»   1558»   1560»   1577»   1573»   1596»   1543»   1604»   1541»   1589»   1586»   1606»   1601»   1552»   64» 1571»   1542»   1574»   1574»   1587»   1609»   1546»   1565»   1577»   121»1685»   1600»   1542»   1563»   1588»   1587»   1599»   1574»   1610»   672»1565»   1542»   1580»   115»1538»   1553»   1554»   1572»   652»1579»   1704»   1603»   1580»   1545»   1591»   1546»   1575»   662»1611»   1573»   1578»   1581»   246»1571»   1574»   322»1574»   1541»   1706»   1585»   1552»   318»1558»   124»78» 1688»   1614»   1607»   1605»   1601»   1607»   1574»   1612»   1574»   1589» |
| 89 |  | 8867-03\_lib968» 1537»   1593»   1556»   1529»   1586»   1568»   676»1561»   1537»   1539»   1556»   1552»   1575»   1522»   1585»   1520»   1568»   1565»   1585»   1580»   1531»   163»1550»   1521»   1553»   1553»   1566»   1588»   1525»   1546»   1556»   156»1664»   1581»   1521»   1542»   1567»   1566»   1578»   1553»   1589»   651»1544»   1521»   1559»   152»1517»   1534»   1535»   1551»   631»1558»   1683»   1582»   1559»   1524»   1570»   1525»   1556»   641»1590»   1552»   1557»   1560»   225»1550»   1553»   301»1553»   1520»   1685»   1564»   1531»   297»1537»   161»163»1667»   1595»   1586»   1584»   1580»   1586»   1555»   1591»   1555»   1568»   177» |  | 89 |  | 8867-03\_lib968» 1537»   1593»   1556»   1529»   1586»   1568»   676»1561»   1537»   1539»   1556»   1552»   1575»   1522»   1585»   1520»   1568»   1565»   1585»   1580»   1531»   163»1550»   1521»   1553»   1553»   1566»   1588»   1525»   1546»   1556»   156»1664»   1581»   1521»   1542»   1567»   1566»   1578»   1553»   1589»   651»1544»   1521»   1559»   152»1517»   1534»   1535»   1551»   631»1558»   1683»   1582»   1559»   1524»   1570»   1525»   1556»   641»1590»   1552»   1557»   1560»   225»1550»   1553»   301»1553»   1520»   1685»   1564»   1531»   297»1537»   161»163»1667»   1595»   1586»   1584»   1580»   1586»   1555»   1591»   1555»   1568»   177» |  | 89 |  | 8867-03\_lib968» 1537»   1593»   1556»   1529»   1586»   1568»   676»1561»   1537»   1539»   1556»   1552»   1575»   1522»   1585»   1520»   1568»   1565»   1585»   1580»   1531»   163»1550»   1521»   1553»   1553»   1566»   1588»   1525»   1546»   1556»   156»1664»   1581»   1521»   1542»   1567»   1566»   1578»   1553»   1589»   651»1544»   1521»   1559»   152»1517»   1534»   1535»   1551»   631»1558»   1683»   1582»   1559»   1524»   1570»   1525»   1556»   641»1590»   1552»   1557»   1560»   225»1550»   1553»   301»1553»   1520»   1685»   1564»   1531»   297»1537»   161»163»1667»   1595»   1586»   1584»   1580»   1586»   1555»   1591»   1555»   1568»   177» |
| 90 |  | 8868-03\_lib950» 336»1524»   1489»   1458»   1517»   1499»   1617»   1490»   1470»   1472»   1485»   1481»   1508»   1455»   382»1453»   1499»   1498»   1515»   1511»   1463»   1584»   1479»   1454»   1482»   1482»   1499»   1519»   1456»   1479»   1485»   1577»   1597»   378»1454»   1473»   1498»   1495»   1511»   1482»   1520»   1593»   1474»   1454»   1488»   1573»   1450»   1467»   332»1480»   1573»   1490»   1616»   1512»   1488»   1457»   1501»   1458»   1483»   1582»   1519»   1481»   1486»   1489»   1478»   1481»   1486»   1562»   1482»   1453»   1618»   1493»   1464»   1558»   1470»   1582»   1584»   1598»   392»1515»   1514»   1509»   1515»   1482»   1522»   1482»   1499»   1596»   1577» |  | 90 |  | 8868-03\_lib950» 336»1524»   1489»   1458»   1517»   1499»   1617»   1490»   1470»   1472»   1485»   1481»   1508»   1455»   382»1453»   1499»   1498»   1515»   1511»   1463»   1584»   1479»   1454»   1482»   1482»   1499»   1519»   1456»   1479»   1485»   1577»   1597»   378»1454»   1473»   1498»   1495»   1511»   1482»   1520»   1593»   1474»   1454»   1488»   1573»   1450»   1467»   332»1480»   1573»   1490»   1616»   1512»   1488»   1457»   1501»   1458»   1483»   1582»   1519»   1481»   1486»   1489»   1478»   1481»   1486»   1562»   1482»   1453»   1618»   1493»   1464»   1558»   1470»   1582»   1584»   1598»   392»1515»   1514»   1509»   1515»   1482»   1522»   1482»   1499»   1596»   1577» |  | 90 |  | 8868-03\_lib950» 336»1524»   1489»   1458»   1517»   1499»   1617»   1490»   1470»   1472»   1485»   1481»   1508»   1455»   382»1453»   1499»   1498»   1515»   1511»   1463»   1584»   1479»   1454»   1482»   1482»   1499»   1519»   1456»   1479»   1485»   1577»   1597»   378»1454»   1473»   1498»   1495»   1511»   1482»   1520»   1593»   1474»   1454»   1488»   1573»   1450»   1467»   332»1480»   1573»   1490»   1616»   1512»   1488»   1457»   1501»   1458»   1483»   1582»   1519»   1481»   1486»   1489»   1478»   1481»   1486»   1562»   1482»   1453»   1618»   1493»   1464»   1558»   1470»   1582»   1584»   1598»   392»1515»   1514»   1509»   1515»   1482»   1522»   1482»   1499»   1596»   1577» |
| 91 |  |  |  | 91 |  |  |  | 91 |  |  |

Araxis Merge (but not the data content of this report) is Copyright © 1993–2024 Araxis Ltd (www.araxis.com). All rights reserved.
