## Supplementary material for "MTBseq-nf: Enabling Scalable Tuberculosis Genomics “Big Data” Analysis through a User-Friendly Nextflow Wrapper for MTBseq pipeline": SD-6 Intra-modal analysis, with 3-way HTML diff reports generated by Araxis merge software: SD-6-06-intra-modal-araxiscompare-pub-90samples-mtbseq-nf-runs-cluster-groups.html

xml version="1.0" encoding="utf-8"?


Araxis Merge File Comparison Report 

### **Araxis Merge File Comparison Report**

Produced by **Araxis Merge** on **2024/12/18, 16:32 GMT+02:00**. See www.araxis.com for information about Merge. This report uses XHTML and CSS2, and is best viewed with a modern standards-compliant browser. For optimum results when printing this report, use landscape orientation and enable printing of background images and colours in your browser.

##### **1. Files compared**

| # | Location | File | Last Modified |
| --- | --- | --- | --- |
| 1 | /Users/abhi/projects/MTBseq-nf/\_resources/publication/manuscript-and-analysis/v2/pub-90samples-mtbseq-nf-run1/tbgroups/Groups | mtbseqnf\_joint\_cf4\_cr4\_fr75\_ph4\_samples90\_amended\_u95\_phylo\_w12\_d12.groups | 2024/09/20, 10:10 GMT+02:00 |
| 2 | /Users/abhi/projects/MTBseq-nf/\_resources/publication/manuscript-and-analysis/v2/pub-90samples-mtbseq-nf-run2/tbgroups/Groups | mtbseqnf\_joint\_cf4\_cr4\_fr75\_ph4\_samples90\_amended\_u95\_phylo\_w12\_d12.groups | 2024/09/21, 12:10 GMT+02:00 |
| 3 | /Users/abhi/projects/MTBseq-nf/\_resources/publication/manuscript-and-analysis/v2/pub-90samples-mtbseq-nf-run3/tbgroups/Groups | mtbseqnf\_joint\_cf4\_cr4\_fr75\_ph4\_samples90\_amended\_u95\_phylo\_w12\_d12.groups | 2024/09/23, 09:06 GMT+02:00 |
| **Note:** Merge considers the second file to be the common ancestor of the others. | | | |

##### **2. Comparison summary**

| Description | Between Files 1 and 2 | | Between Files 2 and 3 | | Relative to Common Ancestor | |
| --- | --- | --- | --- | --- | --- | --- |
|  | Text Blocks | Lines | Text Blocks | Lines | Text Blocks | Lines |
| Unchanged | 4 | 212 | 4 | 208 |  |  |
| Changed | 2 | 14 | 2 | 18 | 4 | 32 |
| Inserted | 1 | 2 | 0 | 0 | 0 | 0 |
| Removed | 0 | 0 | 1 | 2 | 2 | 4 |
| **Note:** An automatic merge would leave 0 conflict(s). | | | | | | |
