## Supplementary material for "MTBseq-nf: Enabling Scalable Tuberculosis Genomics “Big Data” Analysis through a User-Friendly Nextflow Wrapper for MTBseq pipeline": SD-6 Intra-modal analysis, with 3-way HTML diff reports generated by Araxis merge software: SD-6-09-intra-modal-araxiscompare-pub-90samples-mtbseq-nf-parallel-runs-classification.html

xml version="1.0" encoding="utf-8"?


Araxis Merge File Comparison Report 

### **Araxis Merge File Comparison Report**

Produced by **Araxis Merge** on **2024/12/18, 16:56 GMT+02:00**. See www.araxis.com for information about Merge. This report uses XHTML and CSS2, and is best viewed with a modern standards-compliant browser. For optimum results when printing this report, use landscape orientation and enable printing of background images and colours in your browser.

##### **1. Files compared**

| # | Location | File | Last Modified |
| --- | --- | --- | --- |
| 1 | /Users/abhi/projects/MTBseq-nf/\_resources/publication/manuscript-and-analysis/v2/pub-90samples-mtbseq-nf-parallel-run1/tbstrains/Classification | Strain\_Classification.tab | 2024/09/13, 17:17 GMT+02:00 |
| 2 | /Users/abhi/projects/MTBseq-nf/\_resources/publication/manuscript-and-analysis/v2/pub-90samples-mtbseq-nf-parallel-run2/tbstrains/Classification | Strain\_Classification.tab | 2024/09/17, 20:27 GMT+02:00 |
| 3 | /Users/abhi/projects/MTBseq-nf/\_resources/publication/manuscript-and-analysis/v2/pub-90samples-mtbseq-nf-parallel-run3/tbstrains/Classification | Strain\_Classification.tab | 2024/09/18, 17:08 GMT+02:00 |
| **Note:** Merge considers the second file to be the common ancestor of the others. | | | |

##### **2. Comparison summary**

| Description | Between Files 1 and 2 | | Between Files 2 and 3 | | Relative to Common Ancestor | |
| --- | --- | --- | --- | --- | --- | --- |
|  | Text Blocks | Lines | Text Blocks | Lines | Text Blocks | Lines |
| Unchanged | 1 | 2 | 1 | 2 |  |  |
| Changed | 1 | 180 | 1 | 180 | 0 | 0 |
| Inserted | 0 | 0 | 0 | 0 | 0 | 0 |
| Removed | 0 | 0 | 0 | 0 | 0 | 0 |
| **Note:** An automatic merge would leave 1 conflict(s). | | | | | | |

##### **3. Comparison options**

|  |  |
| --- | --- |
| Whitespace | Consecutive whitespace is treated as a single space |
| Character case | Differences in character case are significant |
| Line endings | Differences in line endings (`CR` and `LF` characters) are ignored |
| CR/LF characters | Not shown in the comparison detail |
| Active line-pairing rules |  |

##### **4. Active regular expressions**

No regular expressions were active.

##### **5. Comparison detail**

|  |  |  |  |  |  |  |  |  |  |  |
| --- | --- | --- | --- | --- | --- | --- | --- | --- | --- | --- |
| 1 |  | Date»   SampleID»   LibraryID»  FullID» Homolka species»Homolka lineage»Homolka group»  Quality»Coll lineage (branch)»  Coll lineage\_name (branch)» Coll quality (branch)»  Coll lineage (easy)»Coll lineage\_name (easy)»   Coll quality (easy)»Beijing lineage (easy)» Beijing quality (easy) |  | 1 |  | Date»   SampleID»   LibraryID»  FullID» Homolka species»Homolka lineage»Homolka group»  Quality»Coll lineage (branch)»  Coll lineage\_name (branch)» Coll quality (branch)»  Coll lineage (easy)»Coll lineage\_name (easy)»   Coll quality (easy)»Beijing lineage (easy)» Beijing quality (easy) |  | 1 |  | Date»   SampleID»   LibraryID»  FullID» Homolka species»Homolka lineage»Homolka group»  Quality»Coll lineage (branch)»  Coll lineage\_name (branch)» Coll quality (branch)»  Coll lineage (easy)»Coll lineage\_name (easy)»   Coll quality (easy)»Beijing lineage (easy)» Beijing quality (easy) |
| 2 |  | '2024-09-13»'10010-03»  'lib951»'10010-03\_lib951»   'M. africanum»  '5» 'West African 1a»   'good»  '5» 'West-Africa 1» 'bad»   '5» 'West-Africa 1» 'bad»   'unknown»   'good |  | 2 |  | '2024-09-17»'10010-03»  'lib951»'10010-03\_lib951»   'M. africanum»  '5» 'West African 1a»   'good»  '5» 'West-Africa 1» 'bad»   '5» 'West-Africa 1» 'bad»   'unknown»   'good |  | 2 |  | '2024-09-18»'10010-03»  'lib951»'10010-03\_lib951»   'M. africanum»  '5» 'West African 1a»   'good»  '5» 'West-Africa 1» 'bad»   '5» 'West-Africa 1» 'bad»   'unknown»   'good |
| 3 |  | '2024-09-13»'10011-03»  'lib914»'10011-03\_lib914»   'M. tuberculosis»   '4.3»   'LAM»   'good»  '4.3.3» 'LAM»   'good»  '4.3.3» 'LAM»   'good»  'unknown»   'good |  | 3 |  | '2024-09-17»'10011-03»  'lib914»'10011-03\_lib914»   'M. tuberculosis»   '4.3»   'LAM»   'good»  '4.3.3» 'LAM»   'good»  '4.3.3» 'LAM»   'good»  'unknown»   'good |  | 3 |  | '2024-09-18»'10011-03»  'lib914»'10011-03\_lib914»   'M. tuberculosis»   '4.3»   'LAM»   'good»  '4.3.3» 'LAM»   'good»  '4.3.3» 'LAM»   'good»  'unknown»   'good |
| 4 |  | '2024-09-13»'10012-03»  'lib970»'10012-03\_lib970»   'M. tuberculosis»   'unknown»   'Clade 1»   'good»  '4.1»   'Euro-American» 'good»  '4.1»   'Euro-American» 'good»  'unknown»   'good |  | 4 |  | '2024-09-17»'10012-03»  'lib970»'10012-03\_lib970»   'M. tuberculosis»   'unknown»   'Clade 1»   'good»  '4.1»   'Euro-American» 'good»  '4.1»   'Euro-American» 'good»  'unknown»   'good |  | 4 |  | '2024-09-18»'10012-03»  'lib970»'10012-03\_lib970»   'M. tuberculosis»   'unknown»   'Clade 1»   'good»  '4.1»   'Euro-American» 'good»  '4.1»   'Euro-American» 'good»  'unknown»   'good |
| 5 |  | '2024-09-13»'10205-03»  'lib915»'10205-03\_lib915»   'M. tuberculosis»   '4.3»   'LAM»   'good»  '4.3.3» 'LAM»   'bad»   '4.3.3» 'LAM»   'bad»   'unknown»   'bad |  | 5 |  | '2024-09-17»'10205-03»  'lib915»'10205-03\_lib915»   'M. tuberculosis»   '4.3»   'LAM»   'good»  '4.3.3» 'LAM»   'bad»   '4.3.3» 'LAM»   'bad»   'unknown»   'bad |  | 5 |  | '2024-09-18»'10205-03»  'lib915»'10205-03\_lib915»   'M. tuberculosis»   '4.3»   'LAM»   'good»  '4.3.3» 'LAM»   'bad»   '4.3.3» 'LAM»   'bad»   'unknown»   'bad |
| 6 |  | '2024-09-13»'10206-03»  'lib916»'10206-03\_lib916»   'M. tuberculosis»   '4.3»   'LAM»   'good»  '4.3.3» 'LAM»   'good»  '4.3.3» 'LAM»   'good»  'unknown»   'good |  | 6 |  | '2024-09-17»'10206-03»  'lib916»'10206-03\_lib916»   'M. tuberculosis»   '4.3»   'LAM»   'good»  '4.3.3» 'LAM»   'good»  '4.3.3» 'LAM»   'good»  'unknown»   'good |  | 6 |  | '2024-09-18»'10206-03»  'lib916»'10206-03\_lib916»   'M. tuberculosis»   '4.3»   'LAM»   'good»  '4.3.3» 'LAM»   'good»  '4.3.3» 'LAM»   'good»  'unknown»   'good |
| 7 |  | '2024-09-13»'10207-03»  'lib973»'10207-03\_lib973»   'M. tuberculosis»   '4.6.2.2»   'Cameroon»  'good»  '4.6.2.2»   'Cameroon»  'good»  '4.6.2.2»   'Cameroon»  'good»  'unknown»   'good |  | 7 |  | '2024-09-17»'10207-03»  'lib973»'10207-03\_lib973»   'M. tuberculosis»   '4.6.2.2»   'Cameroon»  'good»  '4.6.2.2»   'Cameroon»  'good»  '4.6.2.2»   'Cameroon»  'good»  'unknown»   'good |  | 7 |  | '2024-09-18»'10207-03»  'lib973»'10207-03\_lib973»   'M. tuberculosis»   '4.6.2.2»   'Cameroon»  'good»  '4.6.2.2»   'Cameroon»  'good»  '4.6.2.2»   'Cameroon»  'good»  'unknown»   'good |
| 8 |  | '2024-09-13»'10208-03»  'lib1613»   '10208-03\_lib1613»  'M. africanum»  '6» 'West African 2»'good»  '6» 'West-Africa 2» 'good»  '6» 'West-Africa 2» 'good»  'unknown»   'good |  | 8 |  | '2024-09-17»'10208-03»  'lib1613»   '10208-03\_lib1613»  'M. africanum»  '6» 'West African 2»'good»  '6» 'West-Africa 2» 'good»  '6» 'West-Africa 2» 'good»  'unknown»   'good |  | 8 |  | '2024-09-18»'10208-03»  'lib1613»   '10208-03\_lib1613»  'M. africanum»  '6» 'West African 2»'good»  '6» 'West-Africa 2» 'good»  '6» 'West-Africa 2» 'good»  'unknown»   'good |
| 9 |  | '2024-09-13»'10348-03»  'lib917»'10348-03\_lib917»   'M. tuberculosis»   '4.3»   'LAM»   'good»  '4.3.4.2»   'LAM»   'good»  '4.3.4.2»   'LAM»   'good»  'unknown»   'good |  | 9 |  | '2024-09-17»'10348-03»  'lib917»'10348-03\_lib917»   'M. tuberculosis»   '4.3»   'LAM»   'good»  '4.3.4.2»   'LAM»   'good»  '4.3.4.2»   'LAM»   'good»  'unknown»   'good |  | 9 |  | '2024-09-18»'10348-03»  'lib917»'10348-03\_lib917»   'M. tuberculosis»   '4.3»   'LAM»   'good»  '4.3.4.2»   'LAM»   'good»  '4.3.4.2»   'LAM»   'good»  'unknown»   'good |
| 10 |  | '2024-09-13»'10349-03»  'lib924»'10349-03\_lib924»   'M. tuberculosis»   'unknown»   'Clade 1»   'good»  '4.1»   'Euro-American» 'bad»   '4.1»   'Euro-American» 'bad»   'unknown»   'bad |  | 10 |  | '2024-09-17»'10349-03»  'lib924»'10349-03\_lib924»   'M. tuberculosis»   'unknown»   'Clade 1»   'good»  '4.1»   'Euro-American» 'bad»   '4.1»   'Euro-American» 'bad»   'unknown»   'bad |  | 10 |  | '2024-09-18»'10349-03»  'lib924»'10349-03\_lib924»   'M. tuberculosis»   'unknown»   'Clade 1»   'good»  '4.1»   'Euro-American» 'bad»   '4.1»   'Euro-American» 'bad»   'unknown»   'bad |
| 11 |  | '2024-09-13»'10350-03»  'lib1470»   '10350-03\_lib1470»  'M. tuberculosis»   '4.1.2.1»   'Haarlem»   'good»  '4.1.2.1»   'Haarlem»   'good»  '4.1.2.1»   'Haarlem»   'good»  'unknown»   'good |  | 11 |  | '2024-09-17»'10350-03»  'lib1470»   '10350-03\_lib1470»  'M. tuberculosis»   '4.1.2.1»   'Haarlem»   'good»  '4.1.2.1»   'Haarlem»   'good»  '4.1.2.1»   'Haarlem»   'good»  'unknown»   'good |  | 11 |  | '2024-09-18»'10350-03»  'lib1470»   '10350-03\_lib1470»  'M. tuberculosis»   '4.1.2.1»   'Haarlem»   'good»  '4.1.2.1»   'Haarlem»   'good»  '4.1.2.1»   'Haarlem»   'good»  'unknown»   'good |
| 12 |  | '2024-09-13»'10517-03»  'lib939»'10517-03\_lib939»   'M. tuberculosis»   'unknown»   'Clade 1»   'good»  '4.8»   'mainly T»  'good»  '4.8»   'mainly T»  'good»  'unknown»   'good |  | 12 |  | '2024-09-17»'10517-03»  'lib939»'10517-03\_lib939»   'M. tuberculosis»   'unknown»   'Clade 1»   'good»  '4.8»   'mainly T»  'good»  '4.8»   'mainly T»  'good»  'unknown»   'good |  | 12 |  | '2024-09-18»'10517-03»  'lib939»'10517-03\_lib939»   'M. tuberculosis»   'unknown»   'Clade 1»   'good»  '4.8»   'mainly T»  'good»  '4.8»   'mainly T»  'good»  'unknown»   'good |
| 13 |  | '2024-09-13»'11096-03»  'lib940»'11096-03\_lib940»   'M. tuberculosis»   'unknown»   'Clade 1»   'good»  '4.8»   'mainly T»  'good»  '4.8»   'mainly T»  'good»  'unknown»   'good |  | 13 |  | '2024-09-17»'11096-03»  'lib940»'11096-03\_lib940»   'M. tuberculosis»   'unknown»   'Clade 1»   'good»  '4.8»   'mainly T»  'good»  '4.8»   'mainly T»  'good»  'unknown»   'good |  | 13 |  | '2024-09-18»'11096-03»  'lib940»'11096-03\_lib940»   'M. tuberculosis»   'unknown»   'Clade 1»   'good»  '4.8»   'mainly T»  'good»  '4.8»   'mainly T»  'good»  'unknown»   'good |
| 14 |  | '2024-09-13»'11097-03»  'lib933»'11097-03\_lib933»   'M. tuberculosis»   'unknown»   'Clade 1»   'good»  '4.1»   'Euro-American» 'good»  '4.1»   'Euro-American» 'good»  'unknown»   'good |  | 14 |  | '2024-09-17»'11097-03»  'lib933»'11097-03\_lib933»   'M. tuberculosis»   'unknown»   'Clade 1»   'good»  '4.1»   'Euro-American» 'good»  '4.1»   'Euro-American» 'good»  'unknown»   'good |  | 14 |  | '2024-09-18»'11097-03»  'lib933»'11097-03\_lib933»   'M. tuberculosis»   'unknown»   'Clade 1»   'good»  '4.1»   'Euro-American» 'good»  '4.1»   'Euro-American» 'good»  'unknown»   'good |
| 15 |  | '2024-09-13»'11818-03»  'lib902»'11818-03\_lib902»   'M. tuberculosis»   '4.1.2.1»   'Haarlem»   'good»  '4.1.2.1»   'Haarlem»   'good»  '4.1.2.1»   'Haarlem»   'good»  'unknown»   'good |  | 15 |  | '2024-09-17»'11818-03»  'lib902»'11818-03\_lib902»   'M. tuberculosis»   '4.1.2.1»   'Haarlem»   'good»  '4.1.2.1»   'Haarlem»   'good»  '4.1.2.1»   'Haarlem»   'good»  'unknown»   'good |  | 15 |  | '2024-09-18»'11818-03»  'lib902»'11818-03\_lib902»   'M. tuberculosis»   '4.1.2.1»   'Haarlem»   'good»  '4.1.2.1»   'Haarlem»   'good»  '4.1.2.1»   'Haarlem»   'good»  'unknown»   'good |
| 16 |  | '2024-09-13»'11821-03»  'lib952»'11821-03\_lib952»   'M. africanum»  '5» 'West African 1a»   'good»  '5» 'West-Africa 1» 'bad»   '5» 'West-Africa 1» 'bad»   'unknown»   'good |  | 16 |  | '2024-09-17»'11821-03»  'lib952»'11821-03\_lib952»   'M. africanum»  '5» 'West African 1a»   'good»  '5» 'West-Africa 1» 'bad»   '5» 'West-Africa 1» 'bad»   'unknown»   'good |  | 16 |  | '2024-09-18»'11821-03»  'lib952»'11821-03\_lib952»   'M. africanum»  '5» 'West African 1a»   'good»  '5» 'West-Africa 1» 'bad»   '5» 'West-Africa 1» 'bad»   'unknown»   'good |
| 17 |  | '2024-09-13»'11822-03»  'lib903»'11822-03\_lib903»   'M. tuberculosis»   '4.1.2.1»   'Haarlem»   'good»  '4.1.2.1»   'Haarlem»   'good»  '4.1.2.1»   'Haarlem»   'good»  'unknown»   'good |  | 17 |  | '2024-09-17»'11822-03»  'lib903»'11822-03\_lib903»   'M. tuberculosis»   '4.1.2.1»   'Haarlem»   'good»  '4.1.2.1»   'Haarlem»   'good»  '4.1.2.1»   'Haarlem»   'good»  'unknown»   'good |  | 17 |  | '2024-09-18»'11822-03»  'lib903»'11822-03\_lib903»   'M. tuberculosis»   '4.1.2.1»   'Haarlem»   'good»  '4.1.2.1»   'Haarlem»   'good»  '4.1.2.1»   'Haarlem»   'good»  'unknown»   'good |
| 18 |  | '2024-09-13»'12655-03»  'lib1516»   '12655-03\_lib1516»  'M. tuberculosis»   '4.6.2.2»   'Cameroon»  'good»  '4.6.2.2»   'Cameroon»  'good»  '4.6.2.2»   'Cameroon»  'good»  'unknown»   'good |  | 18 |  | '2024-09-17»'12655-03»  'lib1516»   '12655-03\_lib1516»  'M. tuberculosis»   '4.6.2.2»   'Cameroon»  'good»  '4.6.2.2»   'Cameroon»  'good»  '4.6.2.2»   'Cameroon»  'good»  'unknown»   'good |  | 18 |  | '2024-09-18»'12655-03»  'lib1516»   '12655-03\_lib1516»  'M. tuberculosis»   '4.6.2.2»   'Cameroon»  'good»  '4.6.2.2»   'Cameroon»  'good»  '4.6.2.2»   'Cameroon»  'good»  'unknown»   'good |
| 19 |  | '2024-09-13»'12657-03»  'lib934»'12657-03\_lib934»   'M. tuberculosis»   'unknown»   'Clade 1»   'good»  '4.1»   'Euro-American» 'good»  '4.1»   'Euro-American» 'good»  'unknown»   'good |  | 19 |  | '2024-09-17»'12657-03»  'lib934»'12657-03\_lib934»   'M. tuberculosis»   'unknown»   'Clade 1»   'good»  '4.1»   'Euro-American» 'good»  '4.1»   'Euro-American» 'good»  'unknown»   'good |  | 19 |  | '2024-09-18»'12657-03»  'lib934»'12657-03\_lib934»   'M. tuberculosis»   'unknown»   'Clade 1»   'good»  '4.1»   'Euro-American» 'good»  '4.1»   'Euro-American» 'good»  'unknown»   'good |
| 20 |  | '2024-09-13»'12658-03»  'lib1469»   '12658-03\_lib1469»  'M. tuberculosis»   '2» 'Beijing»   'good»  '2.2.1» 'Beijing»   'good»  '2.2.1» 'Beijing»   'good»  'Ancestral 3»   'good |  | 20 |  | '2024-09-17»'12658-03»  'lib1469»   '12658-03\_lib1469»  'M. tuberculosis»   '2» 'Beijing»   'good»  '2.2.1» 'Beijing»   'good»  '2.2.1» 'Beijing»   'good»  'Ancestral 3»   'good |  | 20 |  | '2024-09-18»'12658-03»  'lib1469»   '12658-03\_lib1469»  'M. tuberculosis»   '2» 'Beijing»   'good»  '2.2.1» 'Beijing»   'good»  '2.2.1» 'Beijing»   'good»  'Ancestral 3»   'good |
| 21 |  | '2024-09-13»'1322-04»   'lib925»'1322-04\_lib925»'M. tuberculosis»   'unknown»   'Clade 1»   'good»  '4.8»   'mainly T»  'good»  '4.8»   'mainly T»  'good»  'unknown»   'good |  | 21 |  | '2024-09-17»'1322-04»   'lib925»'1322-04\_lib925»'M. tuberculosis»   'unknown»   'Clade 1»   'good»  '4.8»   'mainly T»  'good»  '4.8»   'mainly T»  'good»  'unknown»   'good |  | 21 |  | '2024-09-18»'1322-04»   'lib925»'1322-04\_lib925»'M. tuberculosis»   'unknown»   'Clade 1»   'good»  '4.8»   'mainly T»  'good»  '4.8»   'mainly T»  'good»  'unknown»   'good |
| 22 |  | '2024-09-13»'1324-04»   'lib931»'1324-04\_lib931»'M. tuberculosis»   '4.4.1.1»   'S-type»'good»  '4.4.1.1»   'S-type»'good»  '4.4.1.1»   'S-type»'good»  'unknown»   'good |  | 22 |  | '2024-09-17»'1324-04»   'lib931»'1324-04\_lib931»'M. tuberculosis»   '4.4.1.1»   'S-type»'good»  '4.4.1.1»   'S-type»'good»  '4.4.1.1»   'S-type»'good»  'unknown»   'good |  | 22 |  | '2024-09-18»'1324-04»   'lib931»'1324-04\_lib931»'M. tuberculosis»   '4.4.1.1»   'S-type»'good»  '4.4.1.1»   'S-type»'good»  '4.4.1.1»   'S-type»'good»  'unknown»   'good |
| 23 |  | '2024-09-13»'1327-04»   'lib957»'1327-04\_lib957»'M. africanum»  '6» 'West African 2»'good»  '6» 'West-Africa 2» 'good»  '6» 'West-Africa 2» 'good»  'unknown»   'good |  | 23 |  | '2024-09-17»'1327-04»   'lib957»'1327-04\_lib957»'M. africanum»  '6» 'West African 2»'good»  '6» 'West-Africa 2» 'good»  '6» 'West-Africa 2» 'good»  'unknown»   'good |  | 23 |  | '2024-09-18»'1327-04»   'lib957»'1327-04\_lib957»'M. africanum»  '6» 'West African 2»'good»  '6» 'West-Africa 2» 'good»  '6» 'West-Africa 2» 'good»  'unknown»   'good |
| 24 |  | '2024-09-13»'1597-04»   'lib918»'1597-04\_lib918»'M. tuberculosis»   '4.3»   'LAM»   'good»  '4.3.3» 'LAM»   'bad»   '4.3.3» 'LAM»   'bad»   'unknown»   'good |  | 24 |  | '2024-09-17»'1597-04»   'lib918»'1597-04\_lib918»'M. tuberculosis»   '4.3»   'LAM»   'good»  '4.3.3» 'LAM»   'bad»   '4.3.3» 'LAM»   'bad»   'unknown»   'good |  | 24 |  | '2024-09-18»'1597-04»   'lib918»'1597-04\_lib918»'M. tuberculosis»   '4.3»   'LAM»   'good»  '4.3.3» 'LAM»   'bad»   '4.3.3» 'LAM»   'bad»   'unknown»   'good |
| 25 |  | '2024-09-13»'1599-04»   'lib974»'1599-04\_lib974»'M. tuberculosis»   '4.1.2.1»   'Haarlem»   'good»  '4.1.2.1»   'Haarlem»   'good»  '4.1.2.1»   'Haarlem»   'good»  'unknown»   'good |  | 25 |  | '2024-09-17»'1599-04»   'lib974»'1599-04\_lib974»'M. tuberculosis»   '4.1.2.1»   'Haarlem»   'good»  '4.1.2.1»   'Haarlem»   'good»  '4.1.2.1»   'Haarlem»   'good»  'unknown»   'good |  | 25 |  | '2024-09-18»'1599-04»   'lib974»'1599-04\_lib974»'M. tuberculosis»   '4.1.2.1»   'Haarlem»   'good»  '4.1.2.1»   'Haarlem»   'good»  '4.1.2.1»   'Haarlem»   'good»  'unknown»   'good |
| 26 |  | '2024-09-13»'1779-04»   'lib941»'1779-04\_lib941»'M. tuberculosis»   'unknown»   'Clade 1»   'good»  '4.8»   'mainly T»  'good»  '4.8»   'mainly T»  'good»  'unknown»   'good |  | 26 |  | '2024-09-17»'1779-04»   'lib941»'1779-04\_lib941»'M. tuberculosis»   'unknown»   'Clade 1»   'good»  '4.8»   'mainly T»  'good»  '4.8»   'mainly T»  'good»  'unknown»   'good |  | 26 |  | '2024-09-18»'1779-04»   'lib941»'1779-04\_lib941»'M. tuberculosis»   'unknown»   'Clade 1»   'good»  '4.8»   'mainly T»  'good»  '4.8»   'mainly T»  'good»  'unknown»   'good |
| 27 |  | '2024-09-13»'1780-04»   'lib942»'1780-04\_lib942»'M. tuberculosis»   'unknown»   'Clade 1»   'good»  '4.8»   'mainly T»  'bad»   '4.8»   'mainly T»  'bad»   'unknown»   'good |  | 27 |  | '2024-09-17»'1780-04»   'lib942»'1780-04\_lib942»'M. tuberculosis»   'unknown»   'Clade 1»   'good»  '4.8»   'mainly T»  'bad»   '4.8»   'mainly T»  'bad»   'unknown»   'good |  | 27 |  | '2024-09-18»'1780-04»   'lib942»'1780-04\_lib942»'M. tuberculosis»   'unknown»   'Clade 1»   'good»  '4.8»   'mainly T»  'bad»   '4.8»   'mainly T»  'bad»   'unknown»   'good |
| 28 |  | '2024-09-13»'1783-04»   'lib936»'1783-04\_lib936»'M. tuberculosis»   'unknown»   'Clade 1»   'good»  '4.1»   'Euro-American» 'good»  '4.1»   'Euro-American» 'good»  'unknown»   'good |  | 28 |  | '2024-09-17»'1783-04»   'lib936»'1783-04\_lib936»'M. tuberculosis»   'unknown»   'Clade 1»   'good»  '4.1»   'Euro-American» 'good»  '4.1»   'Euro-American» 'good»  'unknown»   'good |  | 28 |  | '2024-09-18»'1783-04»   'lib936»'1783-04\_lib936»'M. tuberculosis»   'unknown»   'Clade 1»   'good»  '4.1»   'Euro-American» 'good»  '4.1»   'Euro-American» 'good»  'unknown»   'good |
| 29 |  | '2024-09-13»'2509-04»   'lib1471»   '2509-04\_lib1471»   'M. tuberculosis»   '4.3»   'LAM»   'good»  '4.3.3» 'LAM»   'good»  '4.3.3» 'LAM»   'good»  'unknown»   'good |  | 29 |  | '2024-09-17»'2509-04»   'lib1471»   '2509-04\_lib1471»   'M. tuberculosis»   '4.3»   'LAM»   'good»  '4.3.3» 'LAM»   'good»  '4.3.3» 'LAM»   'good»  'unknown»   'good |  | 29 |  | '2024-09-18»'2509-04»   'lib1471»   '2509-04\_lib1471»   'M. tuberculosis»   '4.3»   'LAM»   'good»  '4.3.3» 'LAM»   'good»  '4.3.3» 'LAM»   'good»  'unknown»   'good |
| 30 |  | '2024-09-13»'3154-04»   'lib905»'3154-04\_lib905»'M. tuberculosis»   '4.1.2.1»   'Haarlem»   'good»  '4.1.2.1»   'Haarlem»   'good»  '4.1.2.1»   'Haarlem»   'good»  'unknown»   'good |  | 30 |  | '2024-09-17»'3154-04»   'lib905»'3154-04\_lib905»'M. tuberculosis»   '4.1.2.1»   'Haarlem»   'good»  '4.1.2.1»   'Haarlem»   'good»  '4.1.2.1»   'Haarlem»   'good»  'unknown»   'good |  | 30 |  | '2024-09-18»'3154-04»   'lib905»'3154-04\_lib905»'M. tuberculosis»   '4.1.2.1»   'Haarlem»   'good»  '4.1.2.1»   'Haarlem»   'good»  '4.1.2.1»   'Haarlem»   'good»  'unknown»   'good |
| 31 |  | '2024-09-13»'3156-04»   'lib926»'3156-04\_lib926»'M. tuberculosis»   'unknown»   'Clade 1»   'good»  '4.1.1.3»   'X-type»'bad»   '4.1.1.3»   'X-type»'bad»   'unknown»   'good |  | 31 |  | '2024-09-17»'3156-04»   'lib926»'3156-04\_lib926»'M. tuberculosis»   'unknown»   'Clade 1»   'good»  '4.1.1.3»   'X-type»'bad»   '4.1.1.3»   'X-type»'bad»   'unknown»   'good |  | 31 |  | '2024-09-18»'3156-04»   'lib926»'3156-04\_lib926»'M. tuberculosis»   'unknown»   'Clade 1»   'good»  '4.1.1.3»   'X-type»'bad»   '4.1.1.3»   'X-type»'bad»   'unknown»   'good |
| 32 |  | '2024-09-13»'3158-04»   'lib943»'3158-04\_lib943»'M. tuberculosis»   'unknown»   'Clade 1»   'good»  '4.8»   'mainly T»  'good»  '4.8»   'mainly T»  'good»  'unknown»   'good |  | 32 |  | '2024-09-17»'3158-04»   'lib943»'3158-04\_lib943»'M. tuberculosis»   'unknown»   'Clade 1»   'good»  '4.8»   'mainly T»  'good»  '4.8»   'mainly T»  'good»  'unknown»   'good |  | 32 |  | '2024-09-18»'3158-04»   'lib943»'3158-04\_lib943»'M. tuberculosis»   'unknown»   'Clade 1»   'good»  '4.8»   'mainly T»  'good»  '4.8»   'mainly T»  'good»  'unknown»   'good |
| 33 |  | '2024-09-13»'3160-04»   'lib958»'3160-04\_lib958»'M. africanum»  '6» 'West African 2»'good»  '6» 'West-Africa 2» 'good»  '6» 'West-Africa 2» 'good»  'unknown»   'good |  | 33 |  | '2024-09-17»'3160-04»   'lib958»'3160-04\_lib958»'M. africanum»  '6» 'West African 2»'good»  '6» 'West-Africa 2» 'good»  '6» 'West-Africa 2» 'good»  'unknown»   'good |  | 33 |  | '2024-09-18»'3160-04»   'lib958»'3160-04\_lib958»'M. africanum»  '6» 'West African 2»'good»  '6» 'West-Africa 2» 'good»  '6» 'West-Africa 2» 'good»  'unknown»   'good |
| 34 |  | '2024-09-13»'3491-04»   'lib897»'3491-04\_lib897»'M. tuberculosis»   '1» 'EAI»   'good»  '1.1.1» 'EAI»   'bad»   '1.1.1» 'EAI»   'bad»   'unknown»   'good |  | 34 |  | '2024-09-17»'3491-04»   'lib897»'3491-04\_lib897»'M. tuberculosis»   '1» 'EAI»   'good»  '1.1.1» 'EAI»   'bad»   '1.1.1» 'EAI»   'bad»   'unknown»   'good |  | 34 |  | '2024-09-18»'3491-04»   'lib897»'3491-04\_lib897»'M. tuberculosis»   '1» 'EAI»   'good»  '1.1.1» 'EAI»   'bad»   '1.1.1» 'EAI»   'bad»   'unknown»   'good |
| 35 |  | '2024-09-13»'3494-04»   'lib953»'3494-04\_lib953»'M. africanum»  '5» 'West African 1a»   'good»  '5» 'West-Africa 1» 'bad»   '5» 'West-Africa 1» 'bad»   'unknown»   'good |  | 35 |  | '2024-09-17»'3494-04»   'lib953»'3494-04\_lib953»'M. africanum»  '5» 'West African 1a»   'good»  '5» 'West-Africa 1» 'bad»   '5» 'West-Africa 1» 'bad»   'unknown»   'good |  | 35 |  | '2024-09-18»'3494-04»   'lib953»'3494-04\_lib953»'M. africanum»  '5» 'West African 1a»   'good»  '5» 'West-Africa 1» 'bad»   '5» 'West-Africa 1» 'bad»   'unknown»   'good |
| 36 |  | '2024-09-13»'3496-04»   'lib906»'3496-04\_lib906»'M. tuberculosis»   '4.1.2.1»   'Haarlem»   'good»  '4.1.2.1»   'Haarlem»   'good»  '4.1.2.1»   'Haarlem»   'good»  'unknown»   'good |  | 36 |  | '2024-09-17»'3496-04»   'lib906»'3496-04\_lib906»'M. tuberculosis»   '4.1.2.1»   'Haarlem»   'good»  '4.1.2.1»   'Haarlem»   'good»  '4.1.2.1»   'Haarlem»   'good»  'unknown»   'good |  | 36 |  | '2024-09-18»'3496-04»   'lib906»'3496-04\_lib906»'M. tuberculosis»   '4.1.2.1»   'Haarlem»   'good»  '4.1.2.1»   'Haarlem»   'good»  '4.1.2.1»   'Haarlem»   'good»  'unknown»   'good |
| 37 |  | '2024-09-13»'3497-04»   'lib927»'3497-04\_lib927»'M. tuberculosis»   'unknown»   'Clade 1»   'good»  '4.8»   'mainly T»  'bad»   '4.8»   'mainly T»  'bad»   'unknown»   'good |  | 37 |  | '2024-09-17»'3497-04»   'lib927»'3497-04\_lib927»'M. tuberculosis»   'unknown»   'Clade 1»   'good»  '4.8»   'mainly T»  'bad»   '4.8»   'mainly T»  'bad»   'unknown»   'good |  | 37 |  | '2024-09-18»'3497-04»   'lib927»'3497-04\_lib927»'M. tuberculosis»   'unknown»   'Clade 1»   'good»  '4.8»   'mainly T»  'bad»   '4.8»   'mainly T»  'bad»   'unknown»   'good |
| 38 |  | '2024-09-13»'3734-04»   'lib895»'3734-04\_lib895»'M. tuberculosis»   '4.6.2.2»   'Cameroon»  'good»  '4.6.2.2»   'Cameroon»  'good»  '4.6.2.2»   'Cameroon»  'good»  'unknown»   'good |  | 38 |  | '2024-09-17»'3734-04»   'lib895»'3734-04\_lib895»'M. tuberculosis»   '4.6.2.2»   'Cameroon»  'good»  '4.6.2.2»   'Cameroon»  'good»  '4.6.2.2»   'Cameroon»  'good»  'unknown»   'good |  | 38 |  | '2024-09-18»'3734-04»   'lib895»'3734-04\_lib895»'M. tuberculosis»   '4.6.2.2»   'Cameroon»  'good»  '4.6.2.2»   'Cameroon»  'good»  '4.6.2.2»   'Cameroon»  'good»  'unknown»   'good |
| 39 |  | '2024-09-13»'3736-04»   'lib937»'3736-04\_lib937»'M. tuberculosis»   'unknown»   'Clade 1»   'good»  '4.1»   'Euro-American» 'good»  '4.1»   'Euro-American» 'good»  'unknown»   'good |  | 39 |  | '2024-09-17»'3736-04»   'lib937»'3736-04\_lib937»'M. tuberculosis»   'unknown»   'Clade 1»   'good»  '4.1»   'Euro-American» 'good»  '4.1»   'Euro-American» 'good»  'unknown»   'good |  | 39 |  | '2024-09-18»'3736-04»   'lib937»'3736-04\_lib937»'M. tuberculosis»   'unknown»   'Clade 1»   'good»  '4.1»   'Euro-American» 'good»  '4.1»   'Euro-American» 'good»  'unknown»   'good |
| 40 |  | '2024-09-13»'3859-03»   'lib921»'3859-03\_lib921»'M. tuberculosis»   'unknown»   'Clade 1»   'good»  '4.1»   'Euro-American» 'good»  '4.1»   'Euro-American» 'good»  'unknown»   'good |  | 40 |  | '2024-09-17»'3859-03»   'lib921»'3859-03\_lib921»'M. tuberculosis»   'unknown»   'Clade 1»   'good»  '4.1»   'Euro-American» 'good»  '4.1»   'Euro-American» 'good»  'unknown»   'good |  | 40 |  | '2024-09-18»'3859-03»   'lib921»'3859-03\_lib921»'M. tuberculosis»   'unknown»   'Clade 1»   'good»  '4.1»   'Euro-American» 'good»  '4.1»   'Euro-American» 'good»  'unknown»   'good |
| 41 |  | '2024-09-13»'3861-03»   'lib922»'3861-03\_lib922»'M. tuberculosis»   'unknown»   'Clade 1»   'good»  '4.6»   'Euro-American» 'good»  '4.6»   'Euro-American» 'good»  'unknown»   'good |  | 41 |  | '2024-09-17»'3861-03»   'lib922»'3861-03\_lib922»'M. tuberculosis»   'unknown»   'Clade 1»   'good»  '4.6»   'Euro-American» 'good»  '4.6»   'Euro-American» 'good»  'unknown»   'good |  | 41 |  | '2024-09-18»'3861-03»   'lib922»'3861-03\_lib922»'M. tuberculosis»   'unknown»   'Clade 1»   'good»  '4.6»   'Euro-American» 'good»  '4.6»   'Euro-American» 'good»  'unknown»   'good |
| 42 |  | '2024-09-13»'3865-03»   'lib971»'3865-03\_lib971»'M. tuberculosis»   '4.3»   'LAM»   'good»  '4.3.3» 'LAM»   'good»  '4.3.3» 'LAM»   'good»  'unknown»   'good |  | 42 |  | '2024-09-17»'3865-03»   'lib971»'3865-03\_lib971»'M. tuberculosis»   '4.3»   'LAM»   'good»  '4.3.3» 'LAM»   'good»  '4.3.3» 'LAM»   'good»  'unknown»   'good |  | 42 |  | '2024-09-18»'3865-03»   'lib971»'3865-03\_lib971»'M. tuberculosis»   '4.3»   'LAM»   'good»  '4.3.3» 'LAM»   'good»  '4.3.3» 'LAM»   'good»  'unknown»   'good |
| 43 |  | '2024-09-13»'4139-04»   'lib959»'4139-04\_lib959»'M. africanum»  '6» 'West African 2»'good»  '6» 'West-Africa 2» 'good»  '6» 'West-Africa 2» 'good»  'unknown»   'good |  | 43 |  | '2024-09-17»'4139-04»   'lib959»'4139-04\_lib959»'M. africanum»  '6» 'West African 2»'good»  '6» 'West-Africa 2» 'good»  '6» 'West-Africa 2» 'good»  'unknown»   'good |  | 43 |  | '2024-09-18»'4139-04»   'lib959»'4139-04\_lib959»'M. africanum»  '6» 'West African 2»'good»  '6» 'West-Africa 2» 'good»  '6» 'West-Africa 2» 'good»  'unknown»   'good |
| 44 |  | '2024-09-13»'4145-04»   'lib919»'4145-04\_lib919»'M. tuberculosis»   '4.3»   'LAM»   'good»  '4.3.3» 'LAM»   'good»  '4.3.3» 'LAM»   'good»  'unknown»   'good |  | 44 |  | '2024-09-17»'4145-04»   'lib919»'4145-04\_lib919»'M. tuberculosis»   '4.3»   'LAM»   'good»  '4.3.3» 'LAM»   'good»  '4.3.3» 'LAM»   'good»  'unknown»   'good |  | 44 |  | '2024-09-18»'4145-04»   'lib919»'4145-04\_lib919»'M. tuberculosis»   '4.3»   'LAM»   'good»  '4.3.3» 'LAM»   'good»  '4.3.3» 'LAM»   'good»  'unknown»   'good |
| 45 |  | '2024-09-13»'4148-04»   'lib907»'4148-04\_lib907»'M. tuberculosis»   '4.1.2.1»   'Haarlem»   'good»  '4.1.2.1»   'Haarlem»   'good»  '4.1.2.1»   'Haarlem»   'good»  'unknown»   'good |  | 45 |  | '2024-09-17»'4148-04»   'lib907»'4148-04\_lib907»'M. tuberculosis»   '4.1.2.1»   'Haarlem»   'good»  '4.1.2.1»   'Haarlem»   'good»  '4.1.2.1»   'Haarlem»   'good»  'unknown»   'good |  | 45 |  | '2024-09-18»'4148-04»   'lib907»'4148-04\_lib907»'M. tuberculosis»   '4.1.2.1»   'Haarlem»   'good»  '4.1.2.1»   'Haarlem»   'good»  '4.1.2.1»   'Haarlem»   'good»  'unknown»   'good |
| 46 |  | '2024-09-13»'420-04»'lib935»'420-04\_lib935» 'M. tuberculosis»   'unknown»   'Clade 1»   'good»  '4.1»   'Euro-American» 'good»  '4.1»   'Euro-American» 'good»  'unknown»   'good |  | 46 |  | '2024-09-17»'420-04»'lib935»'420-04\_lib935» 'M. tuberculosis»   'unknown»   'Clade 1»   'good»  '4.1»   'Euro-American» 'good»  '4.1»   'Euro-American» 'good»  'unknown»   'good |  | 46 |  | '2024-09-18»'420-04»'lib935»'420-04\_lib935» 'M. tuberculosis»   'unknown»   'Clade 1»   'good»  '4.1»   'Euro-American» 'good»  '4.1»   'Euro-American» 'good»  'unknown»   'good |
| 47 |  | '2024-09-13»'421-04»'lib956»'421-04\_lib956» 'M. africanum»  '6» 'West African 2»'good»  '6» 'West-Africa 2» 'good»  '6» 'West-Africa 2» 'good»  'unknown»   'good |  | 47 |  | '2024-09-17»'421-04»'lib956»'421-04\_lib956» 'M. africanum»  '6» 'West African 2»'good»  '6» 'West-Africa 2» 'good»  '6» 'West-Africa 2» 'good»  'unknown»   'good |  | 47 |  | '2024-09-18»'421-04»'lib956»'421-04\_lib956» 'M. africanum»  '6» 'West African 2»'good»  '6» 'West-Africa 2» 'good»  '6» 'West-Africa 2» 'good»  'unknown»   'good |
| 48 |  | '2024-09-13»'4514-03»   'lib901»'4514-03\_lib901»'M. tuberculosis»   '4.1.2.1»   'Haarlem»   'bad»   '4.1.2.1»   'Haarlem»   'bad»   '4.1.2.1»   'Haarlem»   'bad»   'unknown»   'bad |  | 48 |  | '2024-09-17»'4514-03»   'lib901»'4514-03\_lib901»'M. tuberculosis»   '4.1.2.1»   'Haarlem»   'bad»   '4.1.2.1»   'Haarlem»   'bad»   '4.1.2.1»   'Haarlem»   'bad»   'unknown»   'bad |  | 48 |  | '2024-09-18»'4514-03»   'lib901»'4514-03\_lib901»'M. tuberculosis»   '4.1.2.1»   'Haarlem»   'bad»   '4.1.2.1»   'Haarlem»   'bad»   '4.1.2.1»   'Haarlem»   'bad»   'unknown»   'bad |
| 49 |  | '2024-09-13»'4516-03»   'lib947»'4516-03\_lib947»'M. tuberculosis»   'unknown»   'Clade 1»   'good»  '4.1.1.1»   'X-type»'good»  '4.1.1.1»   'X-type»'good»  'unknown»   'good |  | 49 |  | '2024-09-17»'4516-03»   'lib947»'4516-03\_lib947»'M. tuberculosis»   'unknown»   'Clade 1»   'good»  '4.1.1.1»   'X-type»'good»  '4.1.1.1»   'X-type»'good»  'unknown»   'good |  | 49 |  | '2024-09-18»'4516-03»   'lib947»'4516-03\_lib947»'M. tuberculosis»   'unknown»   'Clade 1»   'good»  '4.1.1.1»   'X-type»'good»  '4.1.1.1»   'X-type»'good»  'unknown»   'good |
| 50 |  | '2024-09-13»'4518-03»   'lib949»'4518-03\_lib949»'M. africanum»  '5» 'West African 1a»   'good»  '5» 'West-Africa 1» 'bad»   '5» 'West-Africa 1» 'bad»   'unknown»   'good |  | 50 |  | '2024-09-17»'4518-03»   'lib949»'4518-03\_lib949»'M. africanum»  '5» 'West African 1a»   'good»  '5» 'West-Africa 1» 'bad»   '5» 'West-Africa 1» 'bad»   'unknown»   'good |  | 50 |  | '2024-09-18»'4518-03»   'lib949»'4518-03\_lib949»'M. africanum»  '5» 'West African 1a»   'good»  '5» 'West-Africa 1» 'bad»   '5» 'West-Africa 1» 'bad»   'unknown»   'good |
| 51 |  | '2024-09-13»'4523-03»   'lib972»'4523-03\_lib972»'M. tuberculosis»   '4.3»   'LAM»   'good»  '4.3.3» 'LAM»   'good»  '4.3.3» 'LAM»   'good»  'unknown»   'good |  | 51 |  | '2024-09-17»'4523-03»   'lib972»'4523-03\_lib972»'M. tuberculosis»   '4.3»   'LAM»   'good»  '4.3.3» 'LAM»   'good»  '4.3.3» 'LAM»   'good»  'unknown»   'good |  | 51 |  | '2024-09-18»'4523-03»   'lib972»'4523-03\_lib972»'M. tuberculosis»   '4.3»   'LAM»   'good»  '4.3.3» 'LAM»   'good»  '4.3.3» 'LAM»   'good»  'unknown»   'good |
| 52 |  | '2024-09-13»'4712-04»   'lib960»'4712-04\_lib960»'M. africanum»  '6» 'West African 2»'good»  '6» 'West-Africa 2» 'good»  '6» 'West-Africa 2» 'good»  'unknown»   'good |  | 52 |  | '2024-09-17»'4712-04»   'lib960»'4712-04\_lib960»'M. africanum»  '6» 'West African 2»'good»  '6» 'West-Africa 2» 'good»  '6» 'West-Africa 2» 'good»  'unknown»   'good |  | 52 |  | '2024-09-18»'4712-04»   'lib960»'4712-04\_lib960»'M. africanum»  '6» 'West African 2»'good»  '6» 'West-Africa 2» 'good»  '6» 'West-Africa 2» 'good»  'unknown»   'good |
| 53 |  | '2024-09-13»'4714-04»   'lib1472»   '4714-04\_lib1472»   'M. tuberculosis»   '4.4.1.1»   'S-type»'good»  '4.4.1.1»   'S-type»'good»  '4.4.1.1»   'S-type»'good»  'unknown»   'good |  | 53 |  | '2024-09-17»'4714-04»   'lib1472»   '4714-04\_lib1472»   'M. tuberculosis»   '4.4.1.1»   'S-type»'good»  '4.4.1.1»   'S-type»'good»  '4.4.1.1»   'S-type»'good»  'unknown»   'good |  | 53 |  | '2024-09-18»'4714-04»   'lib1472»   '4714-04\_lib1472»   'M. tuberculosis»   '4.4.1.1»   'S-type»'good»  '4.4.1.1»   'S-type»'good»  '4.4.1.1»   'S-type»'good»  'unknown»   'good |
| 54 |  | '2024-09-13»'4717-04»   'lib898»'4717-04\_lib898»'M. tuberculosis»   '1» 'EAI»   'good»  '1.1.1» 'EAI»   'bad»   '1.1.1» 'EAI»   'bad»   'unknown»   'good |  | 54 |  | '2024-09-17»'4717-04»   'lib898»'4717-04\_lib898»'M. tuberculosis»   '1» 'EAI»   'good»  '1.1.1» 'EAI»   'bad»   '1.1.1» 'EAI»   'bad»   'unknown»   'good |  | 54 |  | '2024-09-18»'4717-04»   'lib898»'4717-04\_lib898»'M. tuberculosis»   '1» 'EAI»   'good»  '1.1.1» 'EAI»   'bad»   '1.1.1» 'EAI»   'bad»   'unknown»   'good |
| 55 |  | '2024-09-13»'4724-03»   'lib1517»   '4724-03\_lib1517»   'M. tuberculosis»   '2» 'Beijing»   'good»  '2.2.1» 'Beijing»   'good»  '2.2.1» 'Beijing»   'good»  'Ancestral 3»   'good |  | 55 |  | '2024-09-17»'4724-03»   'lib1517»   '4724-03\_lib1517»   'M. tuberculosis»   '2» 'Beijing»   'good»  '2.2.1» 'Beijing»   'good»  '2.2.1» 'Beijing»   'good»  'Ancestral 3»   'good |  | 55 |  | '2024-09-18»'4724-03»   'lib1517»   '4724-03\_lib1517»   'M. tuberculosis»   '2» 'Beijing»   'good»  '2.2.1» 'Beijing»   'good»  '2.2.1» 'Beijing»   'good»  'Ancestral 3»   'good |
| 56 |  | '2024-09-13»'4779-04»   'lib944»'4779-04\_lib944»'M. tuberculosis»   'unknown»   'Clade 1»   'good»  '4.8»   'mainly T»  'good»  '4.8»   'mainly T»  'good»  'unknown»   'good |  | 56 |  | '2024-09-17»'4779-04»   'lib944»'4779-04\_lib944»'M. tuberculosis»   'unknown»   'Clade 1»   'good»  '4.8»   'mainly T»  'good»  '4.8»   'mainly T»  'good»  'unknown»   'good |  | 56 |  | '2024-09-18»'4779-04»   'lib944»'4779-04\_lib944»'M. tuberculosis»   'unknown»   'Clade 1»   'good»  '4.8»   'mainly T»  'good»  '4.8»   'mainly T»  'good»  'unknown»   'good |
| 57 |  | '2024-09-13»'4781-04»   'lib908»'4781-04\_lib908»'M. tuberculosis»   '4.1.2.1»   'Haarlem»   'good»  '4.1.2.1»   'Haarlem»   'good»  '4.1.2.1»   'Haarlem»   'good»  'unknown»   'good |  | 57 |  | '2024-09-17»'4781-04»   'lib908»'4781-04\_lib908»'M. tuberculosis»   '4.1.2.1»   'Haarlem»   'good»  '4.1.2.1»   'Haarlem»   'good»  '4.1.2.1»   'Haarlem»   'good»  'unknown»   'good |  | 57 |  | '2024-09-18»'4781-04»   'lib908»'4781-04\_lib908»'M. tuberculosis»   '4.1.2.1»   'Haarlem»   'good»  '4.1.2.1»   'Haarlem»   'good»  '4.1.2.1»   'Haarlem»   'good»  'unknown»   'good |
| 58 |  | '2024-09-13»'4783-04»   'lib896»'4783-04\_lib896»'M. tuberculosis»   '4.6.2.2»   'Cameroon»  'good»  '4.6.2.2»   'Cameroon»  'good»  '4.6.2.2»   'Cameroon»  'good»  'unknown»   'good |  | 58 |  | '2024-09-17»'4783-04»   'lib896»'4783-04\_lib896»'M. tuberculosis»   '4.6.2.2»   'Cameroon»  'good»  '4.6.2.2»   'Cameroon»  'good»  '4.6.2.2»   'Cameroon»  'good»  'unknown»   'good |  | 58 |  | '2024-09-18»'4783-04»   'lib896»'4783-04\_lib896»'M. tuberculosis»   '4.6.2.2»   'Cameroon»  'good»  '4.6.2.2»   'Cameroon»  'good»  '4.6.2.2»   'Cameroon»  'good»  'unknown»   'good |
| 59 |  | '2024-09-13»'4785-04»   'lib909»'4785-04\_lib909»'M. tuberculosis»   '4.1.2.1»   'Haarlem»   'good»  '4.1.2.1»   'Haarlem»   'good»  '4.1.2.1»   'Haarlem»   'good»  'unknown»   'good |  | 59 |  | '2024-09-17»'4785-04»   'lib909»'4785-04\_lib909»'M. tuberculosis»   '4.1.2.1»   'Haarlem»   'good»  '4.1.2.1»   'Haarlem»   'good»  '4.1.2.1»   'Haarlem»   'good»  'unknown»   'good |  | 59 |  | '2024-09-18»'4785-04»   'lib909»'4785-04\_lib909»'M. tuberculosis»   '4.1.2.1»   'Haarlem»   'good»  '4.1.2.1»   'Haarlem»   'good»  '4.1.2.1»   'Haarlem»   'good»  'unknown»   'good |
| 60 |  | '2024-09-13»'5248-04»   'lib928»'5248-04\_lib928»'M. tuberculosis»   'unknown»   'Clade 1»   'good»  '4.1.1.3»   'X-type»'good»  '4.1.1.3»   'X-type»'good»  'unknown»   'good |  | 60 |  | '2024-09-17»'5248-04»   'lib928»'5248-04\_lib928»'M. tuberculosis»   'unknown»   'Clade 1»   'good»  '4.1.1.3»   'X-type»'good»  '4.1.1.3»   'X-type»'good»  'unknown»   'good |  | 60 |  | '2024-09-18»'5248-04»   'lib928»'5248-04\_lib928»'M. tuberculosis»   'unknown»   'Clade 1»   'good»  '4.1.1.3»   'X-type»'good»  '4.1.1.3»   'X-type»'good»  'unknown»   'good |
| 61 |  | '2024-09-13»'5253-04»   'lib961»'5253-04\_lib961»'M. africanum»  '6» 'West African 2»'good»  '6» 'West-Africa 2» 'good»  '6» 'West-Africa 2» 'good»  'unknown»   'good |  | 61 |  | '2024-09-17»'5253-04»   'lib961»'5253-04\_lib961»'M. africanum»  '6» 'West African 2»'good»  '6» 'West-Africa 2» 'good»  '6» 'West-Africa 2» 'good»  'unknown»   'good |  | 61 |  | '2024-09-18»'5253-04»   'lib961»'5253-04\_lib961»'M. africanum»  '6» 'West African 2»'good»  '6» 'West-Africa 2» 'good»  '6» 'West-Africa 2» 'good»  'unknown»   'good |
| 62 |  | '2024-09-13»'5468-03»   'lib912»'5468-03\_lib912»'M. tuberculosis»   '4.3»   'LAM»   'good»  '4.3.3» 'LAM»   'good»  '4.3.3» 'LAM»   'good»  'unknown»   'good |  | 62 |  | '2024-09-17»'5468-03»   'lib912»'5468-03\_lib912»'M. tuberculosis»   '4.3»   'LAM»   'good»  '4.3.3» 'LAM»   'good»  '4.3.3» 'LAM»   'good»  'unknown»   'good |  | 62 |  | '2024-09-18»'5468-03»   'lib912»'5468-03\_lib912»'M. tuberculosis»   '4.3»   'LAM»   'good»  '4.3.3» 'LAM»   'good»  '4.3.3» 'LAM»   'good»  'unknown»   'good |
| 63 |  | '2024-09-13»'5472-03»   'lib938»'5472-03\_lib938»'M. tuberculosis»   'unknown»   'Clade 1»   'good»  '4.8»   'mainly T»  'good»  '4.8»   'mainly T»  'good»  'unknown»   'good |  | 63 |  | '2024-09-17»'5472-03»   'lib938»'5472-03\_lib938»'M. tuberculosis»   'unknown»   'Clade 1»   'good»  '4.8»   'mainly T»  'good»  '4.8»   'mainly T»  'good»  'unknown»   'good |  | 63 |  | '2024-09-18»'5472-03»   'lib938»'5472-03\_lib938»'M. tuberculosis»   'unknown»   'Clade 1»   'good»  '4.8»   'mainly T»  'good»  '4.8»   'mainly T»  'good»  'unknown»   'good |
| 64 |  | '2024-09-13»'5685-04»   'lib945»'5685-04\_lib945»'M. tuberculosis»   'unknown»   'Clade 1»   'good»  '4.8»   'mainly T»  'good»  '4.8»   'mainly T»  'good»  'unknown»   'good |  | 64 |  | '2024-09-17»'5685-04»   'lib945»'5685-04\_lib945»'M. tuberculosis»   'unknown»   'Clade 1»   'good»  '4.8»   'mainly T»  'good»  '4.8»   'mainly T»  'good»  'unknown»   'good |  | 64 |  | '2024-09-18»'5685-04»   'lib945»'5685-04\_lib945»'M. tuberculosis»   'unknown»   'Clade 1»   'good»  '4.8»   'mainly T»  'good»  '4.8»   'mainly T»  'good»  'unknown»   'good |
| 65 |  | '2024-09-13»'5687-04»   'lib946»'5687-04\_lib946»'M. tuberculosis»   'unknown»   'Clade 1»   'good»  '4.8»   'mainly T»  'good»  '4.8»   'mainly T»  'good»  'unknown»   'good |  | 65 |  | '2024-09-17»'5687-04»   'lib946»'5687-04\_lib946»'M. tuberculosis»   'unknown»   'Clade 1»   'good»  '4.8»   'mainly T»  'good»  '4.8»   'mainly T»  'good»  'unknown»   'good |  | 65 |  | '2024-09-18»'5687-04»   'lib946»'5687-04\_lib946»'M. tuberculosis»   'unknown»   'Clade 1»   'good»  '4.8»   'mainly T»  'good»  '4.8»   'mainly T»  'good»  'unknown»   'good |
| 66 |  | '2024-09-13»'5870-03»   'lib966»'5870-03\_lib966»'M. africanum»  '6» 'West African 2»'bad»   '6» 'West-Africa 2» 'bad»   '6» 'West-Africa 2» 'bad»   'unknown»   'bad |  | 66 |  | '2024-09-17»'5870-03»   'lib966»'5870-03\_lib966»'M. africanum»  '6» 'West African 2»'bad»   '6» 'West-Africa 2» 'bad»   '6» 'West-Africa 2» 'bad»   'unknown»   'bad |  | 66 |  | '2024-09-18»'5870-03»   'lib966»'5870-03\_lib966»'M. africanum»  '6» 'West African 2»'bad»   '6» 'West-Africa 2» 'bad»   '6» 'West-Africa 2» 'bad»   'unknown»   'bad |
| 67 |  | '2024-09-13»'5872-03»   'lib900»'5872-03\_lib900»'M. tuberculosis»   'unknown»   'Ghana» 'good»  '4.1»   'Euro-American» 'bad»   '4.1»   'Euro-American» 'bad»   'unknown»   'good |  | 67 |  | '2024-09-17»'5872-03»   'lib900»'5872-03\_lib900»'M. tuberculosis»   'unknown»   'Ghana» 'good»  '4.1»   'Euro-American» 'bad»   '4.1»   'Euro-American» 'bad»   'unknown»   'good |  | 67 |  | '2024-09-18»'5872-03»   'lib900»'5872-03\_lib900»'M. tuberculosis»   'unknown»   'Ghana» 'good»  '4.1»   'Euro-American» 'bad»   '4.1»   'Euro-American» 'bad»   'unknown»   'good |
| 68 |  | '2024-09-13»'6429-03»   'lib923»'6429-03\_lib923»'M. tuberculosis»   'unknown»   'Clade 1»   'good»  '4.1»   'Euro-American» 'good»  '4.1»   'Euro-American» 'good»  'unknown»   'good |  | 68 |  | '2024-09-17»'6429-03»   'lib923»'6429-03\_lib923»'M. tuberculosis»   'unknown»   'Clade 1»   'good»  '4.1»   'Euro-American» 'good»  '4.1»   'Euro-American» 'good»  'unknown»   'good |  | 68 |  | '2024-09-18»'6429-03»   'lib923»'6429-03\_lib923»'M. tuberculosis»   'unknown»   'Clade 1»   'good»  '4.1»   'Euro-American» 'good»  '4.1»   'Euro-American» 'good»  'unknown»   'good |
| 69 |  | '2024-09-13»'6435-03»   'lib955»'6435-03\_lib955»'M. africanum»  '6» 'West African 2»'good»  '6» 'West-Africa 2» 'good»  '6» 'West-Africa 2» 'good»  'unknown»   'good |  | 69 |  | '2024-09-17»'6435-03»   'lib955»'6435-03\_lib955»'M. africanum»  '6» 'West African 2»'good»  '6» 'West-Africa 2» 'good»  '6» 'West-Africa 2» 'good»  'unknown»   'good |  | 69 |  | '2024-09-18»'6435-03»   'lib955»'6435-03\_lib955»'M. africanum»  '6» 'West African 2»'good»  '6» 'West-Africa 2» 'good»  '6» 'West-Africa 2» 'good»  'unknown»   'good |
| 70 |  | '2024-09-13»'6463-04»   'lib910»'6463-04\_lib910»'M. tuberculosis»   'unknown»   'Clade 1»   'good»  '4.8»   'mainly T»  'good»  '4.8»   'mainly T»  'good»  'unknown»   'good |  | 70 |  | '2024-09-17»'6463-04»   'lib910»'6463-04\_lib910»'M. tuberculosis»   'unknown»   'Clade 1»   'good»  '4.8»   'mainly T»  'good»  '4.8»   'mainly T»  'good»  'unknown»   'good |  | 70 |  | '2024-09-18»'6463-04»   'lib910»'6463-04\_lib910»'M. tuberculosis»   'unknown»   'Clade 1»   'good»  '4.8»   'mainly T»  'good»  '4.8»   'mainly T»  'good»  'unknown»   'good |
| 71 |  | '2024-09-13»'6467-04»   'lib1450»   '6467-04\_lib1450»   'M. tuberculosis»   '4.1.2.1»   'Haarlem»   'good»  '4.1.2.1»   'Haarlem»   'good»  '4.1.2.1»   'Haarlem»   'good»  'unknown»   'good |  | 71 |  | '2024-09-17»'6467-04»   'lib1450»   '6467-04\_lib1450»   'M. tuberculosis»   '4.1.2.1»   'Haarlem»   'good»  '4.1.2.1»   'Haarlem»   'good»  '4.1.2.1»   'Haarlem»   'good»  'unknown»   'good |  | 71 |  | '2024-09-18»'6467-04»   'lib1450»   '6467-04\_lib1450»   'M. tuberculosis»   '4.1.2.1»   'Haarlem»   'good»  '4.1.2.1»   'Haarlem»   'good»  '4.1.2.1»   'Haarlem»   'good»  'unknown»   'good |
| 72 |  | '2024-09-13»'6637-04»   'lib899»'6637-04\_lib899»'M. tuberculosis»   '1» 'EAI»   'good»  '1.1.1» 'EAI»   'bad»   '1.1.1» 'EAI»   'bad»   'unknown»   'good |  | 72 |  | '2024-09-17»'6637-04»   'lib899»'6637-04\_lib899»'M. tuberculosis»   '1» 'EAI»   'good»  '1.1.1» 'EAI»   'bad»   '1.1.1» 'EAI»   'bad»   'unknown»   'good |  | 72 |  | '2024-09-18»'6637-04»   'lib899»'6637-04\_lib899»'M. tuberculosis»   '1» 'EAI»   'good»  '1.1.1» 'EAI»   'bad»   '1.1.1» 'EAI»   'bad»   'unknown»   'good |
| 73 |  | '2024-09-13»'6639-04»   'lib965»'6639-04\_lib965»'M. tuberculosis»   '4.3»   'LAM»   'good»  '4.3.4.2»   'LAM»   'good»  '4.3.4.2»   'LAM»   'good»  'unknown»   'good |  | 73 |  | '2024-09-17»'6639-04»   'lib965»'6639-04\_lib965»'M. tuberculosis»   '4.3»   'LAM»   'good»  '4.3.4.2»   'LAM»   'good»  '4.3.4.2»   'LAM»   'good»  'unknown»   'good |  | 73 |  | '2024-09-18»'6639-04»   'lib965»'6639-04\_lib965»'M. tuberculosis»   '4.3»   'LAM»   'good»  '4.3.4.2»   'LAM»   'good»  '4.3.4.2»   'LAM»   'good»  'unknown»   'good |
| 74 |  | '2024-09-13»'6640-04»   'lib929»'6640-04\_lib929»'M. tuberculosis»   '4.1.2.1»   'Haarlem»   'good»  '4.1.2.1»   'Haarlem»   'good»  '4.1.2.1»   'Haarlem»   'good»  'unknown»   'good |  | 74 |  | '2024-09-17»'6640-04»   'lib929»'6640-04\_lib929»'M. tuberculosis»   '4.1.2.1»   'Haarlem»   'good»  '4.1.2.1»   'Haarlem»   'good»  '4.1.2.1»   'Haarlem»   'good»  'unknown»   'good |  | 74 |  | '2024-09-18»'6640-04»   'lib929»'6640-04\_lib929»'M. tuberculosis»   '4.1.2.1»   'Haarlem»   'good»  '4.1.2.1»   'Haarlem»   'good»  '4.1.2.1»   'Haarlem»   'good»  'unknown»   'good |
| 75 |  | '2024-09-13»'6769-04»   'lib962»'6769-04\_lib962»'M. africanum»  '6» 'West African 2»'good»  '6» 'West-Africa 2» 'good»  '6» 'West-Africa 2» 'good»  'unknown»   'good |  | 75 |  | '2024-09-17»'6769-04»   'lib962»'6769-04\_lib962»'M. africanum»  '6» 'West African 2»'good»  '6» 'West-Africa 2» 'good»  '6» 'West-Africa 2» 'good»  'unknown»   'good |  | 75 |  | '2024-09-18»'6769-04»   'lib962»'6769-04\_lib962»'M. africanum»  '6» 'West African 2»'good»  '6» 'West-Africa 2» 'good»  '6» 'West-Africa 2» 'good»  'unknown»   'good |
| 76 |  | '2024-09-13»'6771-04»   'lib1462»   '6771-04\_lib1462»   'M. tuberculosis»   '4.4.1.1»   'S-type»'good»  '4.4.1.1»   'S-type»'good»  '4.4.1.1»   'S-type»'good»  'unknown»   'good |  | 76 |  | '2024-09-17»'6771-04»   'lib1462»   '6771-04\_lib1462»   'M. tuberculosis»   '4.4.1.1»   'S-type»'good»  '4.4.1.1»   'S-type»'good»  '4.4.1.1»   'S-type»'good»  'unknown»   'good |  | 76 |  | '2024-09-18»'6771-04»   'lib1462»   '6771-04\_lib1462»   'M. tuberculosis»   '4.4.1.1»   'S-type»'good»  '4.4.1.1»   'S-type»'good»  '4.4.1.1»   'S-type»'good»  'unknown»   'good |
| 77 |  | '2024-09-13»'6775-04»   'lib963»'6775-04\_lib963»'M. africanum»  '6» 'West African 2»'good»  '6» 'West-Africa 2» 'good»  '6» 'West-Africa 2» 'good»  'unknown»   'good |  | 77 |  | '2024-09-17»'6775-04»   'lib963»'6775-04\_lib963»'M. africanum»  '6» 'West African 2»'good»  '6» 'West-Africa 2» 'good»  '6» 'West-Africa 2» 'good»  'unknown»   'good |  | 77 |  | '2024-09-18»'6775-04»   'lib963»'6775-04\_lib963»'M. africanum»  '6» 'West African 2»'good»  '6» 'West-Africa 2» 'good»  '6» 'West-Africa 2» 'good»  'unknown»   'good |
| 78 |  | '2024-09-13»'6892-04»   'lib964»'6892-04\_lib964»'M. africanum»  '6» 'West African 2»'good»  '6» 'West-Africa 2» 'good»  '6» 'West-Africa 2» 'good»  'unknown»   'good |  | 78 |  | '2024-09-17»'6892-04»   'lib964»'6892-04\_lib964»'M. africanum»  '6» 'West African 2»'good»  '6» 'West-Africa 2» 'good»  '6» 'West-Africa 2» 'good»  'unknown»   'good |  | 78 |  | '2024-09-18»'6892-04»   'lib964»'6892-04\_lib964»'M. africanum»  '6» 'West African 2»'good»  '6» 'West-Africa 2» 'good»  '6» 'West-Africa 2» 'good»  'unknown»   'good |
| 79 |  | '2024-09-13»'6895-04»   'lib1459»   '6895-04\_lib1459»   'M. tuberculosis»   '1» 'EAI»   'good»  '1.1.1» 'EAI»   'bad»   '1.1.1» 'EAI»   'bad»   'unknown»   'good |  | 79 |  | '2024-09-17»'6895-04»   'lib1459»   '6895-04\_lib1459»   'M. tuberculosis»   '1» 'EAI»   'good»  '1.1.1» 'EAI»   'bad»   '1.1.1» 'EAI»   'bad»   'unknown»   'good |  | 79 |  | '2024-09-18»'6895-04»   'lib1459»   '6895-04\_lib1459»   'M. tuberculosis»   '1» 'EAI»   'good»  '1.1.1» 'EAI»   'bad»   '1.1.1» 'EAI»   'bad»   'unknown»   'good |
| 80 |  | '2024-09-13»'6897-04»   'lib954»'6897-04\_lib954»'M. africanum»  '5» 'West African 1a»   'good»  '5» 'West-Africa 1» 'bad»   '5» 'West-Africa 1» 'bad»   'unknown»   'good |  | 80 |  | '2024-09-17»'6897-04»   'lib954»'6897-04\_lib954»'M. africanum»  '5» 'West African 1a»   'good»  '5» 'West-Africa 1» 'bad»   '5» 'West-Africa 1» 'bad»   'unknown»   'good |  | 80 |  | '2024-09-18»'6897-04»   'lib954»'6897-04\_lib954»'M. africanum»  '5» 'West African 1a»   'good»  '5» 'West-Africa 1» 'bad»   '5» 'West-Africa 1» 'bad»   'unknown»   'good |
| 81 |  | '2024-09-13»'7000-03»   'lib913»'7000-03\_lib913»'M. tuberculosis»   '4.3»   'LAM»   'good»  '4.3.4.2»   'LAM»   'good»  '4.3.4.2»   'LAM»   'good»  'unknown»   'good |  | 81 |  | '2024-09-17»'7000-03»   'lib913»'7000-03\_lib913»'M. tuberculosis»   '4.3»   'LAM»   'good»  '4.3.4.2»   'LAM»   'good»  '4.3.4.2»   'LAM»   'good»  'unknown»   'good |  | 81 |  | '2024-09-18»'7000-03»   'lib913»'7000-03\_lib913»'M. tuberculosis»   '4.3»   'LAM»   'good»  '4.3.4.2»   'LAM»   'good»  '4.3.4.2»   'LAM»   'good»  'unknown»   'good |
| 82 |  | '2024-09-13»'7135-04»   'lib987»'7135-04\_lib987»'M. tuberculosis»   '2» 'Beijing»   'good»  '2.2.1» 'Beijing»   'good»  '2.2.1» 'Beijing»   'good»  'Ancestral 3»   'good |  | 82 |  | '2024-09-17»'7135-04»   'lib987»'7135-04\_lib987»'M. tuberculosis»   '2» 'Beijing»   'good»  '2.2.1» 'Beijing»   'good»  '2.2.1» 'Beijing»   'good»  'Ancestral 3»   'good |  | 82 |  | '2024-09-18»'7135-04»   'lib987»'7135-04\_lib987»'M. tuberculosis»   '2» 'Beijing»   'good»  '2.2.1» 'Beijing»   'good»  '2.2.1» 'Beijing»   'good»  'Ancestral 3»   'good |
| 83 |  | '2024-09-13»'7514-04»   'lib1460»   '7514-04\_lib1460»   'M. tuberculosis»   '4.3»   'LAM»   'good»  '4.3.4.1»   'LAM»   'good»  '4.3.4.1»   'LAM»   'good»  'unknown»   'good |  | 83 |  | '2024-09-17»'7514-04»   'lib1460»   '7514-04\_lib1460»   'M. tuberculosis»   '4.3»   'LAM»   'good»  '4.3.4.1»   'LAM»   'good»  '4.3.4.1»   'LAM»   'good»  'unknown»   'good |  | 83 |  | '2024-09-18»'7514-04»   'lib1460»   '7514-04\_lib1460»   'M. tuberculosis»   '4.3»   'LAM»   'good»  '4.3.4.1»   'LAM»   'good»  '4.3.4.1»   'LAM»   'good»  'unknown»   'good |
| 84 |  | '2024-09-13»'7516-04»   'lib920»'7516-04\_lib920»'M. tuberculosis»   '4.3»   'LAM»   'good»  '4.3.4.1»   'LAM»   'good»  '4.3.4.1»   'LAM»   'good»  'unknown»   'good |  | 84 |  | '2024-09-17»'7516-04»   'lib920»'7516-04\_lib920»'M. tuberculosis»   '4.3»   'LAM»   'good»  '4.3.4.1»   'LAM»   'good»  '4.3.4.1»   'LAM»   'good»  'unknown»   'good |  | 84 |  | '2024-09-18»'7516-04»   'lib920»'7516-04\_lib920»'M. tuberculosis»   '4.3»   'LAM»   'good»  '4.3.4.1»   'LAM»   'good»  '4.3.4.1»   'LAM»   'good»  'unknown»   'good |
| 85 |  | '2024-09-13»'7517-04»   'lib930»'7517-04\_lib930»'M. tuberculosis»   'unknown»   'Clade 1»   'good»  '4.1.1.3»   'X-type»'good»  '4.1.1.3»   'X-type»'good»  'unknown»   'good |  | 85 |  | '2024-09-17»'7517-04»   'lib930»'7517-04\_lib930»'M. tuberculosis»   'unknown»   'Clade 1»   'good»  '4.1.1.3»   'X-type»'good»  '4.1.1.3»   'X-type»'good»  'unknown»   'good |  | 85 |  | '2024-09-18»'7517-04»   'lib930»'7517-04\_lib930»'M. tuberculosis»   'unknown»   'Clade 1»   'good»  '4.1.1.3»   'X-type»'good»  '4.1.1.3»   'X-type»'good»  'unknown»   'good |
| 86 |  | '2024-09-13»'7520-04»   'lib1461»   '7520-04\_lib1461»   'M. tuberculosis»   '4.3»   'LAM»   'good»  '4.3.3» 'LAM»   'good»  '4.3.3» 'LAM»   'good»  'unknown»   'good |  | 86 |  | '2024-09-17»'7520-04»   'lib1461»   '7520-04\_lib1461»   'M. tuberculosis»   '4.3»   'LAM»   'good»  '4.3.3» 'LAM»   'good»  '4.3.3» 'LAM»   'good»  'unknown»   'good |  | 86 |  | '2024-09-18»'7520-04»   'lib1461»   '7520-04\_lib1461»   'M. tuberculosis»   '4.3»   'LAM»   'good»  '4.3.3» 'LAM»   'good»  '4.3.3» 'LAM»   'good»  'unknown»   'good |
| 87 |  | '2024-09-13»'7538-03»   'lib948»'7538-03\_lib948»'M. tuberculosis»   'unknown»   'Clade 1»   'good»  '4.1.1.3»   'X-type»'good»  '4.1.1.3»   'X-type»'good»  'unknown»   'good |  | 87 |  | '2024-09-17»'7538-03»   'lib948»'7538-03\_lib948»'M. tuberculosis»   'unknown»   'Clade 1»   'good»  '4.1.1.3»   'X-type»'good»  '4.1.1.3»   'X-type»'good»  'unknown»   'good |  | 87 |  | '2024-09-18»'7538-03»   'lib948»'7538-03\_lib948»'M. tuberculosis»   'unknown»   'Clade 1»   'good»  '4.1.1.3»   'X-type»'good»  '4.1.1.3»   'X-type»'good»  'unknown»   'good |
| 88 |  | '2024-09-13»'8082-03»   'lib932»'8082-03\_lib932»'M. tuberculosis»   'unknown»   'Clade 1»   'good»  '4.1»   'Euro-American» 'good»  '4.1»   'Euro-American» 'good»  'unknown»   'good |  | 88 |  | '2024-09-17»'8082-03»   'lib932»'8082-03\_lib932»'M. tuberculosis»   'unknown»   'Clade 1»   'good»  '4.1»   'Euro-American» 'good»  '4.1»   'Euro-American» 'good»  'unknown»   'good |  | 88 |  | '2024-09-18»'8082-03»   'lib932»'8082-03\_lib932»'M. tuberculosis»   'unknown»   'Clade 1»   'good»  '4.1»   'Euro-American» 'good»  '4.1»   'Euro-American» 'good»  'unknown»   'good |
| 89 |  | '2024-09-13»'8864-03»   'lib967»'8864-03\_lib967»'M. africanum»  '6» 'West African 2»'good»  '6» 'West-Africa 2» 'good»  '6» 'West-Africa 2» 'good»  'unknown»   'good |  | 89 |  | '2024-09-17»'8864-03»   'lib967»'8864-03\_lib967»'M. africanum»  '6» 'West African 2»'good»  '6» 'West-Africa 2» 'good»  '6» 'West-Africa 2» 'good»  'unknown»   'good |  | 89 |  | '2024-09-18»'8864-03»   'lib967»'8864-03\_lib967»'M. africanum»  '6» 'West African 2»'good»  '6» 'West-Africa 2» 'good»  '6» 'West-Africa 2» 'good»  'unknown»   'good |
| 90 |  | '2024-09-13»'8867-03»   'lib968»'8867-03\_lib968»'M. africanum»  '6» 'West African 2»'good»  '6» 'West-Africa 2» 'good»  '6» 'West-Africa 2» 'good»  'unknown»   'good |  | 90 |  | '2024-09-17»'8867-03»   'lib968»'8867-03\_lib968»'M. africanum»  '6» 'West African 2»'good»  '6» 'West-Africa 2» 'good»  '6» 'West-Africa 2» 'good»  'unknown»   'good |  | 90 |  | '2024-09-18»'8867-03»   'lib968»'8867-03\_lib968»'M. africanum»  '6» 'West African 2»'good»  '6» 'West-Africa 2» 'good»  '6» 'West-Africa 2» 'good»  'unknown»   'good |
| 91 |  | '2024-09-13»'8868-03»   'lib950»'8868-03\_lib950»'M. africanum»  '5» 'West African 1a»   'good»  '5» 'West-Africa 1» 'bad»   '5» 'West-Africa 1» 'bad»   'unknown»   'good |  | 91 |  | '2024-09-17»'8868-03»   'lib950»'8868-03\_lib950»'M. africanum»  '5» 'West African 1a»   'good»  '5» 'West-Africa 1» 'bad»   '5» 'West-Africa 1» 'bad»   'unknown»   'good |  | 91 |  | '2024-09-18»'8868-03»   'lib950»'8868-03\_lib950»'M. africanum»  '5» 'West African 1a»   'good»  '5» 'West-Africa 1» 'bad»   '5» 'West-Africa 1» 'bad»   'unknown»   'good |

Araxis Merge (but not the data content of this report) is Copyright © 1993–2024 Araxis Ltd (www.araxis.com). All rights reserved.
