## Supplementary material for "MTBseq-nf: Enabling Scalable Tuberculosis Genomics “Big Data” Analysis through a User-Friendly Nextflow Wrapper for MTBseq pipeline": SD-6 Intra-modal analysis, with 3-way HTML diff reports generated by Araxis merge software: SD-6-12-intra-modal-araxiscompare-pub-90samples-mtbseq-nf-parallel-runs-statistics.html

xml version="1.0" encoding="utf-8"?


Araxis Merge File Comparison Report 

### **Araxis Merge File Comparison Report**

Produced by **Araxis Merge** on **2024/12/18, 16:55 GMT+02:00**. See www.araxis.com for information about Merge. This report uses XHTML and CSS2, and is best viewed with a modern standards-compliant browser. For optimum results when printing this report, use landscape orientation and enable printing of background images and colours in your browser.

##### **1. Files compared**

| # | Location | File | Last Modified |
| --- | --- | --- | --- |
| 1 | /Users/abhi/projects/MTBseq-nf/\_resources/publication/manuscript-and-analysis/v2/pub-90samples-mtbseq-nf-parallel-run1/tbstats/Statistics | Mapping\_and\_Variant\_Statistics.tab | 2024/09/13, 18:59 GMT+02:00 |
| 2 | /Users/abhi/projects/MTBseq-nf/\_resources/publication/manuscript-and-analysis/v2/pub-90samples-mtbseq-nf-parallel-run2/tbstats/Statistics | Mapping\_and\_Variant\_Statistics.tab | 2024/09/17, 22:12 GMT+02:00 |
| 3 | /Users/abhi/projects/MTBseq-nf/\_resources/publication/manuscript-and-analysis/v2/pub-90samples-mtbseq-nf-parallel-run3/tbstats/Statistics | Mapping\_and\_Variant\_Statistics.tab | 2024/09/18, 18:49 GMT+02:00 |
| **Note:** Merge considers the second file to be the common ancestor of the others. | | | |

##### **2. Comparison summary**

| Description | Between Files 1 and 2 | | Between Files 2 and 3 | | Relative to Common Ancestor | |
| --- | --- | --- | --- | --- | --- | --- |
|  | Text Blocks | Lines | Text Blocks | Lines | Text Blocks | Lines |
| Unchanged | 1 | 2 | 1 | 2 |  |  |
| Changed | 1 | 180 | 1 | 180 | 0 | 0 |
| Inserted | 0 | 0 | 0 | 0 | 0 | 0 |
| Removed | 0 | 0 | 0 | 0 | 0 | 0 |
| **Note:** An automatic merge would leave 1 conflict(s). | | | | | | |

##### **3. Comparison options**

|  |  |
| --- | --- |
| Whitespace | Consecutive whitespace is treated as a single space |
| Character case | Differences in character case are significant |
| Line endings | Differences in line endings (`CR` and `LF` characters) are ignored |
| CR/LF characters | Not shown in the comparison detail |
| Active line-pairing rules |  |

##### **4. Active regular expressions**

No regular expressions were active.

##### **5. Comparison detail**

|  |  |  |  |  |  |  |  |  |  |  |
| --- | --- | --- | --- | --- | --- | --- | --- | --- | --- | --- |
| 1 |  | Date»   SampleID»   LibraryID»  FullID» Total Reads»Mapped Reads»   % Mapped Reads» Genome Size»Genome GC»  (Any) Total Bases»  % (Any) Total Bases»(Any) GC-Content»   (Any) Coverage mean»(Any) Coverage median»  (Unambiguous) Total Bases»  % (Unambiguous) Total Bases»(Unambiguous) GC-Content»   (Unambiguous) Coverage mean»(Unambiguous) Coverage median»  SNPs»   Deletions»  Insertions» Uncovered»  Substitutions (Including Stop Codons) |  | 1 |  | Date»   SampleID»   LibraryID»  FullID» Total Reads»Mapped Reads»   % Mapped Reads» Genome Size»Genome GC»  (Any) Total Bases»  % (Any) Total Bases»(Any) GC-Content»   (Any) Coverage mean»(Any) Coverage median»  (Unambiguous) Total Bases»  % (Unambiguous) Total Bases»(Unambiguous) GC-Content»   (Unambiguous) Coverage mean»(Unambiguous) Coverage median»  SNPs»   Deletions»  Insertions» Uncovered»  Substitutions (Including Stop Codons) |  | 1 |  | Date»   SampleID»   LibraryID»  FullID» Total Reads»Mapped Reads»   % Mapped Reads» Genome Size»Genome GC»  (Any) Total Bases»  % (Any) Total Bases»(Any) GC-Content»   (Any) Coverage mean»(Any) Coverage median»  (Unambiguous) Total Bases»  % (Unambiguous) Total Bases»(Unambiguous) GC-Content»   (Unambiguous) Coverage mean»(Unambiguous) Coverage median»  SNPs»   Deletions»  Insertions» Uncovered»  Substitutions (Including Stop Codons) |
| 2 |  | '2024-09-13»'10010-03»  'lib951»'10010-03\_lib951»   '1359180»   '1345417»   '98.99» '4411532»   '65.61» '4370731»   '0.99»  '65.54» '68.05» '69»'4275440»   '0.97»  '65.33» '69.32» '70»'2100»  '663»   '226»   '40801» '1127 |  | 2 |  | '2024-09-17»'10010-03»  'lib951»'10010-03\_lib951»   '1359180»   '1345417»   '98.99» '4411532»   '65.61» '4370731»   '0.99»  '65.54» '68.05» '69»'4275440»   '0.97»  '65.33» '69.32» '70»'2100»  '663»   '226»   '40801» '1127 |  | 2 |  | '2024-09-18»'10010-03»  'lib951»'10010-03\_lib951»   '1359180»   '1345417»   '98.99» '4411532»   '65.61» '4370731»   '0.99»  '65.54» '68.05» '69»'4275440»   '0.97»  '65.33» '69.32» '70»'2100»  '663»   '226»   '40801» '1127 |
| 3 |  | '2024-09-13»'10011-03»  'lib914»'10011-03\_lib914»   '546250»'543451»'99.49» '4411532»   '65.61» '4360648»   '0.99»  '65.53» '28.64» '29»'4174446»   '0.95»  '65.21» '29.57» '29»'871»   '468»   '139»   '50884» '472 |  | 3 |  | '2024-09-17»'10011-03»  'lib914»'10011-03\_lib914»   '546250»'543451»'99.49» '4411532»   '65.61» '4360648»   '0.99»  '65.53» '28.64» '29»'4174446»   '0.95»  '65.21» '29.57» '29»'871»   '468»   '139»   '50884» '472 |  | 3 |  | '2024-09-18»'10011-03»  'lib914»'10011-03\_lib914»   '546250»'543451»'99.49» '4411532»   '65.61» '4360648»   '0.99»  '65.53» '28.64» '29»'4174446»   '0.95»  '65.21» '29.57» '29»'871»   '468»   '139»   '50884» '472 |
| 4 |  | '2024-09-13»'10012-03»  'lib970»'10012-03\_lib970»   '2552722»   '2530420»   '99.13» '4411532»   '65.61» '4394284»   '1.00»  '65.58» '126.75»'130»   '4339304»   '0.98»  '65.43» '128.22»'130»   '980»   '331»   '113»   '17248» '482 |  | 4 |  | '2024-09-17»'10012-03»  'lib970»'10012-03\_lib970»   '2552722»   '2530420»   '99.13» '4411532»   '65.61» '4394284»   '1.00»  '65.58» '126.75»'130»   '4339304»   '0.98»  '65.43» '128.22»'130»   '980»   '331»   '113»   '17248» '482 |  | 4 |  | '2024-09-18»'10012-03»  'lib970»'10012-03\_lib970»   '2552722»   '2530420»   '99.13» '4411532»   '65.61» '4394284»   '1.00»  '65.58» '126.75»'130»   '4339304»   '0.98»  '65.43» '128.22»'130»   '980»   '331»   '113»   '17248» '482 |
| 5 |  | '2024-09-13»'10205-03»  'lib915»'10205-03\_lib915»   '292577»'286579»'97.95» '4411532»   '65.61» '4338092»   '0.98»  '65.48» '14.40» '14»'3311058»   '0.75»  '65.21» '16.26» '16»'607»   '98»'62»'73440» '312 |  | 5 |  | '2024-09-17»'10205-03»  'lib915»'10205-03\_lib915»   '292577»'286579»'97.95» '4411532»   '65.61» '4338092»   '0.98»  '65.48» '14.40» '14»'3311058»   '0.75»  '65.21» '16.26» '16»'607»   '98»'62»'73440» '312 |  | 5 |  | '2024-09-18»'10205-03»  'lib915»'10205-03\_lib915»   '292577»'286579»'97.95» '4411532»   '65.61» '4338092»   '0.98»  '65.48» '14.40» '14»'3311058»   '0.75»  '65.21» '16.26» '16»'607»   '98»'62»'73440» '312 |
| 6 |  | '2024-09-13»'10206-03»  'lib916»'10206-03\_lib916»   '532964»'529320»'99.32» '4411532»   '65.61» '4361069»   '0.99»  '65.52» '27.98» '28»'4172738»   '0.95»  '65.20» '28.90» '28»'852»   '364»   '144»   '50463» '465 |  | 6 |  | '2024-09-17»'10206-03»  'lib916»'10206-03\_lib916»   '532964»'529320»'99.32» '4411532»   '65.61» '4361069»   '0.99»  '65.52» '27.98» '28»'4172738»   '0.95»  '65.20» '28.90» '28»'852»   '364»   '144»   '50463» '465 |  | 6 |  | '2024-09-18»'10206-03»  'lib916»'10206-03\_lib916»   '532964»'529320»'99.32» '4411532»   '65.61» '4361069»   '0.99»  '65.52» '27.98» '28»'4172738»   '0.95»  '65.20» '28.90» '28»'852»   '364»   '144»   '50463» '465 |
| 7 |  | '2024-09-13»'10207-03»  'lib973»'10207-03\_lib973»   '1571919»   '1563056»   '99.44» '4411532»   '65.61» '4366060»   '0.99»  '65.57» '79.04» '80»'4296894»   '0.97»  '65.39» '80.18» '80»'816»   '402»   '118»   '45472» '408 |  | 7 |  | '2024-09-17»'10207-03»  'lib973»'10207-03\_lib973»   '1571919»   '1563056»   '99.44» '4411532»   '65.61» '4366060»   '0.99»  '65.57» '79.04» '80»'4296894»   '0.97»  '65.39» '80.18» '80»'816»   '402»   '118»   '45472» '408 |  | 7 |  | '2024-09-18»'10207-03»  'lib973»'10207-03\_lib973»   '1571919»   '1563056»   '99.44» '4411532»   '65.61» '4366060»   '0.99»  '65.57» '79.04» '80»'4296894»   '0.97»  '65.39» '80.18» '80»'816»   '402»   '118»   '45472» '408 |
| 8 |  | '2024-09-13»'10208-03»  'lib1613»   '10208-03\_lib1613»  '3592891»   '3563381»   '99.18» '4411532»   '65.61» '4359095»   '0.99»  '65.57» '209.49»'214»   '4333381»   '0.98»  '65.51» '210.68»'214»   '2395»  '605»   '351»   '52437» '1277 |  | 8 |  | '2024-09-17»'10208-03»  'lib1613»   '10208-03\_lib1613»  '3592891»   '3563381»   '99.18» '4411532»   '65.61» '4359095»   '0.99»  '65.57» '209.49»'214»   '4333381»   '0.98»  '65.51» '210.68»'214»   '2395»  '605»   '351»   '52437» '1277 |  | 8 |  | '2024-09-18»'10208-03»  'lib1613»   '10208-03\_lib1613»  '3592891»   '3563381»   '99.18» '4411532»   '65.61» '4359095»   '0.99»  '65.57» '209.49»'214»   '4333381»   '0.98»  '65.51» '210.68»'214»   '2395»  '605»   '351»   '52437» '1277 |
| 9 |  | '2024-09-13»'10348-03»  'lib917»'10348-03\_lib917»   '509599»'507232»'99.54» '4411532»   '65.61» '4325617»   '0.98»  '65.52» '27.11» '27»'4125386»   '0.94»  '65.22» '28.02» '28»'789»   '199»   '150»   '85915» '414 |  | 9 |  | '2024-09-17»'10348-03»  'lib917»'10348-03\_lib917»   '509599»'507232»'99.54» '4411532»   '65.61» '4325617»   '0.98»  '65.52» '27.11» '27»'4125386»   '0.94»  '65.22» '28.02» '28»'789»   '199»   '150»   '85915» '414 |  | 9 |  | '2024-09-18»'10348-03»  'lib917»'10348-03\_lib917»   '509599»'507232»'99.54» '4411532»   '65.61» '4325617»   '0.98»  '65.52» '27.11» '27»'4125386»   '0.94»  '65.22» '28.02» '28»'789»   '199»   '150»   '85915» '414 |
| 10 |  | '2024-09-13»'10349-03»  'lib924»'10349-03\_lib924»   '358709»'354714»'98.89» '4411532»   '65.61» '4356735»   '0.99»  '65.48» '17.83» '18»'3821510»   '0.87»  '64.95» '19.21» '19»'795»   '159»   '78»'54797» '419 |  | 10 |  | '2024-09-17»'10349-03»  'lib924»'10349-03\_lib924»   '358709»'354714»'98.89» '4411532»   '65.61» '4356735»   '0.99»  '65.48» '17.83» '18»'3821510»   '0.87»  '64.95» '19.21» '19»'795»   '159»   '78»'54797» '419 |  | 10 |  | '2024-09-18»'10349-03»  'lib924»'10349-03\_lib924»   '358709»'354714»'98.89» '4411532»   '65.61» '4356735»   '0.99»  '65.48» '17.83» '18»'3821510»   '0.87»  '64.95» '19.21» '19»'795»   '159»   '78»'54797» '419 |
| 11 |  | '2024-09-13»'10350-03»  'lib1470»   '10350-03\_lib1470»  '1783739»   '1770487»   '99.26» '4411532»   '65.61» '4373384»   '0.99»  '65.57» '104.65»'105»   '4333962»   '0.98»  '65.47» '105.54»'106»   '1014»  '481»   '146»   '38148» '535 |  | 11 |  | '2024-09-17»'10350-03»  'lib1470»   '10350-03\_lib1470»  '1783739»   '1770487»   '99.26» '4411532»   '65.61» '4373384»   '0.99»  '65.57» '104.65»'105»   '4333962»   '0.98»  '65.47» '105.54»'106»   '1014»  '481»   '146»   '38148» '535 |  | 11 |  | '2024-09-18»'10350-03»  'lib1470»   '10350-03\_lib1470»  '1783739»   '1770487»   '99.26» '4411532»   '65.61» '4373384»   '0.99»  '65.57» '104.65»'105»   '4333962»   '0.98»  '65.47» '105.54»'106»   '1014»  '481»   '146»   '38148» '535 |
| 12 |  | '2024-09-13»'10517-03»  'lib939»'10517-03\_lib939»   '1165615»   '1156108»   '99.18» '4411532»   '65.61» '4382870»   '0.99»  '65.56» '58.00» '59»'4267099»   '0.97»  '65.32» '59.28» '59»'527»   '173»   '74»'28662» '291 |  | 12 |  | '2024-09-17»'10517-03»  'lib939»'10517-03\_lib939»   '1165615»   '1156108»   '99.18» '4411532»   '65.61» '4382870»   '0.99»  '65.56» '58.00» '59»'4267099»   '0.97»  '65.32» '59.28» '59»'527»   '173»   '74»'28662» '291 |  | 12 |  | '2024-09-18»'10517-03»  'lib939»'10517-03\_lib939»   '1165615»   '1156108»   '99.18» '4411532»   '65.61» '4382870»   '0.99»  '65.56» '58.00» '59»'4267099»   '0.97»  '65.32» '59.28» '59»'527»   '173»   '74»'28662» '291 |
| 13 |  | '2024-09-13»'11096-03»  'lib940»'11096-03\_lib940»   '1823704»   '1807550»   '99.11» '4411532»   '65.61» '4391302»   '1.00»  '65.57» '91.40» '93»'4316772»   '0.98»  '65.38» '92.80» '93»'549»   '253»   '67»'20230» '303 |  | 13 |  | '2024-09-17»'11096-03»  'lib940»'11096-03\_lib940»   '1823704»   '1807550»   '99.11» '4411532»   '65.61» '4391302»   '1.00»  '65.57» '91.40» '93»'4316772»   '0.98»  '65.38» '92.80» '93»'549»   '253»   '67»'20230» '303 |  | 13 |  | '2024-09-18»'11096-03»  'lib940»'11096-03\_lib940»   '1823704»   '1807550»   '99.11» '4411532»   '65.61» '4391302»   '1.00»  '65.57» '91.40» '93»'4316772»   '0.98»  '65.38» '92.80» '93»'549»   '253»   '67»'20230» '303 |
| 14 |  | '2024-09-13»'11097-03»  'lib933»'11097-03\_lib933»   '2927362»   '2903114»   '99.17» '4411532»   '65.61» '4393861»   '1.00»  '65.58» '142.46»'146»   '4343505»   '0.98»  '65.44» '143.98»'147»   '1010»  '279»   '122»   '17671» '502 |  | 14 |  | '2024-09-17»'11097-03»  'lib933»'11097-03\_lib933»   '2927362»   '2903114»   '99.17» '4411532»   '65.61» '4393861»   '1.00»  '65.58» '142.46»'146»   '4343505»   '0.98»  '65.44» '143.98»'147»   '1010»  '279»   '122»   '17671» '502 |  | 14 |  | '2024-09-18»'11097-03»  'lib933»'11097-03\_lib933»   '2927362»   '2903114»   '99.17» '4411532»   '65.61» '4393861»   '1.00»  '65.58» '142.46»'146»   '4343505»   '0.98»  '65.44» '143.98»'147»   '1010»  '279»   '122»   '17671» '502 |
| 15 |  | '2024-09-13»'11818-03»  'lib902»'11818-03\_lib902»   '591264»'585978»'99.11» '4411532»   '65.61» '4360907»   '0.99»  '65.51» '29.47» '29»'4123713»   '0.93»  '65.16» '30.64» '30»'867»   '203»   '156»   '50625» '435 |  | 15 |  | '2024-09-17»'11818-03»  'lib902»'11818-03\_lib902»   '591264»'585978»'99.11» '4411532»   '65.61» '4360907»   '0.99»  '65.51» '29.47» '29»'4123713»   '0.93»  '65.16» '30.64» '30»'867»   '203»   '156»   '50625» '435 |  | 15 |  | '2024-09-18»'11818-03»  'lib902»'11818-03\_lib902»   '591264»'585978»'99.11» '4411532»   '65.61» '4360907»   '0.99»  '65.51» '29.47» '29»'4123713»   '0.93»  '65.16» '30.64» '30»'867»   '203»   '156»   '50625» '435 |
| 16 |  | '2024-09-13»'11821-03»  'lib952»'11821-03\_lib952»   '2205544»   '2183658»   '99.01» '4411532»   '65.61» '4380116»   '0.99»  '65.56» '110.90»'113»   '4318666»   '0.98»  '65.39» '112.32»'114»   '2209»  '663»   '344»   '31416» '1193 |  | 16 |  | '2024-09-17»'11821-03»  'lib952»'11821-03\_lib952»   '2205544»   '2183658»   '99.01» '4411532»   '65.61» '4380116»   '0.99»  '65.56» '110.90»'113»   '4318666»   '0.98»  '65.39» '112.32»'114»   '2209»  '663»   '344»   '31416» '1193 |  | 16 |  | '2024-09-18»'11821-03»  'lib952»'11821-03\_lib952»   '2205544»   '2183658»   '99.01» '4411532»   '65.61» '4380116»   '0.99»  '65.56» '110.90»'113»   '4318666»   '0.98»  '65.39» '112.32»'114»   '2209»  '663»   '344»   '31416» '1193 |
| 17 |  | '2024-09-13»'11822-03»  'lib903»'11822-03\_lib903»   '1785165»   '1767550»   '99.01» '4411532»   '65.61» '4380455»   '0.99»  '65.59» '89.10» '91»'4301366»   '0.98»  '65.39» '90.55» '91»'948»   '244»   '153»   '31077» '492 |  | 17 |  | '2024-09-17»'11822-03»  'lib903»'11822-03\_lib903»   '1785165»   '1767550»   '99.01» '4411532»   '65.61» '4380455»   '0.99»  '65.59» '89.10» '91»'4301366»   '0.98»  '65.39» '90.55» '91»'948»   '244»   '153»   '31077» '492 |  | 17 |  | '2024-09-18»'11822-03»  'lib903»'11822-03\_lib903»   '1785165»   '1767550»   '99.01» '4411532»   '65.61» '4380455»   '0.99»  '65.59» '89.10» '91»'4301366»   '0.98»  '65.39» '90.55» '91»'948»   '244»   '153»   '31077» '492 |
| 18 |  | '2024-09-13»'12655-03»  'lib1516»   '12655-03\_lib1516»  '2294395»   '2284164»   '99.55» '4411532»   '65.61» '4366750»   '0.99»  '65.58» '118.89»'121»   '4329010»   '0.98»  '65.49» '119.86»'121»   '860»   '548»   '107»   '44782» '426 |  | 18 |  | '2024-09-17»'12655-03»  'lib1516»   '12655-03\_lib1516»  '2294395»   '2284164»   '99.55» '4411532»   '65.61» '4366750»   '0.99»  '65.58» '118.89»'121»   '4329010»   '0.98»  '65.49» '119.86»'121»   '860»   '548»   '107»   '44782» '426 |  | 18 |  | '2024-09-18»'12655-03»  'lib1516»   '12655-03\_lib1516»  '2294395»   '2284164»   '99.55» '4411532»   '65.61» '4366750»   '0.99»  '65.58» '118.89»'121»   '4329010»   '0.98»  '65.49» '119.86»'121»   '860»   '548»   '107»   '44782» '426 |
| 19 |  | '2024-09-13»'12657-03»  'lib934»'12657-03\_lib934»   '1947169»   '1929629»   '99.10» '4411532»   '65.61» '4391636»   '1.00»  '65.59» '94.68» '97»'4325412»   '0.98»  '65.41» '95.97» '97»'985»   '240»   '145»   '19896» '509 |  | 19 |  | '2024-09-17»'12657-03»  'lib934»'12657-03\_lib934»   '1947169»   '1929629»   '99.10» '4411532»   '65.61» '4391636»   '1.00»  '65.59» '94.68» '97»'4325412»   '0.98»  '65.41» '95.97» '97»'985»   '240»   '145»   '19896» '509 |  | 19 |  | '2024-09-18»'12657-03»  'lib934»'12657-03\_lib934»   '1947169»   '1929629»   '99.10» '4411532»   '65.61» '4391636»   '1.00»  '65.59» '94.68» '97»'4325412»   '0.98»  '65.41» '95.97» '97»'985»   '240»   '145»   '19896» '509 |
| 20 |  | '2024-09-13»'12658-03»  'lib1469»   '12658-03\_lib1469»  '1705934»   '1695826»   '99.41» '4411532»   '65.61» '4369558»   '0.99»  '65.58» '99.84» '101»   '4331248»   '0.98»  '65.49» '100.67»'101»   '1503»  '589»   '290»   '41974» '773 |  | 20 |  | '2024-09-17»'12658-03»  'lib1469»   '12658-03\_lib1469»  '1705934»   '1695826»   '99.41» '4411532»   '65.61» '4369558»   '0.99»  '65.58» '99.84» '101»   '4331248»   '0.98»  '65.49» '100.67»'101»   '1503»  '589»   '290»   '41974» '773 |  | 20 |  | '2024-09-18»'12658-03»  'lib1469»   '12658-03\_lib1469»  '1705934»   '1695826»   '99.41» '4411532»   '65.61» '4369558»   '0.99»  '65.58» '99.84» '101»   '4331248»   '0.98»  '65.49» '100.67»'101»   '1503»  '589»   '290»   '41974» '773 |
| 21 |  | '2024-09-13»'1322-04»   'lib925»'1322-04\_lib925»'1614637»   '1604845»   '99.39» '4411532»   '65.61» '4373502»   '0.99»  '65.58» '81.29» '83»'4298413»   '0.97»  '65.38» '82.55» '83»'550»   '219»   '91»'38030» '294 |  | 21 |  | '2024-09-17»'1322-04»   'lib925»'1322-04\_lib925»'1614637»   '1604845»   '99.39» '4411532»   '65.61» '4373502»   '0.99»  '65.58» '81.29» '83»'4298413»   '0.97»  '65.38» '82.55» '83»'550»   '219»   '91»'38030» '294 |  | 21 |  | '2024-09-18»'1322-04»   'lib925»'1322-04\_lib925»'1614637»   '1604845»   '99.39» '4411532»   '65.61» '4373502»   '0.99»  '65.58» '81.29» '83»'4298413»   '0.97»  '65.38» '82.55» '83»'550»   '219»   '91»'38030» '294 |
| 22 |  | '2024-09-13»'1324-04»   'lib931»'1324-04\_lib931»'2761133»   '2745427»   '99.43» '4411532»   '65.61» '4386279»   '0.99»  '65.58» '136.98»'141»   '4334533»   '0.98»  '65.44» '138.48»'141»   '828»   '133»   '87»'25253» '428 |  | 22 |  | '2024-09-17»'1324-04»   'lib931»'1324-04\_lib931»'2761133»   '2745427»   '99.43» '4411532»   '65.61» '4386279»   '0.99»  '65.58» '136.98»'141»   '4334533»   '0.98»  '65.44» '138.48»'141»   '828»   '133»   '87»'25253» '428 |  | 22 |  | '2024-09-18»'1324-04»   'lib931»'1324-04\_lib931»'2761133»   '2745427»   '99.43» '4411532»   '65.61» '4386279»   '0.99»  '65.58» '136.98»'141»   '4334533»   '0.98»  '65.44» '138.48»'141»   '828»   '133»   '87»'25253» '428 |
| 23 |  | '2024-09-13»'1327-04»   'lib957»'1327-04\_lib957»'1291239»   '1275916»   '98.81» '4411532»   '65.61» '4368641»   '0.99»  '65.54» '64.59» '66»'4261104»   '0.97»  '65.34» '66.04» '66»'2221»  '414»   '256»   '42891» '1204 |  | 23 |  | '2024-09-17»'1327-04»   'lib957»'1327-04\_lib957»'1291239»   '1275916»   '98.81» '4411532»   '65.61» '4368641»   '0.99»  '65.54» '64.59» '66»'4261104»   '0.97»  '65.34» '66.04» '66»'2221»  '414»   '256»   '42891» '1204 |  | 23 |  | '2024-09-18»'1327-04»   'lib957»'1327-04\_lib957»'1291239»   '1275916»   '98.81» '4411532»   '65.61» '4368641»   '0.99»  '65.54» '64.59» '66»'4261104»   '0.97»  '65.34» '66.04» '66»'2221»  '414»   '256»   '42891» '1204 |
| 24 |  | '2024-09-13»'1597-04»   'lib918»'1597-04\_lib918»'431877»'429798»'99.52» '4411532»   '65.61» '4343293»   '0.98»  '65.50» '22.65» '22»'4077836»   '0.92»  '65.13» '23.66» '23»'785»   '320»   '123»   '68239» '425 |  | 24 |  | '2024-09-17»'1597-04»   'lib918»'1597-04\_lib918»'431877»'429798»'99.52» '4411532»   '65.61» '4343293»   '0.98»  '65.50» '22.65» '22»'4077836»   '0.92»  '65.13» '23.66» '23»'785»   '320»   '123»   '68239» '425 |  | 24 |  | '2024-09-18»'1597-04»   'lib918»'1597-04\_lib918»'431877»'429798»'99.52» '4411532»   '65.61» '4343293»   '0.98»  '65.50» '22.65» '22»'4077836»   '0.92»  '65.13» '23.66» '23»'785»   '320»   '123»   '68239» '425 |
| 25 |  | '2024-09-13»'1599-04»   'lib974»'1599-04\_lib974»'3035794»   '3009801»   '99.14» '4411532»   '65.61» '4384788»   '0.99»  '65.60» '150.95»'154»   '4333524»   '0.98»  '65.46» '152.60»'155»   '963»   '348»   '173»   '26744» '496 |  | 25 |  | '2024-09-17»'1599-04»   'lib974»'1599-04\_lib974»'3035794»   '3009801»   '99.14» '4411532»   '65.61» '4384788»   '0.99»  '65.60» '150.95»'154»   '4333524»   '0.98»  '65.46» '152.60»'155»   '963»   '348»   '173»   '26744» '496 |  | 25 |  | '2024-09-18»'1599-04»   'lib974»'1599-04\_lib974»'3035794»   '3009801»   '99.14» '4411532»   '65.61» '4384788»   '0.99»  '65.60» '150.95»'154»   '4333524»   '0.98»  '65.46» '152.60»'155»   '963»   '348»   '173»   '26744» '496 |
| 26 |  | '2024-09-13»'1779-04»   'lib941»'1779-04\_lib941»'1703446»   '1687346»   '99.05» '4411532»   '65.61» '4391472»   '1.00»  '65.58» '83.11» '85»'4312865»   '0.98»  '65.39» '84.44» '85»'552»   '174»   '73»'20060» '304 |  | 26 |  | '2024-09-17»'1779-04»   'lib941»'1779-04\_lib941»'1703446»   '1687346»   '99.05» '4411532»   '65.61» '4391472»   '1.00»  '65.58» '83.11» '85»'4312865»   '0.98»  '65.39» '84.44» '85»'552»   '174»   '73»'20060» '304 |  | 26 |  | '2024-09-18»'1779-04»   'lib941»'1779-04\_lib941»'1703446»   '1687346»   '99.05» '4411532»   '65.61» '4391472»   '1.00»  '65.58» '83.11» '85»'4312865»   '0.98»  '65.39» '84.44» '85»'552»   '174»   '73»'20060» '304 |
| 27 |  | '2024-09-13»'1780-04»   'lib942»'1780-04\_lib942»'771658»'764254»'99.04» '4411532»   '65.61» '4378035»   '0.99»  '65.54» '38.25» '38»'4226176»   '0.96»  '65.24» '39.29» '39»'520»   '106»   '82»'33497» '283 |  | 27 |  | '2024-09-17»'1780-04»   'lib942»'1780-04\_lib942»'771658»'764254»'99.04» '4411532»   '65.61» '4378035»   '0.99»  '65.54» '38.25» '38»'4226176»   '0.96»  '65.24» '39.29» '39»'520»   '106»   '82»'33497» '283 |  | 27 |  | '2024-09-18»'1780-04»   'lib942»'1780-04\_lib942»'771658»'764254»'99.04» '4411532»   '65.61» '4378035»   '0.99»  '65.54» '38.25» '38»'4226176»   '0.96»  '65.24» '39.29» '39»'520»   '106»   '82»'33497» '283 |
| 28 |  | '2024-09-13»'1783-04»   'lib936»'1783-04\_lib936»'709561»'698829»'98.49» '4411532»   '65.61» '4377912»   '0.99»  '65.54» '33.42» '33»'4212737»   '0.95»  '65.27» '34.39» '34»'938»   '243»   '107»   '33620» '475 |  | 28 |  | '2024-09-17»'1783-04»   'lib936»'1783-04\_lib936»'709561»'698829»'98.49» '4411532»   '65.61» '4377912»   '0.99»  '65.54» '33.42» '33»'4212737»   '0.95»  '65.27» '34.39» '34»'938»   '243»   '107»   '33620» '475 |  | 28 |  | '2024-09-18»'1783-04»   'lib936»'1783-04\_lib936»'709561»'698829»'98.49» '4411532»   '65.61» '4377912»   '0.99»  '65.54» '33.42» '33»'4212737»   '0.95»  '65.27» '34.39» '34»'938»   '243»   '107»   '33620» '475 |
| 29 |  | '2024-09-13»'2509-04»   'lib1471»   '2509-04\_lib1471»   '1682495»   '1674087»   '99.50» '4411532»   '65.61» '4380451»   '0.99»  '65.57» '99.52» '101»   '4336948»   '0.98»  '65.47» '100.44»'101»   '951»   '571»   '194»   '31081» '503 |  | 29 |  | '2024-09-17»'2509-04»   'lib1471»   '2509-04\_lib1471»   '1682495»   '1674087»   '99.50» '4411532»   '65.61» '4380451»   '0.99»  '65.57» '99.52» '101»   '4336948»   '0.98»  '65.47» '100.44»'101»   '951»   '571»   '194»   '31081» '503 |  | 29 |  | '2024-09-18»'2509-04»   'lib1471»   '2509-04\_lib1471»   '1682495»   '1674087»   '99.50» '4411532»   '65.61» '4380451»   '0.99»  '65.57» '99.52» '101»   '4336948»   '0.98»  '65.47» '100.44»'101»   '951»   '571»   '194»   '31081» '503 |
| 30 |  | '2024-09-13»'3154-04»   'lib905»'3154-04\_lib905»'2667188»   '2639376»   '98.96» '4411532»   '65.61» '4382200»   '0.99»  '65.59» '130.72»'134»   '4325828»   '0.98»  '65.44» '132.28»'135»   '960»   '343»   '125»   '29332» '494 |  | 30 |  | '2024-09-17»'3154-04»   'lib905»'3154-04\_lib905»'2667188»   '2639376»   '98.96» '4411532»   '65.61» '4382200»   '0.99»  '65.59» '130.72»'134»   '4325828»   '0.98»  '65.44» '132.28»'135»   '960»   '343»   '125»   '29332» '494 |  | 30 |  | '2024-09-18»'3154-04»   'lib905»'3154-04\_lib905»'2667188»   '2639376»   '98.96» '4411532»   '65.61» '4382200»   '0.99»  '65.59» '130.72»'134»   '4325828»   '0.98»  '65.44» '132.28»'135»   '960»   '343»   '125»   '29332» '494 |
| 31 |  | '2024-09-13»'3156-04»   'lib926»'3156-04\_lib926»'380483»'377451»'99.20» '4411532»   '65.61» '4353802»   '0.99»  '65.51» '19.84» '20»'3913362»   '0.89»  '64.99» '21.13» '21»'871»   '167»   '123»   '57730» '444 |  | 31 |  | '2024-09-17»'3156-04»   'lib926»'3156-04\_lib926»'380483»'377451»'99.20» '4411532»   '65.61» '4353802»   '0.99»  '65.51» '19.84» '20»'3913362»   '0.89»  '64.99» '21.13» '21»'871»   '167»   '123»   '57730» '444 |  | 31 |  | '2024-09-18»'3156-04»   'lib926»'3156-04\_lib926»'380483»'377451»'99.20» '4411532»   '65.61» '4353802»   '0.99»  '65.51» '19.84» '20»'3913362»   '0.89»  '64.99» '21.13» '21»'871»   '167»   '123»   '57730» '444 |
| 32 |  | '2024-09-13»'3158-04»   'lib943»'3158-04\_lib943»'1707041»   '1695432»   '99.32» '4411532»   '65.61» '4389575»   '1.00»  '65.57» '86.22» '88»'4315850»   '0.98»  '65.37» '87.52» '89»'536»   '233»   '92»'21957» '298 |  | 32 |  | '2024-09-17»'3158-04»   'lib943»'3158-04\_lib943»'1707041»   '1695432»   '99.32» '4411532»   '65.61» '4389575»   '1.00»  '65.57» '86.22» '88»'4315850»   '0.98»  '65.37» '87.52» '89»'536»   '233»   '92»'21957» '298 |  | 32 |  | '2024-09-18»'3158-04»   'lib943»'3158-04\_lib943»'1707041»   '1695432»   '99.32» '4411532»   '65.61» '4389575»   '1.00»  '65.57» '86.22» '88»'4315850»   '0.98»  '65.37» '87.52» '89»'536»   '233»   '92»'21957» '298 |
| 33 |  | '2024-09-13»'3160-04»   'lib958»'3160-04\_lib958»'2300674»   '2276012»   '98.93» '4411532»   '65.61» '4358035»   '0.99»  '65.57» '113.17»'116»   '4304146»   '0.98»  '65.42» '114.45»'116»   '2255»  '438»   '265»   '53497» '1222 |  | 33 |  | '2024-09-17»'3160-04»   'lib958»'3160-04\_lib958»'2300674»   '2276012»   '98.93» '4411532»   '65.61» '4358035»   '0.99»  '65.57» '113.17»'116»   '4304146»   '0.98»  '65.42» '114.45»'116»   '2255»  '438»   '265»   '53497» '1222 |  | 33 |  | '2024-09-18»'3160-04»   'lib958»'3160-04\_lib958»'2300674»   '2276012»   '98.93» '4411532»   '65.61» '4358035»   '0.99»  '65.57» '113.17»'116»   '4304146»   '0.98»  '65.42» '114.45»'116»   '2255»  '438»   '265»   '53497» '1222 |
| 34 |  | '2024-09-13»'3491-04»   'lib897»'3491-04\_lib897»'844550»'832403»'98.56» '4411532»   '65.61» '4366313»   '0.99»  '65.54» '42.86» '43»'4210470»   '0.95»  '65.26» '44.14» '44»'2045»  '337»   '265»   '45219» '1111 |  | 34 |  | '2024-09-17»'3491-04»   'lib897»'3491-04\_lib897»'844550»'832403»'98.56» '4411532»   '65.61» '4366313»   '0.99»  '65.54» '42.86» '43»'4210470»   '0.95»  '65.26» '44.14» '44»'2045»  '337»   '265»   '45219» '1111 |  | 34 |  | '2024-09-18»'3491-04»   'lib897»'3491-04\_lib897»'844550»'832403»'98.56» '4411532»   '65.61» '4366313»   '0.99»  '65.54» '42.86» '43»'4210470»   '0.95»  '65.26» '44.14» '44»'2045»  '337»   '265»   '45219» '1111 |
| 35 |  | '2024-09-13»'3494-04»   'lib953»'3494-04\_lib953»'2202575»   '2177642»   '98.87» '4411532»   '65.61» '4380286»   '0.99»  '65.56» '107.46»'110»   '4316417»   '0.98»  '65.40» '108.87»'110»   '2194»  '538»   '216»   '31246» '1187 |  | 35 |  | '2024-09-17»'3494-04»   'lib953»'3494-04\_lib953»'2202575»   '2177642»   '98.87» '4411532»   '65.61» '4380286»   '0.99»  '65.56» '107.46»'110»   '4316417»   '0.98»  '65.40» '108.87»'110»   '2194»  '538»   '216»   '31246» '1187 |  | 35 |  | '2024-09-18»'3494-04»   'lib953»'3494-04\_lib953»'2202575»   '2177642»   '98.87» '4411532»   '65.61» '4380286»   '0.99»  '65.56» '107.46»'110»   '4316417»   '0.98»  '65.40» '108.87»'110»   '2194»  '538»   '216»   '31246» '1187 |
| 36 |  | '2024-09-13»'3496-04»   'lib906»'3496-04\_lib906»'2082097»   '2065440»   '99.20» '4411532»   '65.61» '4381850»   '0.99»  '65.58» '106.31»'109»   '4318086»   '0.98»  '65.41» '107.73»'109»   '953»   '395»   '202»   '29682» '494 |  | 36 |  | '2024-09-17»'3496-04»   'lib906»'3496-04\_lib906»'2082097»   '2065440»   '99.20» '4411532»   '65.61» '4381850»   '0.99»  '65.58» '106.31»'109»   '4318086»   '0.98»  '65.41» '107.73»'109»   '953»   '395»   '202»   '29682» '494 |  | 36 |  | '2024-09-18»'3496-04»   'lib906»'3496-04\_lib906»'2082097»   '2065440»   '99.20» '4411532»   '65.61» '4381850»   '0.99»  '65.58» '106.31»'109»   '4318086»   '0.98»  '65.41» '107.73»'109»   '953»   '395»   '202»   '29682» '494 |
| 37 |  | '2024-09-13»'3497-04»   'lib927»'3497-04\_lib927»'449971»'446794»'99.29» '4411532»   '65.61» '4369269»   '0.99»  '65.52» '23.06» '23»'4062092»   '0.92»  '65.15» '24.17» '24»'491»   '94»'41»'42263» '252 |  | 37 |  | '2024-09-17»'3497-04»   'lib927»'3497-04\_lib927»'449971»'446794»'99.29» '4411532»   '65.61» '4369269»   '0.99»  '65.52» '23.06» '23»'4062092»   '0.92»  '65.15» '24.17» '24»'491»   '94»'41»'42263» '252 |  | 37 |  | '2024-09-18»'3497-04»   'lib927»'3497-04\_lib927»'449971»'446794»'99.29» '4411532»   '65.61» '4369269»   '0.99»  '65.52» '23.06» '23»'4062092»   '0.92»  '65.15» '24.17» '24»'491»   '94»'41»'42263» '252 |
| 38 |  | '2024-09-13»'3734-04»   'lib895»'3734-04\_lib895»'1318961»   '1310499»   '99.36» '4411532»   '65.61» '4369011»   '0.99»  '65.56» '67.44» '68»'4284525»   '0.97»  '65.35» '68.61» '68»'822»   '414»   '105»   '42521» '413 |  | 38 |  | '2024-09-17»'3734-04»   'lib895»'3734-04\_lib895»'1318961»   '1310499»   '99.36» '4411532»   '65.61» '4369011»   '0.99»  '65.56» '67.44» '68»'4284525»   '0.97»  '65.35» '68.61» '68»'822»   '414»   '105»   '42521» '413 |  | 38 |  | '2024-09-18»'3734-04»   'lib895»'3734-04\_lib895»'1318961»   '1310499»   '99.36» '4411532»   '65.61» '4369011»   '0.99»  '65.56» '67.44» '68»'4284525»   '0.97»  '65.35» '68.61» '68»'822»   '414»   '105»   '42521» '413 |
| 39 |  | '2024-09-13»'3736-04»   'lib937»'3736-04\_lib937»'903487»'893081»'98.85» '4411532»   '65.61» '4355294»   '0.99»  '65.55» '43.28» '44»'4224714»   '0.96»  '65.30» '44.34» '44»'936»   '312»   '125»   '56238» '482 |  | 39 |  | '2024-09-17»'3736-04»   'lib937»'3736-04\_lib937»'903487»'893081»'98.85» '4411532»   '65.61» '4355294»   '0.99»  '65.55» '43.28» '44»'4224714»   '0.96»  '65.30» '44.34» '44»'936»   '312»   '125»   '56238» '482 |  | 39 |  | '2024-09-18»'3736-04»   'lib937»'3736-04\_lib937»'903487»'893081»'98.85» '4411532»   '65.61» '4355294»   '0.99»  '65.55» '43.28» '44»'4224714»   '0.96»  '65.30» '44.34» '44»'936»   '312»   '125»   '56238» '482 |
| 40 |  | '2024-09-13»'3859-03»   'lib921»'3859-03\_lib921»'1651781»   '1569432»   '95.01» '4411532»   '65.61» '4387205»   '0.99»  '65.57» '78.57» '80»'4308801»   '0.98»  '65.36» '79.82» '80»'975»   '259»   '155»   '24327» '486 |  | 40 |  | '2024-09-17»'3859-03»   'lib921»'3859-03\_lib921»'1651781»   '1569432»   '95.01» '4411532»   '65.61» '4387205»   '0.99»  '65.57» '78.57» '80»'4308801»   '0.98»  '65.36» '79.82» '80»'975»   '259»   '155»   '24327» '486 |  | 40 |  | '2024-09-18»'3859-03»   'lib921»'3859-03\_lib921»'1651781»   '1569432»   '95.01» '4411532»   '65.61» '4387205»   '0.99»  '65.57» '78.57» '80»'4308801»   '0.98»  '65.36» '79.82» '80»'975»   '259»   '155»   '24327» '486 |
| 41 |  | '2024-09-13»'3861-03»   'lib922»'3861-03\_lib922»'573292»'568685»'99.20» '4411532»   '65.61» '4358111»   '0.99»  '65.53» '29.67» '29»'4194859»   '0.95»  '65.23» '30.53» '30»'743»   '136»   '64»'53421» '392 |  | 41 |  | '2024-09-17»'3861-03»   'lib922»'3861-03\_lib922»'573292»'568685»'99.20» '4411532»   '65.61» '4358111»   '0.99»  '65.53» '29.67» '29»'4194859»   '0.95»  '65.23» '30.53» '30»'743»   '136»   '64»'53421» '392 |  | 41 |  | '2024-09-18»'3861-03»   'lib922»'3861-03\_lib922»'573292»'568685»'99.20» '4411532»   '65.61» '4358111»   '0.99»  '65.53» '29.67» '29»'4194859»   '0.95»  '65.23» '30.53» '30»'743»   '136»   '64»'53421» '392 |
| 42 |  | '2024-09-13»'3865-03»   'lib971»'3865-03\_lib971»'4381570»   '4355626»   '99.41» '4411532»   '65.61» '4386139»   '0.99»  '65.59» '220.42»'226»   '4344870»   '0.98»  '65.47» '222.38»'227»   '931»   '409»   '154»   '25393» '502 |  | 42 |  | '2024-09-17»'3865-03»   'lib971»'3865-03\_lib971»'4381570»   '4355626»   '99.41» '4411532»   '65.61» '4386139»   '0.99»  '65.59» '220.42»'226»   '4344870»   '0.98»  '65.47» '222.38»'227»   '931»   '409»   '154»   '25393» '502 |  | 42 |  | '2024-09-18»'3865-03»   'lib971»'3865-03\_lib971»'4381570»   '4355626»   '99.41» '4411532»   '65.61» '4386139»   '0.99»  '65.59» '220.42»'226»   '4344870»   '0.98»  '65.47» '222.38»'227»   '931»   '409»   '154»   '25393» '502 |
| 43 |  | '2024-09-13»'4139-04»   'lib959»'4139-04\_lib959»'4160475»   '4121097»   '99.05» '4411532»   '65.61» '4356299»   '0.99»  '65.57» '209.11»'215»   '4316140»   '0.98»  '65.45» '210.92»'215»   '2295»  '405»   '240»   '55233» '1220 |  | 43 |  | '2024-09-17»'4139-04»   'lib959»'4139-04\_lib959»'4160475»   '4121097»   '99.05» '4411532»   '65.61» '4356299»   '0.99»  '65.57» '209.11»'215»   '4316140»   '0.98»  '65.45» '210.92»'215»   '2295»  '405»   '240»   '55233» '1220 |  | 43 |  | '2024-09-18»'4139-04»   'lib959»'4139-04\_lib959»'4160475»   '4121097»   '99.05» '4411532»   '65.61» '4356299»   '0.99»  '65.57» '209.11»'215»   '4316140»   '0.98»  '65.45» '210.92»'215»   '2295»  '405»   '240»   '55233» '1220 |
| 44 |  | '2024-09-13»'4145-04»   'lib919»'4145-04\_lib919»'556645»'553061»'99.36» '4411532»   '65.61» '4361454»   '0.99»  '65.53» '28.81» '29»'4202539»   '0.95»  '65.23» '29.64» '29»'809»   '207»   '117»   '50078» '430 |  | 44 |  | '2024-09-17»'4145-04»   'lib919»'4145-04\_lib919»'556645»'553061»'99.36» '4411532»   '65.61» '4361454»   '0.99»  '65.53» '28.81» '29»'4202539»   '0.95»  '65.23» '29.64» '29»'809»   '207»   '117»   '50078» '430 |  | 44 |  | '2024-09-18»'4145-04»   'lib919»'4145-04\_lib919»'556645»'553061»'99.36» '4411532»   '65.61» '4361454»   '0.99»  '65.53» '28.81» '29»'4202539»   '0.95»  '65.23» '29.64» '29»'809»   '207»   '117»   '50078» '430 |
| 45 |  | '2024-09-13»'4148-04»   'lib907»'4148-04\_lib907»'2694074»   '2673842»   '99.25» '4411532»   '65.61» '4384514»   '0.99»  '65.59» '135.70»'139»   '4334802»   '0.98»  '65.45» '137.12»'139»   '963»   '350»   '107»   '27018» '501 |  | 45 |  | '2024-09-17»'4148-04»   'lib907»'4148-04\_lib907»'2694074»   '2673842»   '99.25» '4411532»   '65.61» '4384514»   '0.99»  '65.59» '135.70»'139»   '4334802»   '0.98»  '65.45» '137.12»'139»   '963»   '350»   '107»   '27018» '501 |  | 45 |  | '2024-09-18»'4148-04»   'lib907»'4148-04\_lib907»'2694074»   '2673842»   '99.25» '4411532»   '65.61» '4384514»   '0.99»  '65.59» '135.70»'139»   '4334802»   '0.98»  '65.45» '137.12»'139»   '963»   '350»   '107»   '27018» '501 |
| 46 |  | '2024-09-13»'420-04»'lib935»'420-04\_lib935» '1683527»   '1671584»   '99.29» '4411532»   '65.61» '4388296»   '0.99»  '65.58» '83.90» '86»'4316083»   '0.98»  '65.39» '85.15» '86»'989»   '286»   '123»   '23236» '503 |  | 46 |  | '2024-09-17»'420-04»'lib935»'420-04\_lib935» '1683527»   '1671584»   '99.29» '4411532»   '65.61» '4388296»   '0.99»  '65.58» '83.90» '86»'4316083»   '0.98»  '65.39» '85.15» '86»'989»   '286»   '123»   '23236» '503 |  | 46 |  | '2024-09-18»'420-04»'lib935»'420-04\_lib935» '1683527»   '1671584»   '99.29» '4411532»   '65.61» '4388296»   '0.99»  '65.58» '83.90» '86»'4316083»   '0.98»  '65.39» '85.15» '86»'989»   '286»   '123»   '23236» '503 |
| 47 |  | '2024-09-13»'421-04»'lib956»'421-04\_lib956» '865157»'856226»'98.97» '4411532»   '65.61» '4342107»   '0.98»  '65.52» '43.52» '44»'4190718»   '0.95»  '65.25» '44.73» '45»'2162»  '331»   '231»   '69425» '1183 |  | 47 |  | '2024-09-17»'421-04»'lib956»'421-04\_lib956» '865157»'856226»'98.97» '4411532»   '65.61» '4342107»   '0.98»  '65.52» '43.52» '44»'4190718»   '0.95»  '65.25» '44.73» '45»'2162»  '331»   '231»   '69425» '1183 |  | 47 |  | '2024-09-18»'421-04»'lib956»'421-04\_lib956» '865157»'856226»'98.97» '4411532»   '65.61» '4342107»   '0.98»  '65.52» '43.52» '44»'4190718»   '0.95»  '65.25» '44.73» '45»'2162»  '331»   '231»   '69425» '1183 |
| 48 |  | '2024-09-13»'4514-03»   'lib901»'4514-03\_lib901»'137567»'136537»'99.25» '4411532»   '65.61» '4292856»   '0.97»  '65.43» '6.97»  '7» '1216194»   '0.28»  '63.75» '11.12» '11»'252»   '10»'16»'118676»'131 |  | 48 |  | '2024-09-17»'4514-03»   'lib901»'4514-03\_lib901»'137567»'136537»'99.25» '4411532»   '65.61» '4292856»   '0.97»  '65.43» '6.97»  '7» '1216194»   '0.28»  '63.75» '11.12» '11»'252»   '10»'16»'118676»'131 |  | 48 |  | '2024-09-18»'4514-03»   'lib901»'4514-03\_lib901»'137567»'136537»'99.25» '4411532»   '65.61» '4292856»   '0.97»  '65.43» '6.97»  '7» '1216194»   '0.28»  '63.75» '11.12» '11»'252»   '10»'16»'118676»'131 |
| 49 |  | '2024-09-13»'4516-03»   'lib947»'4516-03\_lib947»'1903453»   '1887326»   '99.15» '4411532»   '65.61» '4379445»   '0.99»  '65.58» '95.41» '97»'4309152»   '0.98»  '65.40» '96.81» '98»'917»   '297»   '192»   '32087» '479 |  | 49 |  | '2024-09-17»'4516-03»   'lib947»'4516-03\_lib947»'1903453»   '1887326»   '99.15» '4411532»   '65.61» '4379445»   '0.99»  '65.58» '95.41» '97»'4309152»   '0.98»  '65.40» '96.81» '98»'917»   '297»   '192»   '32087» '479 |  | 49 |  | '2024-09-18»'4516-03»   'lib947»'4516-03\_lib947»'1903453»   '1887326»   '99.15» '4411532»   '65.61» '4379445»   '0.99»  '65.58» '95.41» '97»'4309152»   '0.98»  '65.40» '96.81» '98»'917»   '297»   '192»   '32087» '479 |
| 50 |  | '2024-09-13»'4518-03»   'lib949»'4518-03\_lib949»'2208395»   '2186862»   '99.02» '4411532»   '65.61» '4366063»   '0.99»  '65.56» '110.73»'112»   '4304472»   '0.98»  '65.40» '112.15»'113»   '2133»  '658»   '277»   '45469» '1144 |  | 50 |  | '2024-09-17»'4518-03»   'lib949»'4518-03\_lib949»'2208395»   '2186862»   '99.02» '4411532»   '65.61» '4366063»   '0.99»  '65.56» '110.73»'112»   '4304472»   '0.98»  '65.40» '112.15»'113»   '2133»  '658»   '277»   '45469» '1144 |  | 50 |  | '2024-09-18»'4518-03»   'lib949»'4518-03\_lib949»'2208395»   '2186862»   '99.02» '4411532»   '65.61» '4366063»   '0.99»  '65.56» '110.73»'112»   '4304472»   '0.98»  '65.40» '112.15»'113»   '2133»  '658»   '277»   '45469» '1144 |
| 51 |  | '2024-09-13»'4523-03»   'lib972»'4523-03\_lib972»'2988519»   '2967758»   '99.31» '4411532»   '65.61» '4378344»   '0.99»  '65.59» '150.35»'154»   '4325524»   '0.98»  '65.45» '152.05»'154»   '865»   '381»   '118»   '33188» '455 |  | 51 |  | '2024-09-17»'4523-03»   'lib972»'4523-03\_lib972»'2988519»   '2967758»   '99.31» '4411532»   '65.61» '4378344»   '0.99»  '65.59» '150.35»'154»   '4325524»   '0.98»  '65.45» '152.05»'154»   '865»   '381»   '118»   '33188» '455 |  | 51 |  | '2024-09-18»'4523-03»   'lib972»'4523-03\_lib972»'2988519»   '2967758»   '99.31» '4411532»   '65.61» '4378344»   '0.99»  '65.59» '150.35»'154»   '4325524»   '0.98»  '65.45» '152.05»'154»   '865»   '381»   '118»   '33188» '455 |
| 52 |  | '2024-09-13»'4712-04»   'lib960»'4712-04\_lib960»'1509501»   '1492091»   '98.85» '4411532»   '65.61» '4347673»   '0.99»  '65.54» '75.45» '77»'4266829»   '0.97»  '65.35» '76.68» '77»'2197»  '417»   '252»   '63859» '1173 |  | 52 |  | '2024-09-17»'4712-04»   'lib960»'4712-04\_lib960»'1509501»   '1492091»   '98.85» '4411532»   '65.61» '4347673»   '0.99»  '65.54» '75.45» '77»'4266829»   '0.97»  '65.35» '76.68» '77»'2197»  '417»   '252»   '63859» '1173 |  | 52 |  | '2024-09-18»'4712-04»   'lib960»'4712-04\_lib960»'1509501»   '1492091»   '98.85» '4411532»   '65.61» '4347673»   '0.99»  '65.54» '75.45» '77»'4266829»   '0.97»  '65.35» '76.68» '77»'2197»  '417»   '252»   '63859» '1173 |
| 53 |  | '2024-09-13»'4714-04»   'lib1472»   '4714-04\_lib1472»   '1901485»   '1891536»   '99.48» '4411532»   '65.61» '4391028»   '1.00»  '65.58» '111.31»'112»   '4356797»   '0.99»  '65.49» '112.13»'113»   '895»   '248»   '110»   '20504» '456 |  | 53 |  | '2024-09-17»'4714-04»   'lib1472»   '4714-04\_lib1472»   '1901485»   '1891536»   '99.48» '4411532»   '65.61» '4391028»   '1.00»  '65.58» '111.31»'112»   '4356797»   '0.99»  '65.49» '112.13»'113»   '895»   '248»   '110»   '20504» '456 |  | 53 |  | '2024-09-18»'4714-04»   'lib1472»   '4714-04\_lib1472»   '1901485»   '1891536»   '99.48» '4411532»   '65.61» '4391028»   '1.00»  '65.58» '111.31»'112»   '4356797»   '0.99»  '65.49» '112.13»'113»   '895»   '248»   '110»   '20504» '456 |
| 54 |  | '2024-09-13»'4717-04»   'lib898»'4717-04\_lib898»'1405181»   '1389480»   '98.88» '4411532»   '65.61» '4370582»   '0.99»  '65.56» '70.49» '71»'4269358»   '0.97»  '65.35» '71.96» '72»'2117»  '290»   '253»   '40950» '1145 |  | 54 |  | '2024-09-17»'4717-04»   'lib898»'4717-04\_lib898»'1405181»   '1389480»   '98.88» '4411532»   '65.61» '4370582»   '0.99»  '65.56» '70.49» '71»'4269358»   '0.97»  '65.35» '71.96» '72»'2117»  '290»   '253»   '40950» '1145 |  | 54 |  | '2024-09-18»'4717-04»   'lib898»'4717-04\_lib898»'1405181»   '1389480»   '98.88» '4411532»   '65.61» '4370582»   '0.99»  '65.56» '70.49» '71»'4269358»   '0.97»  '65.35» '71.96» '72»'2117»  '290»   '253»   '40950» '1145 |
| 55 |  | '2024-09-13»'4724-03»   'lib1517»   '4724-03\_lib1517»   '1601083»   '1593419»   '99.52» '4411532»   '65.61» '4364979»   '0.99»  '65.58» '88.97» '91»'4314420»   '0.98»  '65.47» '89.93» '91»'1465»  '512»   '284»   '46553» '756 |  | 55 |  | '2024-09-17»'4724-03»   'lib1517»   '4724-03\_lib1517»   '1601083»   '1593419»   '99.52» '4411532»   '65.61» '4364979»   '0.99»  '65.58» '88.97» '91»'4314420»   '0.98»  '65.47» '89.93» '91»'1465»  '512»   '284»   '46553» '756 |  | 55 |  | '2024-09-18»'4724-03»   'lib1517»   '4724-03\_lib1517»   '1601083»   '1593419»   '99.52» '4411532»   '65.61» '4364979»   '0.99»  '65.58» '88.97» '91»'4314420»   '0.98»  '65.47» '89.93» '91»'1465»  '512»   '284»   '46553» '756 |
| 56 |  | '2024-09-13»'4779-04»   'lib944»'4779-04\_lib944»'1218988»   '1185183»   '97.23» '4411532»   '65.61» '4386347»   '0.99»  '65.56» '59.66» '60»'4295236»   '0.97»  '65.33» '60.76» '61»'564»   '244»   '50»'25185» '301 |  | 56 |  | '2024-09-17»'4779-04»   'lib944»'4779-04\_lib944»'1218988»   '1185183»   '97.23» '4411532»   '65.61» '4386347»   '0.99»  '65.56» '59.66» '60»'4295236»   '0.97»  '65.33» '60.76» '61»'564»   '244»   '50»'25185» '301 |  | 56 |  | '2024-09-18»'4779-04»   'lib944»'4779-04\_lib944»'1218988»   '1185183»   '97.23» '4411532»   '65.61» '4386347»   '0.99»  '65.56» '59.66» '60»'4295236»   '0.97»  '65.33» '60.76» '61»'564»   '244»   '50»'25185» '301 |
| 57 |  | '2024-09-13»'4781-04»   'lib908»'4781-04\_lib908»'2487918»   '2466911»   '99.16» '4411532»   '65.61» '4385240»   '0.99»  '65.59» '124.09»'127»   '4334184»   '0.98»  '65.44» '125.42»'127»   '942»   '441»   '167»   '26292» '472 |  | 57 |  | '2024-09-17»'4781-04»   'lib908»'4781-04\_lib908»'2487918»   '2466911»   '99.16» '4411532»   '65.61» '4385240»   '0.99»  '65.59» '124.09»'127»   '4334184»   '0.98»  '65.44» '125.42»'127»   '942»   '441»   '167»   '26292» '472 |  | 57 |  | '2024-09-18»'4781-04»   'lib908»'4781-04\_lib908»'2487918»   '2466911»   '99.16» '4411532»   '65.61» '4385240»   '0.99»  '65.59» '124.09»'127»   '4334184»   '0.98»  '65.44» '125.42»'127»   '942»   '441»   '167»   '26292» '472 |
| 58 |  | '2024-09-13»'4783-04»   'lib896»'4783-04\_lib896»'716744»'710103»'99.07» '4411532»   '65.61» '4348165»   '0.99»  '65.52» '36.44» '37»'4185636»   '0.95»  '65.25» '37.49» '37»'784»   '218»   '120»   '63367» '396 |  | 58 |  | '2024-09-17»'4783-04»   'lib896»'4783-04\_lib896»'716744»'710103»'99.07» '4411532»   '65.61» '4348165»   '0.99»  '65.52» '36.44» '37»'4185636»   '0.95»  '65.25» '37.49» '37»'784»   '218»   '120»   '63367» '396 |  | 58 |  | '2024-09-18»'4783-04»   'lib896»'4783-04\_lib896»'716744»'710103»'99.07» '4411532»   '65.61» '4348165»   '0.99»  '65.52» '36.44» '37»'4185636»   '0.95»  '65.25» '37.49» '37»'784»   '218»   '120»   '63367» '396 |
| 59 |  | '2024-09-13»'4785-04»   'lib909»'4785-04\_lib909»'2283007»   '2261099»   '99.04» '4411532»   '65.61» '4383209»   '0.99»  '65.59» '115.20»'118»   '4322364»   '0.98»  '65.43» '116.67»'118»   '956»   '320»   '216»   '28323» '485 |  | 59 |  | '2024-09-17»'4785-04»   'lib909»'4785-04\_lib909»'2283007»   '2261099»   '99.04» '4411532»   '65.61» '4383209»   '0.99»  '65.59» '115.20»'118»   '4322364»   '0.98»  '65.43» '116.67»'118»   '956»   '320»   '216»   '28323» '485 |  | 59 |  | '2024-09-18»'4785-04»   'lib909»'4785-04\_lib909»'2283007»   '2261099»   '99.04» '4411532»   '65.61» '4383209»   '0.99»  '65.59» '115.20»'118»   '4322364»   '0.98»  '65.43» '116.67»'118»   '956»   '320»   '216»   '28323» '485 |
| 60 |  | '2024-09-13»'5248-04»   'lib928»'5248-04\_lib928»'2060766»   '2040535»   '99.02» '4411532»   '65.61» '4398778»   '1.00»  '65.57» '99.72» '102»   '4322564»   '0.98»  '65.41» '101.30»'103»   '1001»  '304»   '129»   '12754» '510 |  | 60 |  | '2024-09-17»'5248-04»   'lib928»'5248-04\_lib928»'2060766»   '2040535»   '99.02» '4411532»   '65.61» '4398778»   '1.00»  '65.57» '99.72» '102»   '4322564»   '0.98»  '65.41» '101.30»'103»   '1001»  '304»   '129»   '12754» '510 |  | 60 |  | '2024-09-18»'5248-04»   'lib928»'5248-04\_lib928»'2060766»   '2040535»   '99.02» '4411532»   '65.61» '4398778»   '1.00»  '65.57» '99.72» '102»   '4322564»   '0.98»  '65.41» '101.30»'103»   '1001»  '304»   '129»   '12754» '510 |
| 61 |  | '2024-09-13»'5253-04»   'lib961»'5253-04\_lib961»'1605150»   '1584368»   '98.71» '4411532»   '65.61» '4349394»   '0.99»  '65.55» '80.52» '82»'4273595»   '0.97»  '65.35» '81.78» '82»'2236»  '463»   '284»   '62138» '1192 |  | 61 |  | '2024-09-17»'5253-04»   'lib961»'5253-04\_lib961»'1605150»   '1584368»   '98.71» '4411532»   '65.61» '4349394»   '0.99»  '65.55» '80.52» '82»'4273595»   '0.97»  '65.35» '81.78» '82»'2236»  '463»   '284»   '62138» '1192 |  | 61 |  | '2024-09-18»'5253-04»   'lib961»'5253-04\_lib961»'1605150»   '1584368»   '98.71» '4411532»   '65.61» '4349394»   '0.99»  '65.55» '80.52» '82»'4273595»   '0.97»  '65.35» '81.78» '82»'2236»  '463»   '284»   '62138» '1192 |
| 62 |  | '2024-09-13»'5468-03»   'lib912»'5468-03\_lib912»'476720»'473136»'99.25» '4411532»   '65.61» '4353287»   '0.99»  '65.50» '24.78» '25»'4130455»   '0.94»  '65.19» '25.71» '25»'826»   '391»   '141»   '58245» '447 |  | 62 |  | '2024-09-17»'5468-03»   'lib912»'5468-03\_lib912»'476720»'473136»'99.25» '4411532»   '65.61» '4353287»   '0.99»  '65.50» '24.78» '25»'4130455»   '0.94»  '65.19» '25.71» '25»'826»   '391»   '141»   '58245» '447 |  | 62 |  | '2024-09-18»'5468-03»   'lib912»'5468-03\_lib912»'476720»'473136»'99.25» '4411532»   '65.61» '4353287»   '0.99»  '65.50» '24.78» '25»'4130455»   '0.94»  '65.19» '25.71» '25»'826»   '391»   '141»   '58245» '447 |
| 63 |  | '2024-09-13»'5472-03»   'lib938»'5472-03\_lib938»'1348743»   '1335725»   '99.03» '4411532»   '65.61» '4379139»   '0.99»  '65.58» '65.97» '67»'4284572»   '0.97»  '65.36» '67.22» '67»'555»   '173»   '80»'32393» '305 |  | 63 |  | '2024-09-17»'5472-03»   'lib938»'5472-03\_lib938»'1348743»   '1335725»   '99.03» '4411532»   '65.61» '4379139»   '0.99»  '65.58» '65.97» '67»'4284572»   '0.97»  '65.36» '67.22» '67»'555»   '173»   '80»'32393» '305 |  | 63 |  | '2024-09-18»'5472-03»   'lib938»'5472-03\_lib938»'1348743»   '1335725»   '99.03» '4411532»   '65.61» '4379139»   '0.99»  '65.58» '65.97» '67»'4284572»   '0.97»  '65.36» '67.22» '67»'555»   '173»   '80»'32393» '305 |
| 64 |  | '2024-09-13»'5685-04»   'lib945»'5685-04\_lib945»'3097896»   '3081287»   '99.46» '4411532»   '65.61» '4394668»   '1.00»  '65.59» '150.17»'154»   '4351412»   '0.99»  '65.46» '151.56»'155»   '564»   '215»   '69»'16864» '319 |  | 64 |  | '2024-09-17»'5685-04»   'lib945»'5685-04\_lib945»'3097896»   '3081287»   '99.46» '4411532»   '65.61» '4394668»   '1.00»  '65.59» '150.17»'154»   '4351412»   '0.99»  '65.46» '151.56»'155»   '564»   '215»   '69»'16864» '319 |  | 64 |  | '2024-09-18»'5685-04»   'lib945»'5685-04\_lib945»'3097896»   '3081287»   '99.46» '4411532»   '65.61» '4394668»   '1.00»  '65.59» '150.17»'154»   '4351412»   '0.99»  '65.46» '151.56»'155»   '564»   '215»   '69»'16864» '319 |
| 65 |  | '2024-09-13»'5687-04»   'lib946»'5687-04\_lib946»'2444014»   '2421258»   '99.07» '4411532»   '65.61» '4393421»   '1.00»  '65.59» '120.32»'123»   '4338263»   '0.98»  '65.43» '121.72»'123»   '570»   '186»   '52»'18111» '316 |  | 65 |  | '2024-09-17»'5687-04»   'lib946»'5687-04\_lib946»'2444014»   '2421258»   '99.07» '4411532»   '65.61» '4393421»   '1.00»  '65.59» '120.32»'123»   '4338263»   '0.98»  '65.43» '121.72»'123»   '570»   '186»   '52»'18111» '316 |  | 65 |  | '2024-09-18»'5687-04»   'lib946»'5687-04\_lib946»'2444014»   '2421258»   '99.07» '4411532»   '65.61» '4393421»   '1.00»  '65.59» '120.32»'123»   '4338263»   '0.98»  '65.43» '121.72»'123»   '570»   '186»   '52»'18111» '316 |
| 66 |  | '2024-09-13»'5870-03»   'lib966»'5870-03\_lib966»'145985»'143023»'97.97» '4411532»   '65.61» '4221519»   '0.96»  '65.39» '6.46»  '6» '839624»'0.19»  '65.81» '11.69» '11»'409»   '38»'33»'190013»'227 |  | 66 |  | '2024-09-17»'5870-03»   'lib966»'5870-03\_lib966»'145985»'143023»'97.97» '4411532»   '65.61» '4221519»   '0.96»  '65.39» '6.46»  '6» '839624»'0.19»  '65.81» '11.69» '11»'409»   '38»'33»'190013»'227 |  | 66 |  | '2024-09-18»'5870-03»   'lib966»'5870-03\_lib966»'145985»'143023»'97.97» '4411532»   '65.61» '4221519»   '0.96»  '65.39» '6.46»  '6» '839624»'0.19»  '65.81» '11.69» '11»'409»   '38»'33»'190013»'227 |
| 67 |  | '2024-09-13»'5872-03»   'lib900»'5872-03\_lib900»'587650»'582451»'99.12» '4411532»   '65.61» '4369601»   '0.99»  '65.51» '29.95» '30»'4171137»   '0.95»  '65.19» '30.97» '30»'909»   '195»   '153»   '41931» '470 |  | 67 |  | '2024-09-17»'5872-03»   'lib900»'5872-03\_lib900»'587650»'582451»'99.12» '4411532»   '65.61» '4369601»   '0.99»  '65.51» '29.95» '30»'4171137»   '0.95»  '65.19» '30.97» '30»'909»   '195»   '153»   '41931» '470 |  | 67 |  | '2024-09-18»'5872-03»   'lib900»'5872-03\_lib900»'587650»'582451»'99.12» '4411532»   '65.61» '4369601»   '0.99»  '65.51» '29.95» '30»'4171137»   '0.95»  '65.19» '30.97» '30»'909»   '195»   '153»   '41931» '470 |
| 68 |  | '2024-09-13»'6429-03»   'lib923»'6429-03\_lib923»'497218»'494253»'99.40» '4411532»   '65.61» '4369952»   '0.99»  '65.51» '25.92» '26»'4143267»   '0.94»  '65.17» '26.92» '26»'893»   '211»   '128»   '41580» '455 |  | 68 |  | '2024-09-17»'6429-03»   'lib923»'6429-03\_lib923»'497218»'494253»'99.40» '4411532»   '65.61» '4369952»   '0.99»  '65.51» '25.92» '26»'4143267»   '0.94»  '65.17» '26.92» '26»'893»   '211»   '128»   '41580» '455 |  | 68 |  | '2024-09-18»'6429-03»   'lib923»'6429-03\_lib923»'497218»'494253»'99.40» '4411532»   '65.61» '4369952»   '0.99»  '65.51» '25.92» '26»'4143267»   '0.94»  '65.17» '26.92» '26»'893»   '211»   '128»   '41580» '455 |
| 69 |  | '2024-09-13»'6435-03»   'lib955»'6435-03\_lib955»'1309534»   '1297407»   '99.07» '4411532»   '65.61» '4349238»   '0.99»  '65.54» '66.91» '68»'4256678»   '0.96»  '65.33» '68.17» '68»'2213»  '348»   '198»   '62294» '1196 |  | 69 |  | '2024-09-17»'6435-03»   'lib955»'6435-03\_lib955»'1309534»   '1297407»   '99.07» '4411532»   '65.61» '4349238»   '0.99»  '65.54» '66.91» '68»'4256678»   '0.96»  '65.33» '68.17» '68»'2213»  '348»   '198»   '62294» '1196 |  | 69 |  | '2024-09-18»'6435-03»   'lib955»'6435-03\_lib955»'1309534»   '1297407»   '99.07» '4411532»   '65.61» '4349238»   '0.99»  '65.54» '66.91» '68»'4256678»   '0.96»  '65.33» '68.17» '68»'2213»  '348»   '198»   '62294» '1196 |
| 70 |  | '2024-09-13»'6463-04»   'lib910»'6463-04\_lib910»'1643621»   '1628111»   '99.06» '4411532»   '65.61» '4390228»   '1.00»  '65.57» '78.53» '80»'4315278»   '0.98»  '65.37» '79.74» '80»'542»   '217»   '75»'21304» '300 |  | 70 |  | '2024-09-17»'6463-04»   'lib910»'6463-04\_lib910»'1643621»   '1628111»   '99.06» '4411532»   '65.61» '4390228»   '1.00»  '65.57» '78.53» '80»'4315278»   '0.98»  '65.37» '79.74» '80»'542»   '217»   '75»'21304» '300 |  | 70 |  | '2024-09-18»'6463-04»   'lib910»'6463-04\_lib910»'1643621»   '1628111»   '99.06» '4411532»   '65.61» '4390228»   '1.00»  '65.57» '78.53» '80»'4315278»   '0.98»  '65.37» '79.74» '80»'542»   '217»   '75»'21304» '300 |
| 71 |  | '2024-09-13»'6467-04»   'lib1450»   '6467-04\_lib1450»   '1953400»   '1939313»   '99.28» '4411532»   '65.61» '4384992»   '0.99»  '65.60» '116.63»'119»   '4345843»   '0.99»  '65.50» '117.62»'119»   '1000»  '495»   '214»   '26540» '512 |  | 71 |  | '2024-09-17»'6467-04»   'lib1450»   '6467-04\_lib1450»   '1953400»   '1939313»   '99.28» '4411532»   '65.61» '4384992»   '0.99»  '65.60» '116.63»'119»   '4345843»   '0.99»  '65.50» '117.62»'119»   '1000»  '495»   '214»   '26540» '512 |  | 71 |  | '2024-09-18»'6467-04»   'lib1450»   '6467-04\_lib1450»   '1953400»   '1939313»   '99.28» '4411532»   '65.61» '4384992»   '0.99»  '65.60» '116.63»'119»   '4345843»   '0.99»  '65.50» '117.62»'119»   '1000»  '495»   '214»   '26540» '512 |
| 72 |  | '2024-09-13»'6637-04»   'lib899»'6637-04\_lib899»'591853»'582965»'98.50» '4411532»   '65.61» '4350195»   '0.99»  '65.52» '28.84» '29»'4103536»   '0.93»  '65.24» '30.03» '30»'1999»  '310»   '270»   '61337» '1094 |  | 72 |  | '2024-09-17»'6637-04»   'lib899»'6637-04\_lib899»'591853»'582965»'98.50» '4411532»   '65.61» '4350195»   '0.99»  '65.52» '28.84» '29»'4103536»   '0.93»  '65.24» '30.03» '30»'1999»  '310»   '270»   '61337» '1094 |  | 72 |  | '2024-09-18»'6637-04»   'lib899»'6637-04\_lib899»'591853»'582965»'98.50» '4411532»   '65.61» '4350195»   '0.99»  '65.52» '28.84» '29»'4103536»   '0.93»  '65.24» '30.03» '30»'1999»  '310»   '270»   '61337» '1094 |
| 73 |  | '2024-09-13»'6639-04»   'lib965»'6639-04\_lib965»'2836411»   '2807979»   '99.00» '4411532»   '65.61» '4351974»   '0.99»  '65.59» '137.89»'141»   '4295280»   '0.97»  '65.45» '139.56»'141»   '862»   '309»   '111»   '59558» '451 |  | 73 |  | '2024-09-17»'6639-04»   'lib965»'6639-04\_lib965»'2836411»   '2807979»   '99.00» '4411532»   '65.61» '4351974»   '0.99»  '65.59» '137.89»'141»   '4295280»   '0.97»  '65.45» '139.56»'141»   '862»   '309»   '111»   '59558» '451 |  | 73 |  | '2024-09-18»'6639-04»   'lib965»'6639-04\_lib965»'2836411»   '2807979»   '99.00» '4411532»   '65.61» '4351974»   '0.99»  '65.59» '137.89»'141»   '4295280»   '0.97»  '65.45» '139.56»'141»   '862»   '309»   '111»   '59558» '451 |
| 74 |  | '2024-09-13»'6640-04»   'lib929»'6640-04\_lib929»'1396019»   '1371423»   '98.24» '4411532»   '65.61» '4363875»   '0.99»  '65.58» '63.75» '65»'4263229»   '0.97»  '65.38» '65.01» '65»'920»   '359»   '118»   '47657» '460 |  | 74 |  | '2024-09-17»'6640-04»   'lib929»'6640-04\_lib929»'1396019»   '1371423»   '98.24» '4411532»   '65.61» '4363875»   '0.99»  '65.58» '63.75» '65»'4263229»   '0.97»  '65.38» '65.01» '65»'920»   '359»   '118»   '47657» '460 |  | 74 |  | '2024-09-18»'6640-04»   'lib929»'6640-04\_lib929»'1396019»   '1371423»   '98.24» '4411532»   '65.61» '4363875»   '0.99»  '65.58» '63.75» '65»'4263229»   '0.97»  '65.38» '65.01» '65»'920»   '359»   '118»   '47657» '460 |
| 75 |  | '2024-09-13»'6769-04»   'lib962»'6769-04\_lib962»'1418690»   '1403197»   '98.91» '4411532»   '65.61» '4345903»   '0.99»  '65.55» '70.22» '71»'4265464»   '0.97»  '65.35» '71.36» '71»'2227»  '501»   '227»   '65629» '1216 |  | 75 |  | '2024-09-17»'6769-04»   'lib962»'6769-04\_lib962»'1418690»   '1403197»   '98.91» '4411532»   '65.61» '4345903»   '0.99»  '65.55» '70.22» '71»'4265464»   '0.97»  '65.35» '71.36» '71»'2227»  '501»   '227»   '65629» '1216 |  | 75 |  | '2024-09-18»'6769-04»   'lib962»'6769-04\_lib962»'1418690»   '1403197»   '98.91» '4411532»   '65.61» '4345903»   '0.99»  '65.55» '70.22» '71»'4265464»   '0.97»  '65.35» '71.36» '71»'2227»  '501»   '227»   '65629» '1216 |
| 76 |  | '2024-09-13»'6771-04»   'lib1462»   '6771-04\_lib1462»   '1761368»   '1751658»   '99.45» '4411532»   '65.61» '4389469»   '0.99»  '65.58» '104.23»'105»   '4349295»   '0.99»  '65.48» '105.13»'105»   '871»   '364»   '146»   '22063» '451 |  | 76 |  | '2024-09-17»'6771-04»   'lib1462»   '6771-04\_lib1462»   '1761368»   '1751658»   '99.45» '4411532»   '65.61» '4389469»   '0.99»  '65.58» '104.23»'105»   '4349295»   '0.99»  '65.48» '105.13»'105»   '871»   '364»   '146»   '22063» '451 |  | 76 |  | '2024-09-18»'6771-04»   'lib1462»   '6771-04\_lib1462»   '1761368»   '1751658»   '99.45» '4411532»   '65.61» '4389469»   '0.99»  '65.58» '104.23»'105»   '4349295»   '0.99»  '65.48» '105.13»'105»   '871»   '364»   '146»   '22063» '451 |
| 77 |  | '2024-09-13»'6775-04»   'lib963»'6775-04\_lib963»'1736814»   '1717255»   '98.87» '4411532»   '65.61» '4352358»   '0.99»  '65.55» '85.65» '87»'4272118»   '0.97»  '65.36» '87.06» '88»'2236»  '492»   '285»   '59174» '1211 |  | 77 |  | '2024-09-17»'6775-04»   'lib963»'6775-04\_lib963»'1736814»   '1717255»   '98.87» '4411532»   '65.61» '4352358»   '0.99»  '65.55» '85.65» '87»'4272118»   '0.97»  '65.36» '87.06» '88»'2236»  '492»   '285»   '59174» '1211 |  | 77 |  | '2024-09-18»'6775-04»   'lib963»'6775-04\_lib963»'1736814»   '1717255»   '98.87» '4411532»   '65.61» '4352358»   '0.99»  '65.55» '85.65» '87»'4272118»   '0.97»  '65.36» '87.06» '88»'2236»  '492»   '285»   '59174» '1211 |
| 78 |  | '2024-09-13»'6892-04»   'lib964»'6892-04\_lib964»'2315282»   '2290088»   '98.91» '4411532»   '65.61» '4359426»   '0.99»  '65.57» '110.31»'112»   '4307668»   '0.98»  '65.43» '111.50»'112»   '2275»  '508»   '276»   '52106» '1239 |  | 78 |  | '2024-09-17»'6892-04»   'lib964»'6892-04\_lib964»'2315282»   '2290088»   '98.91» '4411532»   '65.61» '4359426»   '0.99»  '65.57» '110.31»'112»   '4307668»   '0.98»  '65.43» '111.50»'112»   '2275»  '508»   '276»   '52106» '1239 |  | 78 |  | '2024-09-18»'6892-04»   'lib964»'6892-04\_lib964»'2315282»   '2290088»   '98.91» '4411532»   '65.61» '4359426»   '0.99»  '65.57» '110.31»'112»   '4307668»   '0.98»  '65.43» '111.50»'112»   '2275»  '508»   '276»   '52106» '1239 |
| 79 |  | '2024-09-13»'6895-04»   'lib1459»   '6895-04\_lib1459»   '2410848»   '2393301»   '99.27» '4411532»   '65.61» '4374284»   '0.99»  '65.57» '140.61»'142»   '4334739»   '0.98»  '65.49» '141.82»'142»   '2225»  '460»   '361»   '37248» '1182 |  | 79 |  | '2024-09-17»'6895-04»   'lib1459»   '6895-04\_lib1459»   '2410848»   '2393301»   '99.27» '4411532»   '65.61» '4374284»   '0.99»  '65.57» '140.61»'142»   '4334739»   '0.98»  '65.49» '141.82»'142»   '2225»  '460»   '361»   '37248» '1182 |  | 79 |  | '2024-09-18»'6895-04»   'lib1459»   '6895-04\_lib1459»   '2410848»   '2393301»   '99.27» '4411532»   '65.61» '4374284»   '0.99»  '65.57» '140.61»'142»   '4334739»   '0.98»  '65.49» '141.82»'142»   '2225»  '460»   '361»   '37248» '1182 |
| 80 |  | '2024-09-13»'6897-04»   'lib954»'6897-04\_lib954»'1912972»   '1894485»   '99.03» '4411532»   '65.61» '4377117»   '0.99»  '65.55» '97.33» '99»'4304764»   '0.98»  '65.37» '98.78» '100»   '2193»  '617»   '255»   '34415» '1181 |  | 80 |  | '2024-09-17»'6897-04»   'lib954»'6897-04\_lib954»'1912972»   '1894485»   '99.03» '4411532»   '65.61» '4377117»   '0.99»  '65.55» '97.33» '99»'4304764»   '0.98»  '65.37» '98.78» '100»   '2193»  '617»   '255»   '34415» '1181 |  | 80 |  | '2024-09-18»'6897-04»   'lib954»'6897-04\_lib954»'1912972»   '1894485»   '99.03» '4411532»   '65.61» '4377117»   '0.99»  '65.55» '97.33» '99»'4304764»   '0.98»  '65.37» '98.78» '100»   '2193»  '617»   '255»   '34415» '1181 |
| 81 |  | '2024-09-13»'7000-03»   'lib913»'7000-03\_lib913»'650076»'647008»'99.53» '4411532»   '65.61» '4331980»   '0.98»  '65.54» '33.77» '34»'4177236»   '0.95»  '65.27» '34.73» '34»'799»   '141»   '111»   '79552» '439 |  | 81 |  | '2024-09-17»'7000-03»   'lib913»'7000-03\_lib913»'650076»'647008»'99.53» '4411532»   '65.61» '4331980»   '0.98»  '65.54» '33.77» '34»'4177236»   '0.95»  '65.27» '34.73» '34»'799»   '141»   '111»   '79552» '439 |  | 81 |  | '2024-09-18»'7000-03»   'lib913»'7000-03\_lib913»'650076»'647008»'99.53» '4411532»   '65.61» '4331980»   '0.98»  '65.54» '33.77» '34»'4177236»   '0.95»  '65.27» '34.73» '34»'799»   '141»   '111»   '79552» '439 |
| 82 |  | '2024-09-13»'7135-04»   'lib987»'7135-04\_lib987»'1419752»   '1407892»   '99.16» '4411532»   '65.61» '4365589»   '0.99»  '65.57» '70.82» '72»'4278072»   '0.97»  '65.36» '72.08» '72»'1390»  '443»   '257»   '45943» '733 |  | 82 |  | '2024-09-17»'7135-04»   'lib987»'7135-04\_lib987»'1419752»   '1407892»   '99.16» '4411532»   '65.61» '4365589»   '0.99»  '65.57» '70.82» '72»'4278072»   '0.97»  '65.36» '72.08» '72»'1390»  '443»   '257»   '45943» '733 |  | 82 |  | '2024-09-18»'7135-04»   'lib987»'7135-04\_lib987»'1419752»   '1407892»   '99.16» '4411532»   '65.61» '4365589»   '0.99»  '65.57» '70.82» '72»'4278072»   '0.97»  '65.36» '72.08» '72»'1390»  '443»   '257»   '45943» '733 |
| 83 |  | '2024-09-13»'7514-04»   'lib1460»   '7514-04\_lib1460»   '1628304»   '1619630»   '99.47» '4411532»   '65.61» '4355320»   '0.99»  '65.57» '96.29» '97»'4312318»   '0.98»  '65.47» '97.18» '97»'974»   '339»   '127»   '56212» '488 |  | 83 |  | '2024-09-17»'7514-04»   'lib1460»   '7514-04\_lib1460»   '1628304»   '1619630»   '99.47» '4411532»   '65.61» '4355320»   '0.99»  '65.57» '96.29» '97»'4312318»   '0.98»  '65.47» '97.18» '97»'974»   '339»   '127»   '56212» '488 |  | 83 |  | '2024-09-18»'7514-04»   'lib1460»   '7514-04\_lib1460»   '1628304»   '1619630»   '99.47» '4411532»   '65.61» '4355320»   '0.99»  '65.57» '96.29» '97»'4312318»   '0.98»  '65.47» '97.18» '97»'974»   '339»   '127»   '56212» '488 |
| 84 |  | '2024-09-13»'7516-04»   'lib920»'7516-04\_lib920»'472211»'469870»'99.50» '4411532»   '65.61» '4335125»   '0.98»  '65.51» '24.70» '24»'4116092»   '0.93»  '65.18» '25.63» '25»'826»   '174»   '143»   '76407» '439 |  | 84 |  | '2024-09-17»'7516-04»   'lib920»'7516-04\_lib920»'472211»'469870»'99.50» '4411532»   '65.61» '4335125»   '0.98»  '65.51» '24.70» '24»'4116092»   '0.93»  '65.18» '25.63» '25»'826»   '174»   '143»   '76407» '439 |  | 84 |  | '2024-09-18»'7516-04»   'lib920»'7516-04\_lib920»'472211»'469870»'99.50» '4411532»   '65.61» '4335125»   '0.98»  '65.51» '24.70» '24»'4116092»   '0.93»  '65.18» '25.63» '25»'826»   '174»   '143»   '76407» '439 |
| 85 |  | '2024-09-13»'7517-04»   'lib930»'7517-04\_lib930»'1817740»   '1802631»   '99.17» '4411532»   '65.61» '4387467»   '0.99»  '65.57» '90.27» '93»'4312168»   '0.98»  '65.39» '91.66» '93»'989»   '343»   '156»   '24065» '500 |  | 85 |  | '2024-09-17»'7517-04»   'lib930»'7517-04\_lib930»'1817740»   '1802631»   '99.17» '4411532»   '65.61» '4387467»   '0.99»  '65.57» '90.27» '93»'4312168»   '0.98»  '65.39» '91.66» '93»'989»   '343»   '156»   '24065» '500 |  | 85 |  | '2024-09-18»'7517-04»   'lib930»'7517-04\_lib930»'1817740»   '1802631»   '99.17» '4411532»   '65.61» '4387467»   '0.99»  '65.57» '90.27» '93»'4312168»   '0.98»  '65.39» '91.66» '93»'989»   '343»   '156»   '24065» '500 |
| 86 |  | '2024-09-13»'7520-04»   'lib1461»   '7520-04\_lib1461»   '1798400»   '1789360»   '99.50» '4411532»   '65.61» '4382640»   '0.99»  '65.58» '105.84»'107»   '4339733»   '0.98»  '65.48» '106.81»'107»   '948»   '514»   '175»   '28892» '507 |  | 86 |  | '2024-09-17»'7520-04»   'lib1461»   '7520-04\_lib1461»   '1798400»   '1789360»   '99.50» '4411532»   '65.61» '4382640»   '0.99»  '65.58» '105.84»'107»   '4339733»   '0.98»  '65.48» '106.81»'107»   '948»   '514»   '175»   '28892» '507 |  | 86 |  | '2024-09-18»'7520-04»   'lib1461»   '7520-04\_lib1461»   '1798400»   '1789360»   '99.50» '4411532»   '65.61» '4382640»   '0.99»  '65.58» '105.84»'107»   '4339733»   '0.98»  '65.48» '106.81»'107»   '948»   '514»   '175»   '28892» '507 |
| 87 |  | '2024-09-13»'7538-03»   'lib948»'7538-03\_lib948»'1154863»   '1142670»   '98.94» '4411532»   '65.61» '4383972»   '0.99»  '65.56» '57.16» '58»'4283969»   '0.97»  '65.35» '58.27» '58»'980»   '320»   '135»   '27560» '496 |  | 87 |  | '2024-09-17»'7538-03»   'lib948»'7538-03\_lib948»'1154863»   '1142670»   '98.94» '4411532»   '65.61» '4383972»   '0.99»  '65.56» '57.16» '58»'4283969»   '0.97»  '65.35» '58.27» '58»'980»   '320»   '135»   '27560» '496 |  | 87 |  | '2024-09-18»'7538-03»   'lib948»'7538-03\_lib948»'1154863»   '1142670»   '98.94» '4411532»   '65.61» '4383972»   '0.99»  '65.56» '57.16» '58»'4283969»   '0.97»  '65.35» '58.27» '58»'980»   '320»   '135»   '27560» '496 |
| 88 |  | '2024-09-13»'8082-03»   'lib932»'8082-03\_lib932»'1903286»   '1892667»   '99.44» '4411532»   '65.61» '4367741»   '0.99»  '65.56» '93.23» '95»'4299548»   '0.97»  '65.38» '94.57» '95»'977»   '262»   '148»   '43791» '495 |  | 88 |  | '2024-09-17»'8082-03»   'lib932»'8082-03\_lib932»'1903286»   '1892667»   '99.44» '4411532»   '65.61» '4367741»   '0.99»  '65.56» '93.23» '95»'4299548»   '0.97»  '65.38» '94.57» '95»'977»   '262»   '148»   '43791» '495 |  | 88 |  | '2024-09-18»'8082-03»   'lib932»'8082-03\_lib932»'1903286»   '1892667»   '99.44» '4411532»   '65.61» '4367741»   '0.99»  '65.56» '93.23» '95»'4299548»   '0.97»  '65.38» '94.57» '95»'977»   '262»   '148»   '43791» '495 |
| 89 |  | '2024-09-13»'8864-03»   'lib967»'8864-03\_lib967»'1350989»   '1335457»   '98.85» '4411532»   '65.61» '4341922»   '0.98»  '65.54» '66.42» '67»'4256802»   '0.96»  '65.35» '67.56» '68»'2254»  '452»   '265»   '69610» '1228 |  | 89 |  | '2024-09-17»'8864-03»   'lib967»'8864-03\_lib967»'1350989»   '1335457»   '98.85» '4411532»   '65.61» '4341922»   '0.98»  '65.54» '66.42» '67»'4256802»   '0.96»  '65.35» '67.56» '68»'2254»  '452»   '265»   '69610» '1228 |  | 89 |  | '2024-09-18»'8864-03»   'lib967»'8864-03\_lib967»'1350989»   '1335457»   '98.85» '4411532»   '65.61» '4341922»   '0.98»  '65.54» '66.42» '67»'4256802»   '0.96»  '65.35» '67.56» '68»'2254»  '452»   '265»   '69610» '1228 |
| 90 |  | '2024-09-13»'8867-03»   'lib968»'8867-03\_lib968»'1547576»   '1532489»   '99.03» '4411532»   '65.61» '4344450»   '0.98»  '65.55» '77.48» '79»'4268677»   '0.97»  '65.36» '78.68» '79»'2229»  '543»   '253»   '67082» '1212 |  | 90 |  | '2024-09-17»'8867-03»   'lib968»'8867-03\_lib968»'1547576»   '1532489»   '99.03» '4411532»   '65.61» '4344450»   '0.98»  '65.55» '77.48» '79»'4268677»   '0.97»  '65.36» '78.68» '79»'2229»  '543»   '253»   '67082» '1212 |  | 90 |  | '2024-09-18»'8867-03»   'lib968»'8867-03\_lib968»'1547576»   '1532489»   '99.03» '4411532»   '65.61» '4344450»   '0.98»  '65.55» '77.48» '79»'4268677»   '0.97»  '65.36» '78.68» '79»'2229»  '543»   '253»   '67082» '1212 |
| 91 |  | '2024-09-13»'8868-03»   'lib950»'8868-03\_lib950»'1257056»   '1244447»   '99.00» '4411532»   '65.61» '4372399»   '0.99»  '65.54» '61.80» '62»'4281178»   '0.97»  '65.34» '62.91» '63»'2161»  '510»   '241»   '39133» '1154 |  | 91 |  | '2024-09-17»'8868-03»   'lib950»'8868-03\_lib950»'1257056»   '1244447»   '99.00» '4411532»   '65.61» '4372399»   '0.99»  '65.54» '61.80» '62»'4281178»   '0.97»  '65.34» '62.91» '63»'2161»  '510»   '241»   '39133» '1154 |  | 91 |  | '2024-09-18»'8868-03»   'lib950»'8868-03\_lib950»'1257056»   '1244447»   '99.00» '4411532»   '65.61» '4372399»   '0.99»  '65.54» '61.80» '62»'4281178»   '0.97»  '65.34» '62.91» '63»'2161»  '510»   '241»   '39133» '1154 |

Araxis Merge (but not the data content of this report) is Copyright © 1993–2024 Araxis Ltd (www.araxis.com). All rights reserved.
