## Supplementary material for "MTBseq-nf: Enabling Scalable Tuberculosis Genomics “Big Data” Analysis through a User-Friendly Nextflow Wrapper for MTBseq pipeline": SD-7 Growth of total execution time of different modes of MTBseq-nf for 5 datasets with increasing cohort size.

**Growth of total execution time (runtime) of different modes of MTB-seq-nf for 5 datasets with increasing cohort size.**

| Number of samples | MTBseq-nf (default) | MTBseq-nf (parallel) |
|---|---|---|
| 5 | 0h53m0s | 0h34m0s |
| 10 | 2h2m0s | 1h0m45s |
| 20 | 4h32m0s | 1h53m0s |
| 40 | 8h57m0s | 4h43m0s |
| 80 | 18h32m0s | 7h14m0s |
