## Supplementary material for "MTBseq-nf: Enabling Scalable Tuberculosis Genomics “Big Data” Analysis through a User-Friendly Nextflow Wrapper for MTBseq pipeline": SD-8 Summary of different executions of MTBseq and MTBseq-nf in the triplicated set of experiments.

| Run ID | Run name | Description | Notable non-default parameter |
| --- | --- | --- | --- |
| mtbseq-standard-run1 | pub-90samples-mtbseq-standard-run1 | MTBseq pipeline | |
| mtbseq-standard-run2 | pub-90samples-mtbseq-standard-run2 | MTBseq pipeline | |
| mtbseq-standard-run3 | pub-90samples-mtbseq-standard-run3 | MTBseq pipeline | |
| mtbseq-nf-run1 | pub-90samples-mtbseq-nf-run1 | MTBseq-nf pipeline | |
| mtbseq-nf-run2 | pub-90samples-mtbseq-nf-run2 | MTBseq-nf pipeline | |
| mtbseq-nf-run3 | pub-90samples-mtbseq-nf-run3 | MTBseq-nf pipeline | |
| mtbseq-nf-parallel-run1 | pub-90samples-mtbseq-nf-parallel-run1 | MTBseq-nf pipeline | –parallel |
| mtbseq-nf-parallel-run2 | pub-90samples-mtbseq-nf-parallel-run2 | MTBseq-nf pipeline | –parallel |
| mtbseq-nf-parallel-run3 | pub-90samples-mtbseq-nf-parallel-run3 | MTBseq-nf pipeline | –parallel |
